# Diversity of Resilin incidence in the insect leg

**DOI:** 10.1101/2025.07.29.667210

**Authors:** Steven Lerch, Bernard Moussian

## Abstract

Insect movements rely on membranous elastic types of cuticle that bridge hard and tanned types, which serve as an exoskeleton. A prominent, polymerous protein of membranous cuticles is Resilin, known for its flexibility and elasticity. In the articulated legs, including the coxa, the trochanter, the femur, the tibia and the tarsi, Resilin incidences in or at the joints of these podomers have been identified in various species such as the fruit fly *Drosophila melanogaster* and the desert locust *Schistocerca grigaria*. A systematic, comparative work on Resilin identification in insect legs is missing.

Here, using a microscopic intensity subtraction method, we identified Resilin incidences in the legs of 29 species belonging to the major insect orders. This method relies on the classical identification of Resilin based on Dityrosine (DT) bonds between Resilin monomers, Pro-Resilin, that emit blue light when illuminated with UV light. By subtracting the background intensity counts per pixel (counts) of a common blue filter from the counts obtained by a specific filter for the DT spectrum in insects (microscopic intensity subtraction method, MISM), we both confirm the presence of known Resilin locations and identify new incidences in the podomers such as a coxa-trochanter signal. Together, we find that Resilin distribution in insect legs is highly diverse with no incidence common to all species tested. This underlines that the use of Resilin follows a rapid and optimizing evolutionary adaptation. Our data pave the path for studies of the comparative function of Resilin in insect legs.

## Introduction

For moving through their environment, insects are capable of flying, walking, jumping, swimming, digging, or crawling. The most important organs for locomotion are – together with the wings – the three pairs of thoracic legs. These have an organisation of alternating long and stiff cylinders interconnected by short and flexible rings (e.g. (Snodgrass 1935; Jabaud and Moussian 2024). The stiff cylinders, or podomeres, have received specific names. A leg starts with the coxa, which joins the thorax. This is followed by the usually short trochanter, which in many insects is quite firmly connected with the commonly long femur following it. Between the latter and the likewise long tibia the “knee” of the insect leg is located, and femur and tibia are the major force-mediating podomeres during locomotion. The tarsus, which is in contact with the substrate, terminates the leg; it is subdivided usually into five tarsomeres. The small terminal pretarsus consists of a small plate on the underside (unguitractor) and may carry specialised devices such as claws, an adhesive arolium, and terminal hairs (Figure 1). The opposing margins of neighbouring podomeres form discrete articulatory structures (such as ball-and-socket-joints), which bridge the flexible rings between them (e.g. Schmidt *et al*. 2009). Thereby, the relative movement between successive podomeres is highly coordinated. Leg movements are affected by muscles inside the leg that connect different (in most cases neighbouring) podomeres, or by muscles coming from the thoracic segments bearing the leg (e.g. Snodgrass (1935); Alsop (1978)). In many cases, the distal attachment of a muscle is placed on a long tendon, which is a tubular invagination of the cuticular cover of the body, often with thread-like to laminar internal parts. Since such a tendon offers much surface for the attachment of a strong muscle, this allows to exert much muscular force to a small area of the body surface and thus exact movement.

**figure 1.**
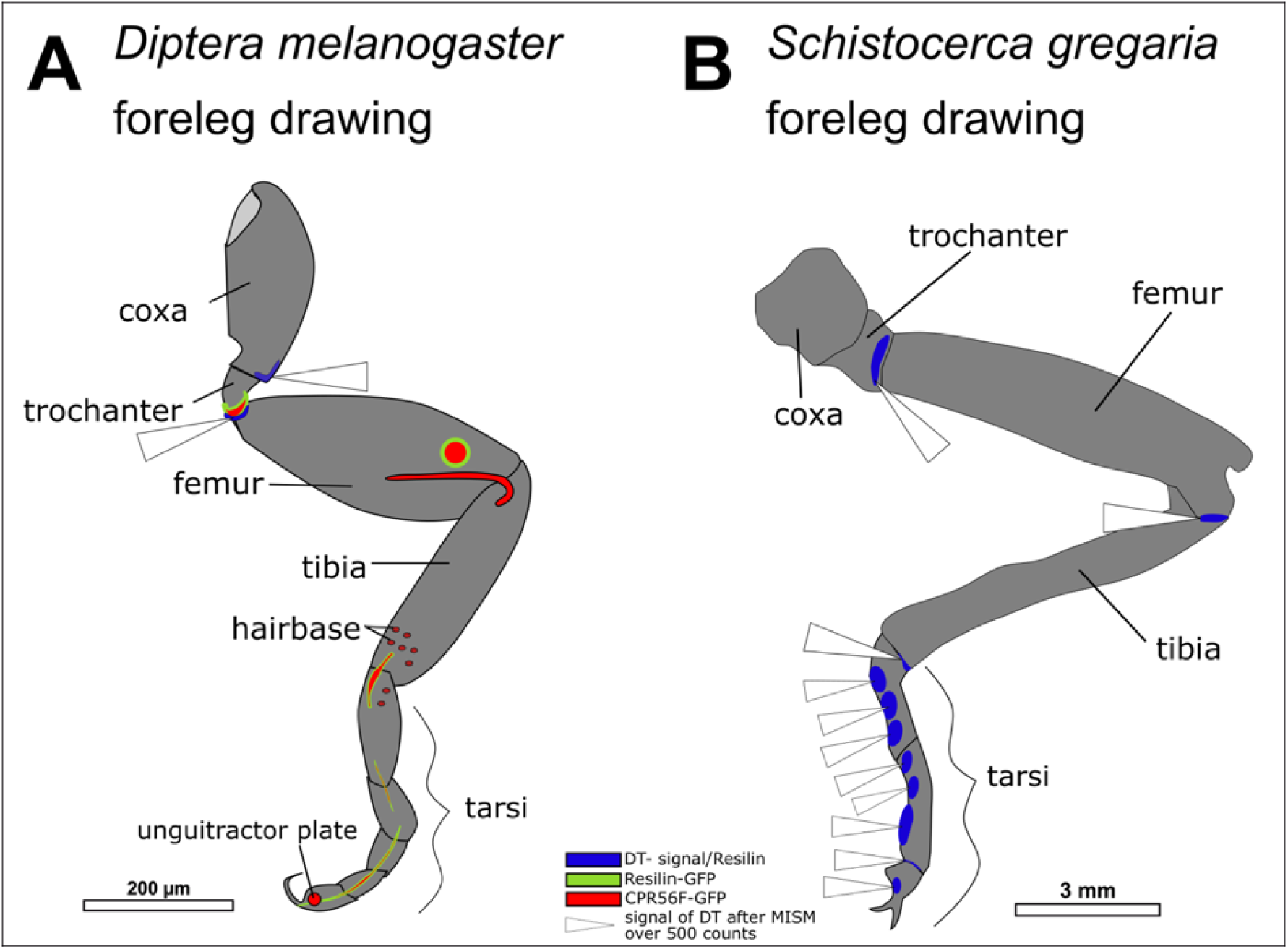
Schemes of typical insect legs. To exemplify the variety of legs and illustrate the general morphological structures of the podomeres, the forelegs of (A) *D. melanogaster* and (B) *S. gregaria* are drawn. In addition to the differences in podomere shape and size, the known localisation of Resilin-network proteins and DT-signals is given (see Lerch *et al*. 2020; 2022). In blue are areas of DT-signals as indication of Resilin; Resilin-GFP localisation is indicated in green, while CPR56F-RFP localisation is marked in red (see Lerch *et al*. 2020). The white arrowheads indicate locations where the signals fit the criteria of MISM.

Despite the conserved organization, there is considerable variation of leg morphology among different species and, to a varied extent, among the leg pairs of an individual. Obviously, strong differences relate to different roles of the legs in locomotion or even in other activities (e.g. prey catching with the fore legs in Mantodea). In, for instance, most Hemiptera, Diptera, Hymenoptera, and Coleoptera, all three pairs of legs are used for linear walking on a horizontal or vertical fundament and have a rather uniform morphology, although there are often differences in podomere length. Siphonaptera and Orthoptera, by contrast, jump using their hindlegs, which are usually longer and thicker than the fore- and midlegs. While the releasing force for the jump of locusts comes from the leg region between the femur and the tibia, fleas use the combination of the coxa and the trochanter to unleash the jump (Bennet-Clark & Lucey 1967; Rothschild & Schlein 1975; Heitler 1977; Sutton & Burrows 2010; Burrows 2012).

The mechanical properties of the legs depend gon the materials used for the cuticular exoskeleton. The long stiff cylinders consist of a hard type of cuticle (sclerotised), whereas the joints are made of a soft type of cuticle (membranous). Altogether, however, in insect cuticle a gradual range from very hard to very soft is represented; this is also true for the legs. In both types of cuticle, the mechanical properties largely depend on the proteins that interact with each other and with the chitin scaffold of the cuticle. In the soft and also elastic cuticle, one of the best studied proteins is Resilin, which was discovered and described by T. Weis-Fogh (Weis-Fogh 1960) and T. Weis-Fogh and S. O. Andersen (Andersen & Weis-Fogh 1964) in the locust. To date, it has been shown that Resilin is a matrix consisting of the Pro-Resilin protein molecules cross-linked via dityrosine (DT) bonds. Extensive repeats in the Pro-Resilin amino acid sequence (Elvin *et al*. 2005; Qin *et al*. 2009; Cheng et al. 2010; Qin et al. 2011; Qin et al. 2012) and the DT network between Pro-Resilin units, possibly involving additional proteins such as Cpr56F, (Cornman 2009; Andersen 2010; Lerch *et al*. 2020) are both responsible for visco-elastic properties of Resilin matrices (Andersen 1964; Elvin *et al*. 2005; Lyons et al. 2009; Cheng et al. 2010; Qin et al. 2011). In a number of cases, Resilin matrices have been visualised and identified via fluorescence microscopy based on the property of DT bonds to emit blue light (around 420 nm wavelength) when excited with UV-light (around 355 nm wavelength). However, a global view on the presence of DT i.e. Resilin matrix and leg morphological structures is missing to date.

The common tool for identifying DT in a number of published reports is the fluorescence microscope with a filter set up originally destined to detect the DNA dye DAPI (4′,6-Diamidino-2-phenylindole) using an excitation range around 350nm. This setup, however, detects a broad signal emission range outside 420nm (Haas *et al*. 2000 a; b; Neff *et al*. 2000; Burrows 2016). In a strict sense, this filter set up possibly allows detection of non-DT signals. Considerable background signal by conventional fluorescence microscopy may additionally complicate detailed analyses. Higher sensitivity is achieved by the confocal microscope, where a specific laser beam with a narrow emission filter is used. However, for large and thick samples (as legs of larger insects are) a routine application of confocal microscopy is not advised. To overcome this limited availability of the confocal microscope, a more specific UV-filter set up for fluorescence microscopes is needed. Here, we used such a specific filter set up (see methods) named DT filter (AF405) covering emission signals at and around 420nm on locust (*Schistocerca gregaria*) and fruit fly (*Drosophila melanogaster*) legs to verify previously reported DT areas. To improve the signal-to-background ratio, we removed the signal obtained by the DAPI filter setup from the signal obtained with the DT filter (Microscopic intensity subtraction method (MISM). With this set up, on a fluorescence microscope, we were able to recapitulate all DT locations in *D. melanogaster* that we had previously determined by a native Pro-Resilin-GFP reporter by confocal microscopy (Lerch *et al*. 2020). Additionally, we detected the half-moon-like DT formation which M. Burrows showed on the jumping leg between femur and tibia in *S. gregaria* (Burrows and Sutton 2012; Bayley, Sutton and Burrows 2012). This new microscopic set-up enabled us to systematically and exactly localise DT-rich areas (and thus likely Resilin) in the cuticle of any morphological structure needed for locomotion in a number of insect species representing various insect orders.

## Methods

### Insect species, rearing and storage

In this work, we studied the following insect species: the fruit fly *Drosophila melanogaster*, the cherry fly *D. suzukii* and *D. hydei* (Diptera: Drosophilidae), the locust *Schistocerca gregaria* (Orthoptera: Acrididae), the red flour beetle *Tribolium castaneum,* the mealworm beetle *Tenebrio molitor* (both Coleoptera: Tenebrionidae), the European flour moth *Ephestia kuehniella* (Lepidoptera: Pyralidae), the honey bee (drones) of *Apis mellifera* and the sun beetle *Pachnoda marginata* (Coleoptera: Scarabaeidae). All animals were kept at room temperature in species-appropriate vials or boxes and diet.

*Drosophila melanogaster* and *suzukii* populations were captured in Tübingen (Wang *et al*. 2020). A population of *Ephestia kuehniella* was started with four caterpillars captured in a flour bag obtained in Bad Wimpfen, Germany. *Tribolium castaneum was* obtained from the Kun Yan Zhu’s laboratory at the Kansas State University in Manhattan, USA. The honeybee drones were obtained from a farm near Pforzheim, Germany. The other insect species were purchased from the animal food supplier in Dresden (Germany) Fressnapf Holding SE. By feeding the larvae of *T. molitor* and *P. marginata* with wheat and vegetables adults were obtained. Several insect species were gathered from the field and used immediately for microscopy. 29 species were tested in this work. In addition, several unidentified individuals with a vague classification were analysed to extend the variety within the taxonomic groups. A complete list of the analysed insect samples including the number of individuals is provided in supplementary table 1.

### Preparation and mounting

For dissection, large insects were immobilized with ether or a short incubation at −20°C for 5 minutes and then killed by decapitation. Dissected body parts including legs were mounted in ROTI®Mount Aqua (Carl-Roth GmbH Germany) between a cover slip and a slide. Small insects were mounted directly in ROTI®Mount Aqua for microscopy. All mounted animals or animal parts were analysed latest 72 hours after preparation. The pH-value of the mounting medium was neither communicated nor determined.

### Microscopy and imaging

The light source was a halogen lamp. To visualize DT in biological materials (excitation at 355nm, emission at 420nm, figure 2), we used a specific filter set up with a narrower excitation field (**AF405**=400-440nm range of emitted light; AHF Analysentechink AG) than provided by the standard DAPI-filter set up. Following filters were present in the inverse Axio Observer Z1 (Zeiss) equipped with an Axiocam Mono camera: **AF405** (DT specific filter): Ex: 357/44 nm Brightline HC, splitter HC BS 389 nm, Em: 420/40 nm ET Bandpass; **DAPI**: Ex: 359/48 nm, splitter: 395 nm, Em: 445/50 nm; and **GFP**: Ex: 470/40 nm, splitter: 495 nm, Em: 525/50 nm. Depending on the size of the specimen the objectives and imaging times were adjusted. These objectives were used: EC Plan-Neofluar 5x/0.67 M27 DRY, EC Plan-Neofluar 10x/0.3 M27 DRY, and EC Plan-Neofluar 20x/0.5 M27 DRY.

**figure 2.**
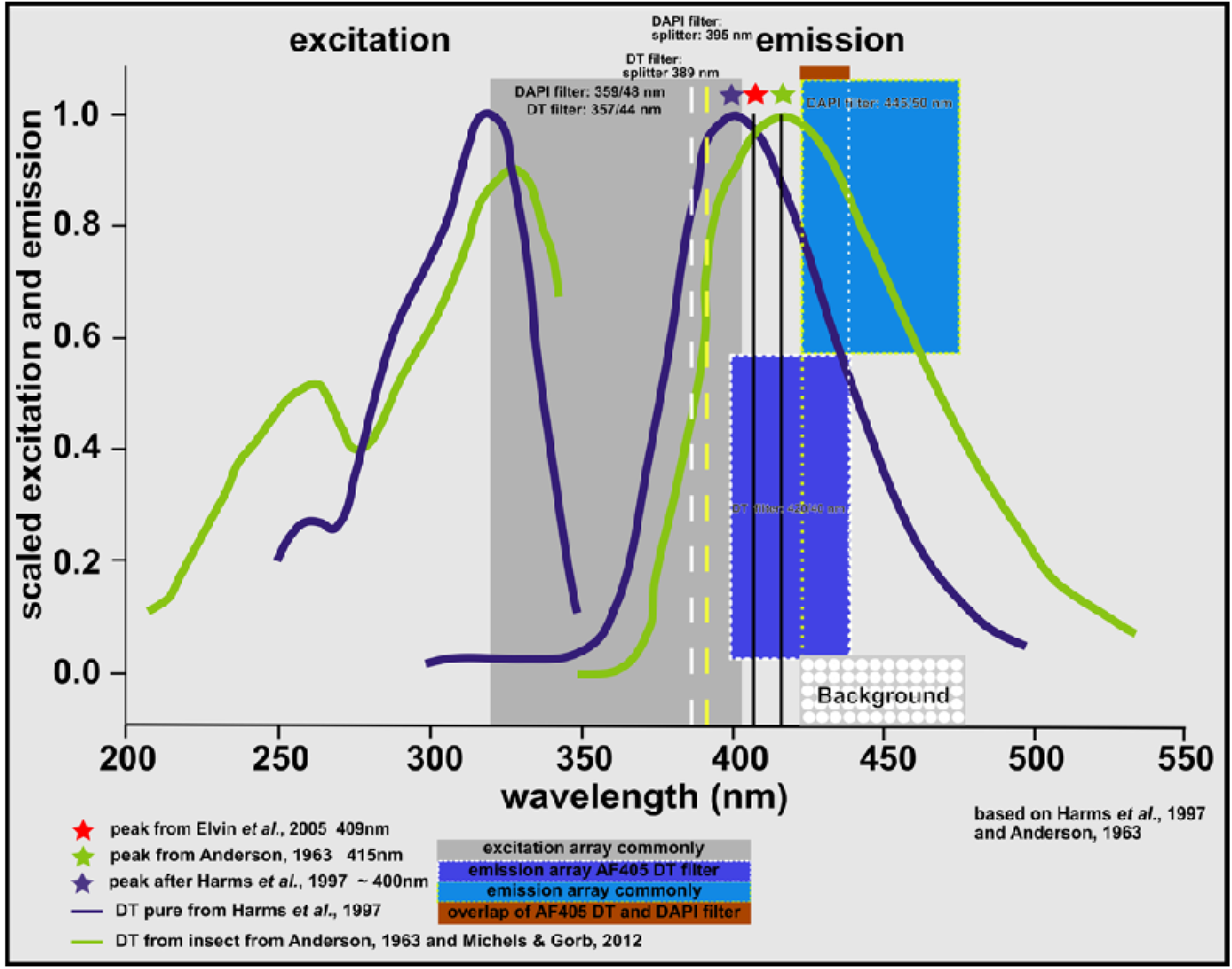
MISM filter-set ups for DT detection. In this graph, we present the commonly used filters, and the DT-filter set up used in this work. Also, we show the emission difference between pure DT auto-fluorescence versus the shift in biological composed material. (Blue line) The spectrum of pure DT monitored by Harms et. al, 1997 differs from the insect auto-fluorescence spectrum detected by Anderson (1963, orange line). The blue asterisk represents the peak at around 400nm determined by Harms *et al*. 1997, the red asterisk shows the peak at 409nm published by Elvin *et al*. (2005). The orange asterisk marks the peak at 415 nm reported by Anderson (1963); this corresponds to the peak published by Michels and Gorb (2012). Both filter sets use the same excitation range but differ immensely in the emission range. The standard DAPI filter covers a broad range right to the peak of insect specific DT, while our filter encompasses the peak in a narrow range. The splitters, avoiding light from the excitation, are close to each other. Hence, DAPI detection covers more background of lesser DT signal than the DT specific filter (see white doted grey box).

All images for a species are displayed with the intensities. Assembly of images of large specimens was done manually. Image analyses and processing were done on ZenBlue and „InkScape“ (www.inkscape.org).

### Microscopic intensity subtraction method (MISM)

Auto-fluorescence of biological material can be used to gain information on structures and by consequence their function; however, localisation and intensity quantification of auto-fluorescent signals for comparative approaches for example between individuals or species is difficult. The two main problems of auto-fluorescence are: 1) how pure is the signal? Does the auto-fluorescence emanate from one or more structural sources? 2) To what extent do other surrounding structural elements dim the auto-fluorescent signal? Both problems have an effect both on the acceptance of the presence of a true signal at low intensity and especially on the spatial extension of the structure of interest at normal and high intensity as often area borders are not sharp but graded.

These problems are evident in the insect cuticle. DT bonds are present in the insect cuticle and display an auto-fluorescence at maximum of 400nm when excited with UV light (e.g. 355nm) reflecting the presence of the dityrosinylated cuticle protein Pro-Resilin that marks visco-elastic and resilient cuticle regions. The intensity of DT as described in hundreds of publications is not uniform even within one species and of course between species (for example in Haas *et al*. 2000a; Voigt *et al*., 2007; Burrows and Sutton 2012; Michels and Gorb 2012; Ma *et al*. 2015; Reinhardt *et al*. 2015; Burrows 2016; Nadein and Betz 2016). Especially weak signals are difficult to discern.

Using the model insect *Drosophila melanogaster*, we define two references in order to be able to evaluate weak DT auto-fluorescence in the insect cuticle. One reference is the expression distribution of a GFP-tagged version of the dityrosinylated cuticle protein Pro-Resilin in the adult fly (Lerch *et al*. 2020). With little exception, we have previously shown that all DT signals overlap with the GFP signal of the GFP-Pro-Resilin source (Lerch *et al*. 2020). The second reference consists of two sets of DT signal intensity determinations (see below). This reference is especially necessary to accept or refute low signal intensities, at which the GFP signal is also ambiguous.

This second reference needs a detailed explanation. Again, it is conceived to allow evaluation of especially weak signals. Are these true or background signals? We know, for instance, that the outermost cuticle layer, the envelope displays auto-fluorescence when illuminated with UV light (Zuber et al. 2018). The emission lies outside the detection range of the DT specific filter (AF405) but is fully detected by the DAPI filter. During our work to recapitulate DT signals with a DT specific filter with a narrow wavelength detection range by fluorescence microscopy, we observed that the intensity of *bona fide* DT signals detected by the specific filter is generally higher than the one detected by the DAPI filter with a broad wavelength detection range that has been used in several published works (figure 3). This detection utilised the software Zenblue from Zeiss by simultaneously showing the intensities of all used filters in each pixel of the sample image (see figures 4 and 5). Based on this observation, to define the spatial distribution of DT bonds with a weak signal, we reckon that in the case of background signal the intensities of both signals should be similar. Thus, subtraction of the DAPI signal from the DT-specific signal should yield a positive signal in different insect species (see figures 4 and 5).

**figure 3.**
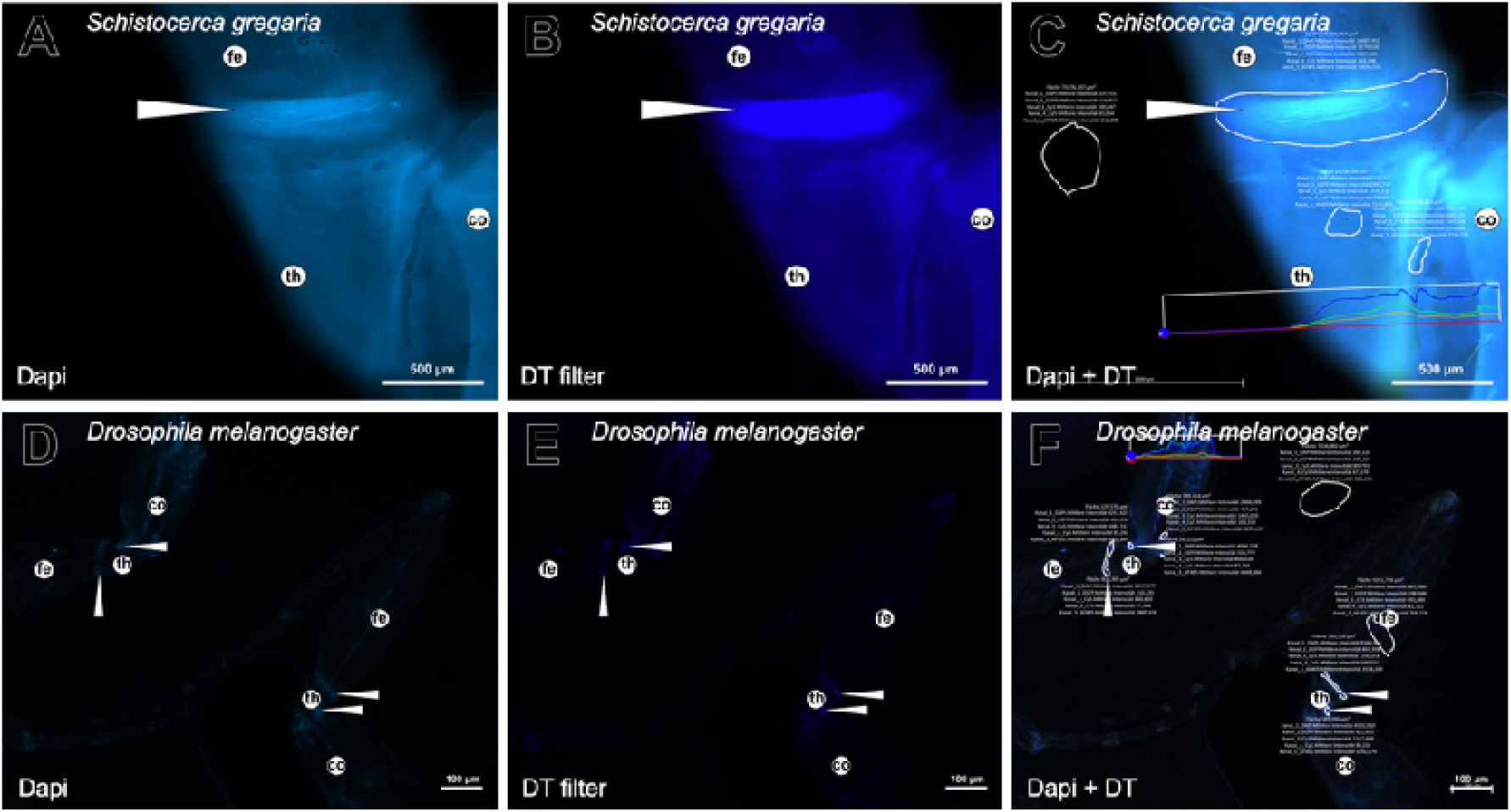
Comparison of signals detected with the DAPI or DT filter in *S. gregaria* and *D. melanogaster.* The problem of auto-fluorescence in biological samples is the overlapping of fluorescence from diverse molecular sources resulting in difficulties in interpretation of the signal of interest. To demonstrate the effect of our approach, here, we compare in detail the auto-fluorescence detected with the generally used (A, D) DAPI filter with the (B, E) auto-fluorescence obtained with the specific DT filter. For this purpose, showing the DT signal in the trochanter-femur junction of the desert locust and the coxa-trochanter signal in the fruit fly, images with all measurement information are presented (C, F). A strong auto-fluorescence is visible especially in the *S. greagaria* tissues outside the bona fide DT signal region. As these parts of the cuticle do not contain Resillin, we conclude that this signal represents a background situation detected with both filters. Consistently, the DT-DAPI signal intensity difference is below 500 counts in these regions, while in the Resilin regions these counts are high. Thus, signal subtraction improves identification and allows reducing false positive signals (more details on the measurement method see methods and supplementary discussion on the threshold and the exposure time). The white arrowheads point to the DT locations standing for Resilin. The images were obtained with the inverse Axio Observer Z1 (Zeiss) microscope and the Axiocam Mono camera. co… coxa, fe… femur, tb… tibia, th… trochanter, tr… tarsi The DAPI and the DT filters are named in German by the software: “Kanal_1_DAPI. Mittlere Intensität” and “Kanal_5_AF405.Mittlere Intensität”, respectively.

**figure 4.**
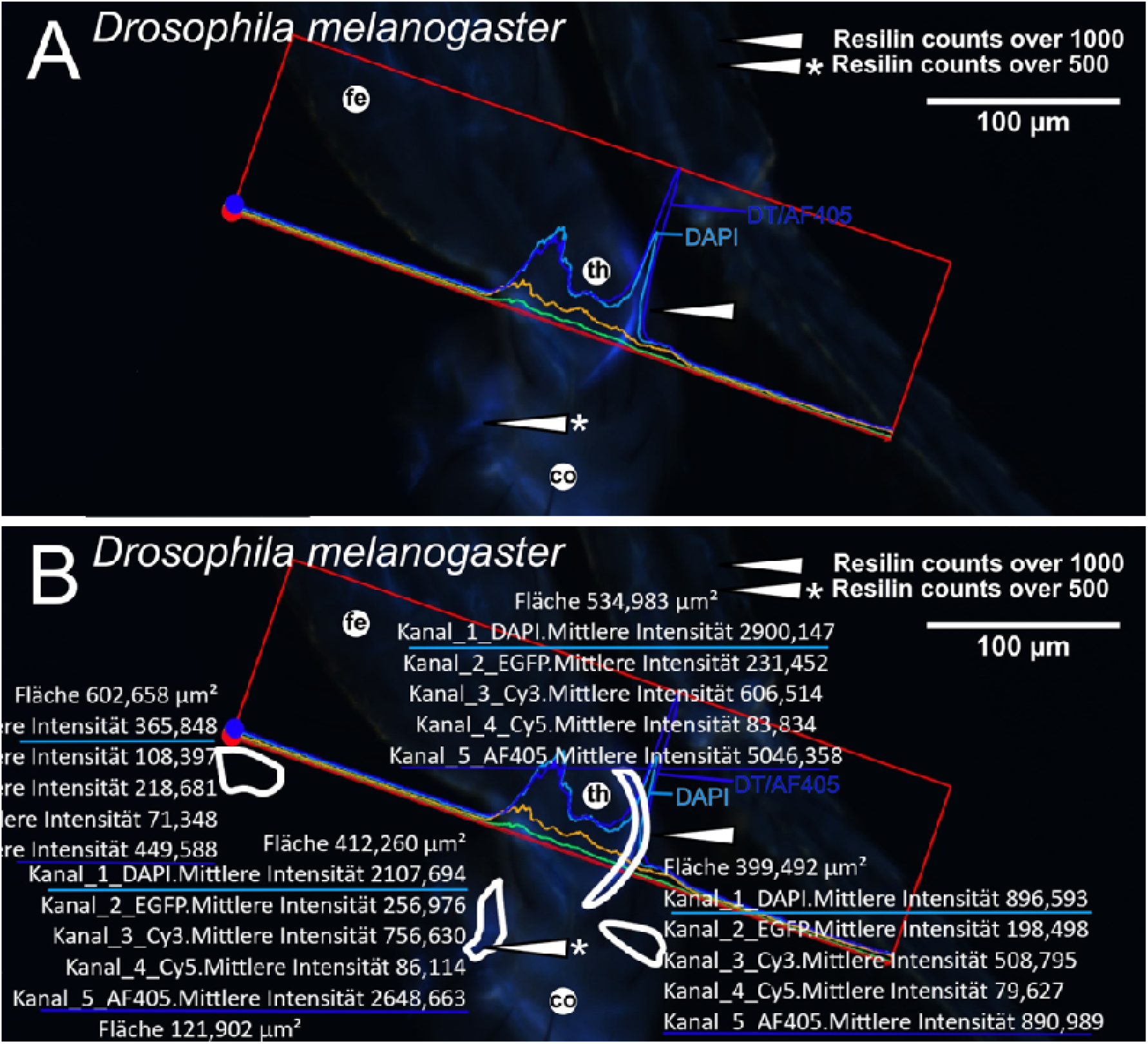
Measurement process on *D. melanogaster* samples. Images were analyzed using the software ZenBlue from Zeiss. Two consecutive measurements were conducted. First, the signal profiles along horizontal lines were determined for the whole image. Four different channels (DAPI, “EGFP”, “Cy3”, “Cy5”, AT405) were used; the values of the DAPI and DT filters are traced in two different blue colors, the brighter one for the DAPI filter and the darker one for the DT filter. Based on the scans, an area of high DT values was identified and manually drawn into the image (white line). The intensities for these areas were re-determined to assign DT incidence or to refute it according to the intensity of the DAPI signal intensity. (A) The trochanter-coxa of the fly leg is characterized by auto-fluorescence detected with both the DAPI and the DT filter. (B) In the same image, the source data of the measurement and the measured areas (confined by the white lines) have been added. The subtraction of the DT signal from the DAPI signal reveals four situations: areas with a weak signal above 500 counts, those with a strong signal above 1000 counts, those below 500 counts that we define as background and areas with no signal (black). The white arrowheads point to the DT locations standing for Resilin. The images were obtained with the inverse Axio Observer Z1 (Zeiss) microscope and the Axiocam Mono camera.co… coxa, fe… femur, tb… tibia, th… trochanter, tr… tarsi The DAPI and the DT filters are named in German by the software: “Kanal_1_DAPI. Mittlere Intensität” and “Kanal_5_AF405.Mittlere Intensität”, respectively.

**figure 5.**
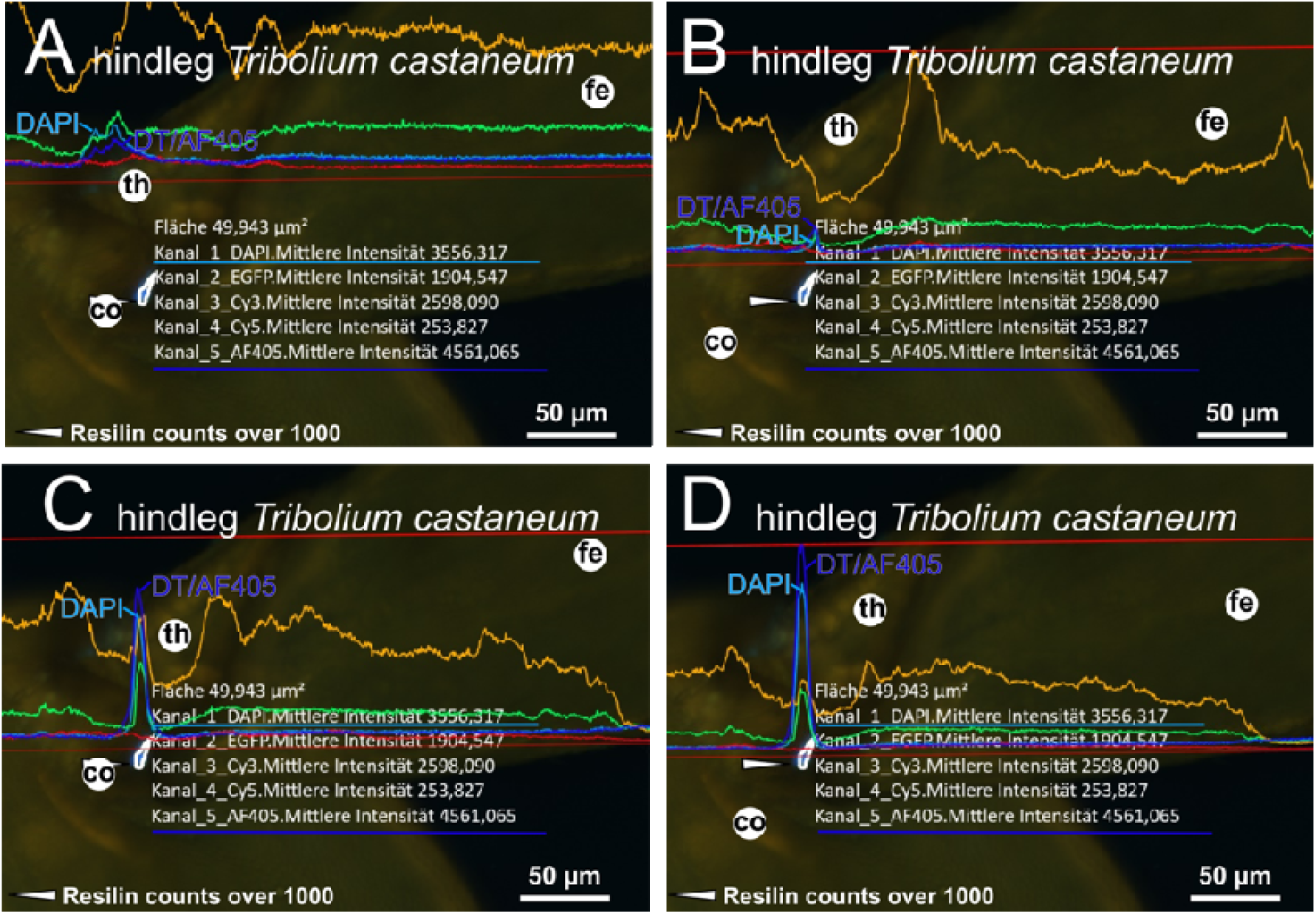
Measurement process on *T. castaneum* samples. The fluorescence signal profiles (generated with the software ZenBlue from Zeiss) in the beetle *T. castaneum* are, compared to *D. melanogaster*, more complex as high signal values are detected in all four channels. Signal quantification is done as for *D. melanogaster* (Figure 4), the resulting graphs for four lines are shown in A, B, C and D. In this case, the coxa-trochanter joint of the hindleg of *T. castaneum* has been analyzed. In brief, we first scanned line for line the whole leg structure in the image and identified areas without a visible difference between the DAPI and DT filters as represented by the graphs. Second, we identified pixel lines, where the graphs the DT signal intensity was higher than that of the DAPI to design a DT area with difference thresholds over 500 (weak) or 1000 (strong). The white arrowheads point to the DT locations standing for Resilin. The images were obtained with the inverse Axio Observer Z1 (Zeiss) microscope and the Axiocam Mono camera. co… coxa, fe… femur, tb… tibia, th… trochanter, tr… tarsi The DAPI and the DT filters are named in German by the software: “Kanal_1_DAPI. Mittlere Intensität” and “Kanal_5_AF405.Mittlere Intensität”, respectively.

To further refine the information on DT localisation, we defined two thresholds of acceptance after subtraction the intensity counts per pixel (counts) of each filter set up with a range between 0 and 16,000 calculated by the ZEN software. That is, together with the un-subtracted intensity, a low and a high threshold of intensity difference with the low intensity difference representing a low specificity and the high intensity difference representing a high specificity of the signal intensity were considered. This method respects also the problem of signal dimming differences in different species that would alter DT and/or DAPI signal intensities. Dimming includes all pathways of signal reduction like blocking, quenching or light absorption. Dimming is discussed in more detail in the supplementary data.

For standardisation, we applied this method on the wing hinge, the leg and the proboscis including the cibarium and the labellum of *D. melanogaster* (Lerch *et al*. 2020). In particular, we examined the DT distribution in three fruit fly lines including wild-type, the *pro-resilin* knock-down (*hdw*) flies and flies expressing a GFP-tagged Pro-Resilin version to localise DT areas defined by Resilin.

First, we verified the co-location of DT and Pro-Resilin-GFP. Second, we compared the intensities of signals detected by both filters in wild-type and *hdw* flies that we have shown to have reduced amounts of DT. Applying both thresholds, with one exception, the differences between signal intensities of DT areas in wild-type and *hdw* flies became more evident. The exception is the cibarium that with does not show significant difference between the intensity values obtained with or without subtraction in wild-type and *hdw* flies. We conclude that subtraction enhances the genuine DT signal.

As a next step, we sought to transfer our method developed in *D. melanogaster* to *D. suzukii* in order to check its robustness. As expected, we encounter a problem in cases where the signal intensity is low *per se*. While in the leg and in the proboscis, the situation with respective single strong signals is clear in both species, in the wing hinge where at least five sources of signal with different and low intensities have been identified in *D. melanogaster* (Lerch *et al*. 2020; 2022), the situation is, by contrast, difficult.

Besides the five DT spots in the wing hinge identified by confocal microscopy, here, by fluorescence microscopy, we found two additional DT spots (the sixth and the seventh), one of which does not overlap with any Pro-Resilin-GFP signal. In the wing hinge of *D. suzukii* we found six DT spots. Applying a threshold of 500 counts, we lose some DT spots. In *D. melanogaster*, we are left with six and in *D. suzukii* with five spots. Applying a threshold of 1000 counts, we lose additionally one or two spots in each species. Taken together, trivially, MISM combined with a threshold of acceptance may lead to the identification of ambiguous signals that need closer attention. This is useful for the tentative identification of DT spots in various insect species that would need further characterization for confirmation.

## Results

### DT detection in legs of Diptera

To implement MISM, we first recapitulated published DT incidences in the leg of the fruit fly *D. melanogaster* (Lerch *et al*. 2020; 2022). To start with, we examined the DT signal in the trochanter. Two other Diptera species i.e. *D. suzukii* and *D. hydei* were analyzed in parallel (figure 6 and sfigure s1). We detect a bright auto-fluorescence band on the trochanter-femur joint in all cases. This measured signal from the Drosophilae species, more than 30 individuals each, exceeds the DT-to-DAPI difference threshold by 1000 counts. The smaller areas in the femur-to-tibia joint, the tibia or the tarsi including the tendons, did not show any DT signal. Interestingly, the Pro-Resilin-GFP patch on the femur did not coincide with a DT signal above 500 counts (data not shown). In contrast, MISM revealed a new location of DT in the coxa-to-trochanter joint with around 600 counts, where Pro-Resilin-GFP was not expressed (Lerch *et. al* 2020). Other Diptera species like hoverflies (Syrphydae), *Musca domestica* and species of the winter crane flies (Tipulidae) had similar signals in their leg joints (see fig. 6 and sfig. s1). In conclusion, MISM confirmed DT distribution in the fly legs.

**figure 6.**
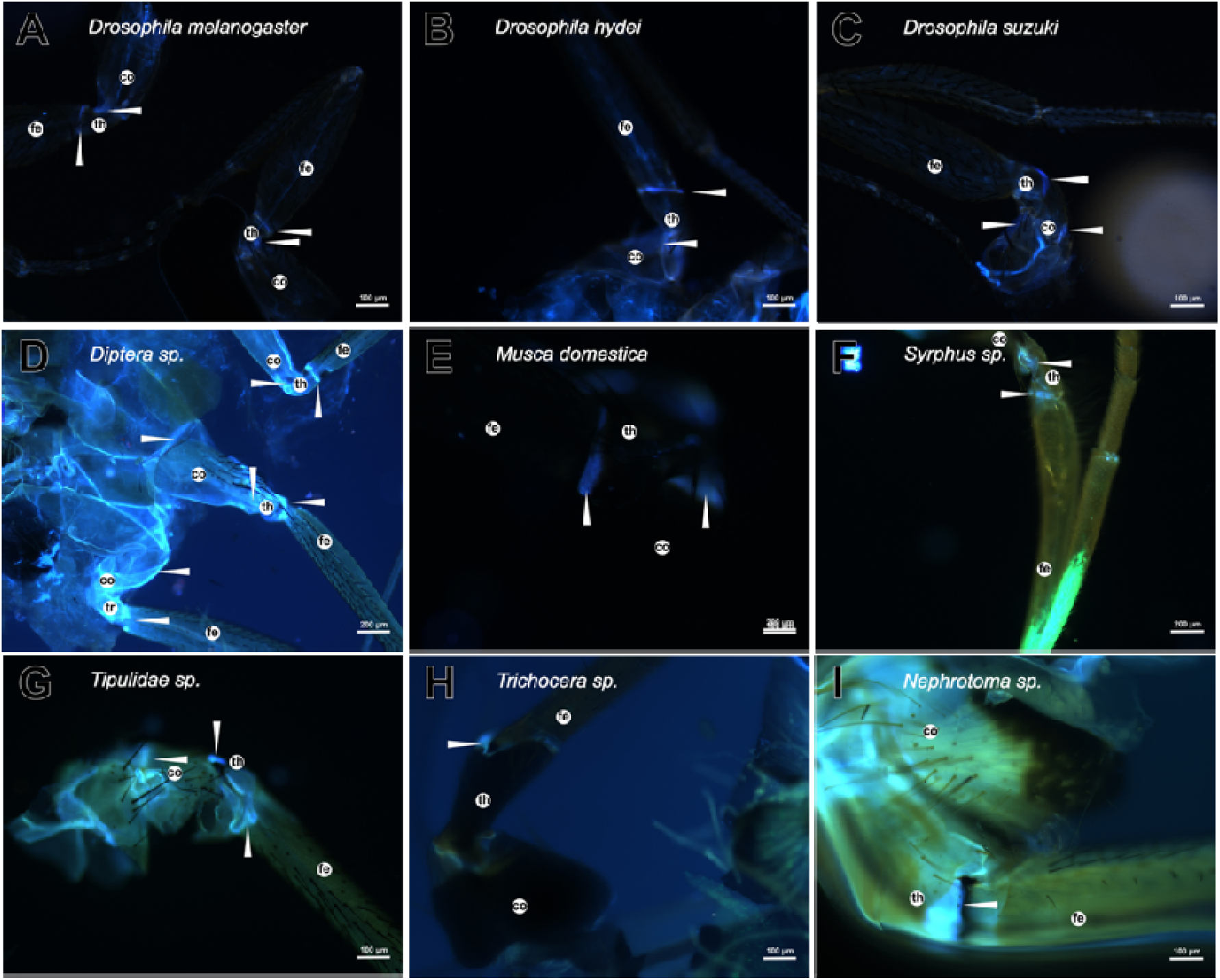
The Diptera legs. Different legs of Diptera species were collected. (A) *D. melanogaster*, (B) *D. hydei* and (C) *D. suzuki*. Other Diptera species show the coxa-trochanter and trochanter-femur DT signals (D-I); the (H) *Trichocera sp.* and (I) *Nephrotoma sp*. samples do not show the signal because of the focus depth. The white arrowheads point to the DT locations. The underlying measurements are shown in sfigure s1. The images were obtained with the inverse Axio Observer Z1 (Zeiss) microscope and the Axiocam Mono camera. co… coxa, fe… femur, tb… tibia, th… trochanter

### DT detection in legs of Orthoptera

As a second sample for evaluation of MISM, we used *S. gregaria*. In this species, we visualized DT in the legs with the settings elaborated in this work (figure 7 and sfigure s2 and s3). As references, the triangle shaped signal of the jump leg and the buckling area in the tibia of the same leg were used (Bayley *et al*. 2012; Burrows & Sutton 2012; Burrows 2016). These incidences were confirmed with a high value threshold. Additional DT singals were detected in the legs of *S. gregaria*. They greatly differ in their morphology and some signals have similar intensity as the tarsal joints and the puvilli display a DT signal over the threshold of 500 counts in all samples. In the trochanter-femur joint of the hindleg of *S. gregaria*, in contrast to the fore- and midleg, we did not detect any positive signal. The fore- and the midleg areas of this species showed a prominent DT signal (over 1000 counts) (see fig. 7 and sfig. s2 and s3). We conclude that MISM validated known DT signals in the desert locust leg.

**figure 7.**
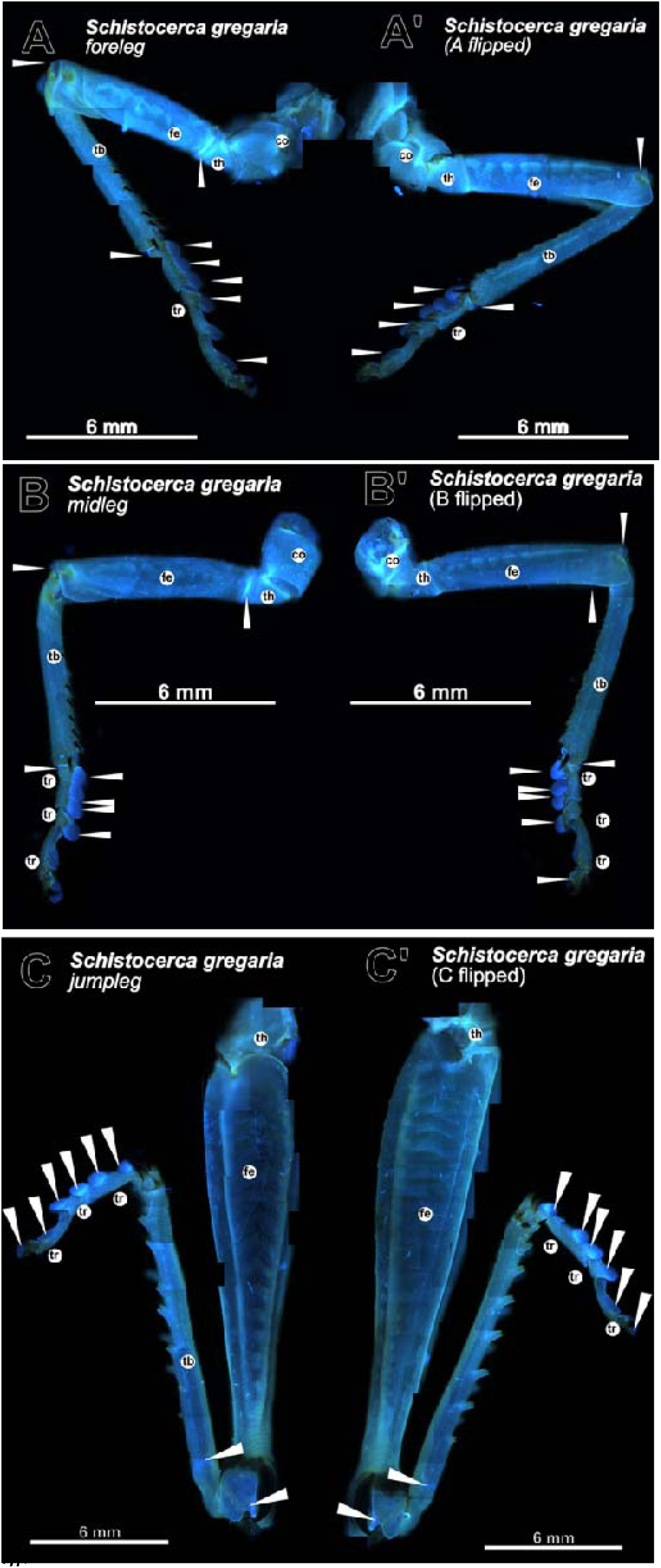
The *Schistocerca greagaria* legs. We present the overview of the (A and A’) foreleg, the (B and B’) midleg and the (C and C’) jump leg. The images in A’, B’ and C’ show the opposite side of their counterparts in A, B and C. All legs show a DT signal in the tarsal adhesive pads and tarsal joins. The (A/A’) foreleg and (B/B’) midleg show a signal in the trochanter to femur, femur to tibia, and tibia to tarsus joints. Mostly, these signals are stronger on one side. An exclusive signal of the jump leg is the buckling area (C/C’) and the system for the jump in the femur to tibia joint (Bayley et al., 2012 and Burrows & Sutton 2012). Obviously, there is a uniform blue fluorescence signal caused by the strong DAPI background signal. DT areas are pointed to with white arrowheads. The underlying measurements are shown in sfigures s2 and s3. This compilation is done by images of the inverse Axio Observer Z1 (Zeiss) microscope with a camera (Axiocam Mono camera). co… coxa, fe… femur, sp… spur, tb… tibia, th… trochanter, tr… tarsi

The respective band-shaped signal was also seen in the trochanter to femur connection of *Gryllus assimilis* (sfig. s4, s5, s6) and confirmed reviewed data (Hayot *et al*. 2013). We noted that the DT distribution of the legs of *G. assimilis* and *S. gregaria* is, however, not identical. The midleg of *G. assimilis* has an oval DT dot with different sizes on each leg side. The bigger ones have a strong signal with over 1000 counts (see sfig. s4, s5 and s6). The femur-tibia joint showed DT areas on the distal edge and the inner area in the front- and midleg only in *S.gregaria*. The triangle of the jump leg of *G. assimilis* is comparably smaller than in *S. gregaria*, moreover, the tarsi are differently formed and lack the adhesive pads in *G. assimilis*. This species does not show the femur-tibia signal of the jump leg (the buckling area after Bayley *et al*. 2012). Like in *S. gregaria*, the jump leg of *G. assimilis* does not show any trochanter signal, whereas the other leg pairs have a signal in the trochanter-femur connection, while in the femur-tibia region the signal is different. The tibia-tarsal joint in both samples contain Resilin (over 1000 counts), only the jump leg of *S. gregaria* did not show any DT in this connective.

### DT detection in legs of Blattodea

*Blaptica dubia* was analyzed as the representative of Blattodea. In the legs of this species, we identified different DT areas (figure 8 and sfigure s7 and s8). Between the coxa and trochanter, there was a triangular signal patch. The distal edge of the trochanter-femur (counts over 1000) was marked by a linear signal with a dot structure at the dorsal side of the leg. The structure from the inside was thicker. A signal in the femur-tibia connection (counts over 1000) was visible as a line and triangle (lateral) and had an area on the downside. We had to open the leg to visualize DT incidences in the adhesive pads that were known to be DT positive (Frazier et al. 1999). The tarsal connections showed a DT signal without dissection. Together, these DT signals were present in all three leg pairs of *B. dubia*.

**figure 8.**
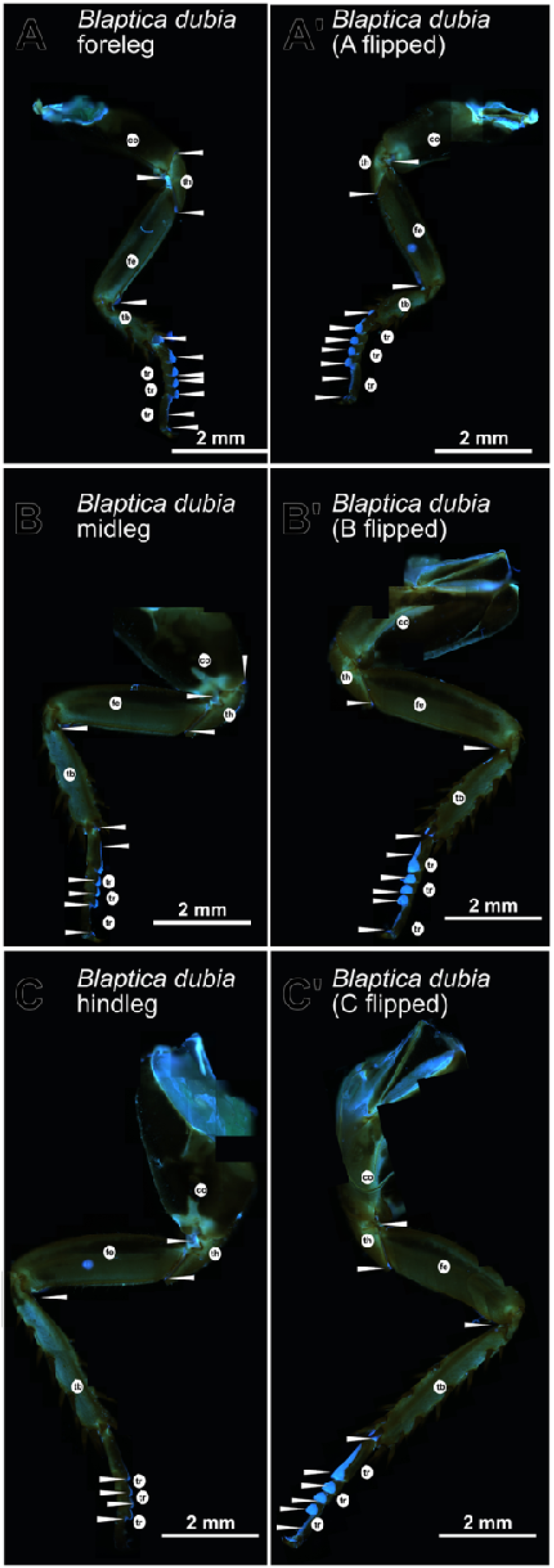
The *Blaptica dubia* legs. An overview of the DT pattern in all three leg pairs shown from two sides: the foreleg (A, A’), the midleg (B, B’) and the hindleg (C, C’) of *Blaptica dubia*. The DT bearing areas are marked by the white arrowheads. All legs are composed of multiple images. *B. dubia* shows a generic morphology in the three leg pairs. The striking difference of their locations is the coxa to thorax structure, which differs in size and shape between the leg pairs. The signals between legs, especially the trochanter signal, are consistent throughout the samples. Besides the opened areas and the DT containing elements, the general cuticle is darker and stiffer and does not contain blue fluorescence. All joints of the legs contain DT, including the tarsal joints and the adhesive pads, visible from inside. DT areas are pointed to with white arrowheads. The underlying measurements are shown in sfigures s7 and s8. An inverse Axio Observer Z1 (Zeiss) microscope were used with a black and white sensitive camera (Axiocam Mono). co… coxa, fe… femur, tb… tibia, th… trochanter, tr… tarsi

### DT detection in legs of Hymenoptera

In Hymenoptera, DT signals only in the antenna cleaner and the tarsi have been reported (Frantsevich & Gorb 2002, 2004; Endlein & Federle 2008; Beutel *et al*. 2020). For the *A. mellifera* legs, we confirmed these data with our method and found additional, yet unpublished signals. The hindleg is different in morphology compared to the midleg and the foreleg especially regarding the tarsi, but the signal in the trochanter-femur articulation was similar in all three types of legs. The DT signal at this position forms an arched band. It is not completely going around the trochanter; the tips often form a hook like structure. We would like to note that by light microscopy, the whole leg is black except the articulation containing DT (figure 9 and sfigure s9). The intensity of the signals was generally over 1000 and never dropped under 500. We detected DT in other areas including the tarsi and trochanter. A clear signal was also found at the coxa-trochanter joint (shape like a fingernail) and the connective area between femur and tibia (triangular). The antenna cleaner was marked by a typical DT signal in the foreleg; at the same position in the mid- and hindleg the signal shape is different. The DT patch was smaller in the hindleg than in the midleg that carries a spur.

**figure 9.**
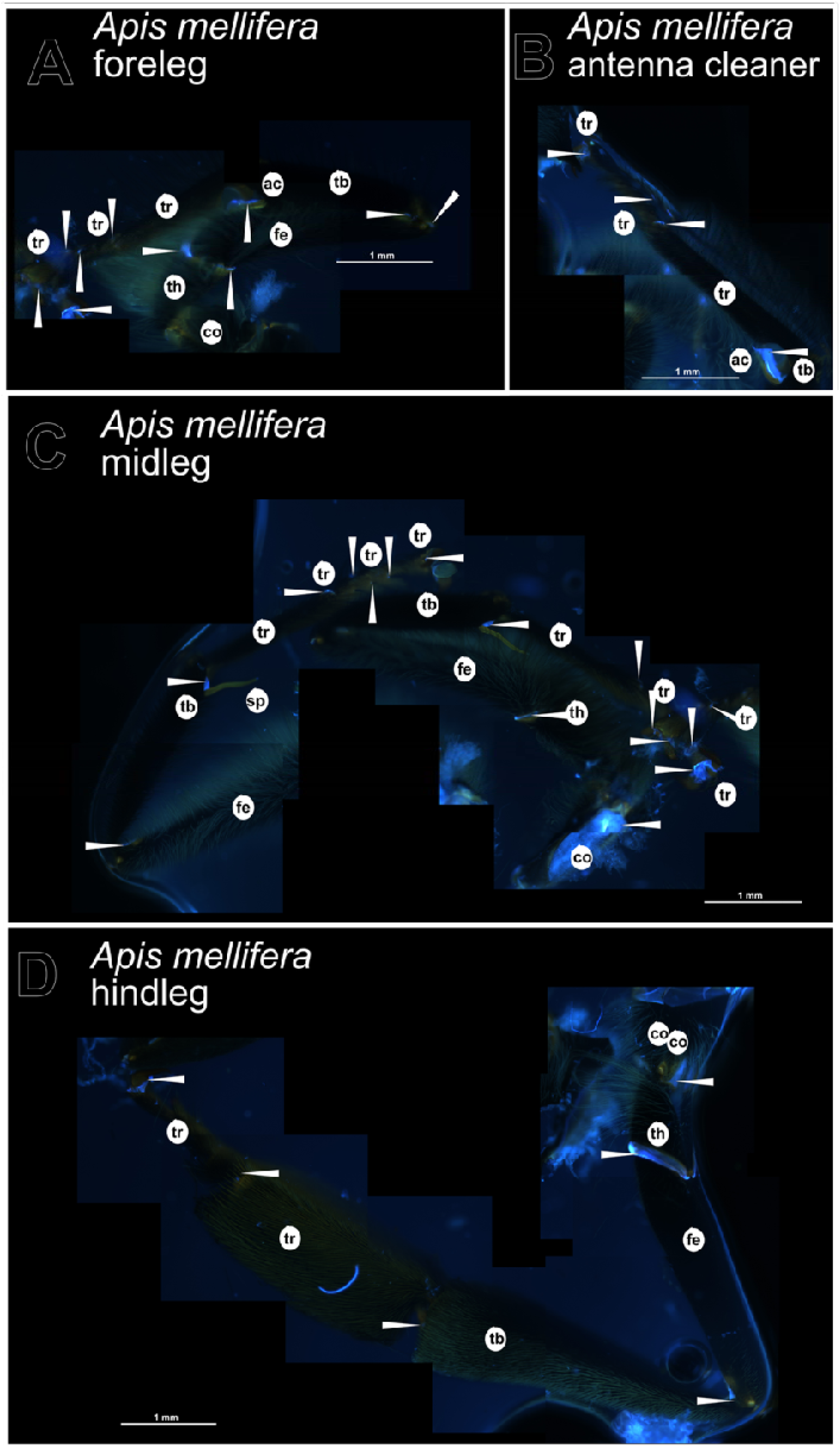
The *Apis mellifera* legs. (A) The whole foreleg of the drone of *A. mellifera* shows DT in all joints including the trochanter-femur and femur-tibia ones or between the tarsal segments. (B) The antenna cleaner also has DT.(C) Two midlegs are shown with all joints containing DT as in the hindleg (D). In addition to the wing vein joints, the wing hinge emits DT signals as shown in sfigure s30 and s31. White arrowheads point to DT locations. The underlying measurements are shown in sfigure. s9. The images were obtained with the inverse Axio Observer Z1 (Zeiss) microscope and the Axiocam Mono camera. ac… antenna cleaner, co… coxa, fe… femur, sp… spur, tb… tibia, th… trochanter, tr… tarsi

Two additional flying Hymenoptera species captured in the field showed a fluorescent patch in the trochanter-femur junction resembling the signal detected in *Apis mellifera*. However, the signal in the presumable *Symphyta* species does not exceed the threshold of 500 (data not shown). A representative of field ants (Formicidae) showed a rather one-sided signal in the trochanter with an over 500-threshold intensity (see sfig. s10 and s11). The antenna cleaner of the ants and *A. mellifera* were different with respect to the DT signal. In ants, the signal is at the basis of the tibia/tarsal joint, while in the bee it exceeds over the calca structure. We note that in the dark samples of ants DT was still detectable.

### DT detection in legs of Coleoptera

The dark cuticle of Coleoptera hindered clear DT signal detection. Indeed, in general, we did not detect a DT signal in all samples. The legs of *P. marginta* are very dark and did not show any fluorescence from the outside (figure 10). Consistently, in the other Coleoptera species studied here, including ladybird beetles (Coccinellidae), leaf beetles (Chrysomelidae), rove beetles (Staphylinidae) and weevils (Curculionidae), no Resilin signal was visible from the outside (see fig. 11 and sfigure s12). In the legs of *T. molitor* and *T. castaneum*, which have a brown cuticle, weak signals were observed in the pretarsi (signal intensity of around 500) and at the joints between the coxa and the trochanter (see fig. 12 and sfig. s13). In both species, the shape of the DT area at the coxa-trochanter position is triangular. We noted that the intensity increased from the front to the hind pairs of legs from the threshold of 500 in the forelegs to the threshold of 1000 in the hindlegs. *T. molitor* and *T. castaneum* larval legs contain DT with above 500 and 1000 counts (see sfig. s14 and s15). DT signal detection was more consistent in a soldier beetle (Cantharidae) species. In these samples, we observed significant DT signals in the coxa-trochanter, femur-tibia and the tarsi (see fig.b12 and sfig. s12). Inspired by previous works (Nadein & Betz 2016, 2018), we sought to verify DT signals inside the legs of *P. magrinata* and *T. molitor*. Indeed, we detected DT spots inside the leg joints in *P. marginata* and inside the coxa-trochanter area of *T. molitor* (sfig. s16 and s17). Because of the dissection process, however, the reliability of these incidences and their shapes is rather weak.

**figure 10.**
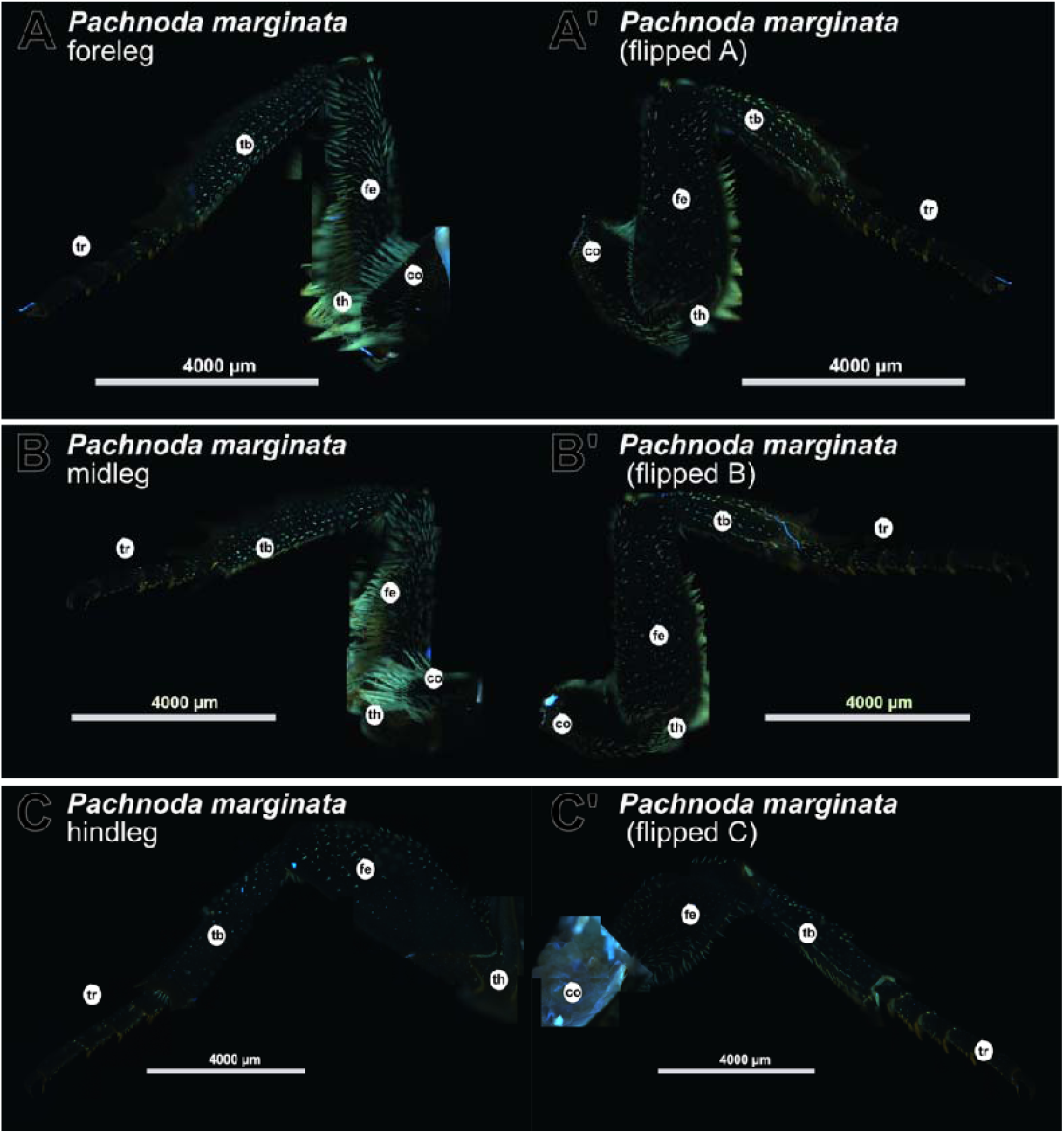
*Pachnoda marginta* leg. All three leg pairs of *P. marginata* are dark. In addition, hairs cover the legs. The hindleg has lesser hairs than the other two type of legs. The foreleg (A, A’), the midleg (B, B’) and the hindleg (C, C’) were analysed from both sides in order to detect any DT signal. However, in all three legs no DT was detectable by MISM. White arrowheads point to DT locations. An inverse Axio Observer Z1 (Zeiss) microscope was used with a black and white sensitive camera (Axiocam Mono). co… coxa, fe… femur, tb… tibia, th… trochanter, tr… tarsi

**figure 11.**
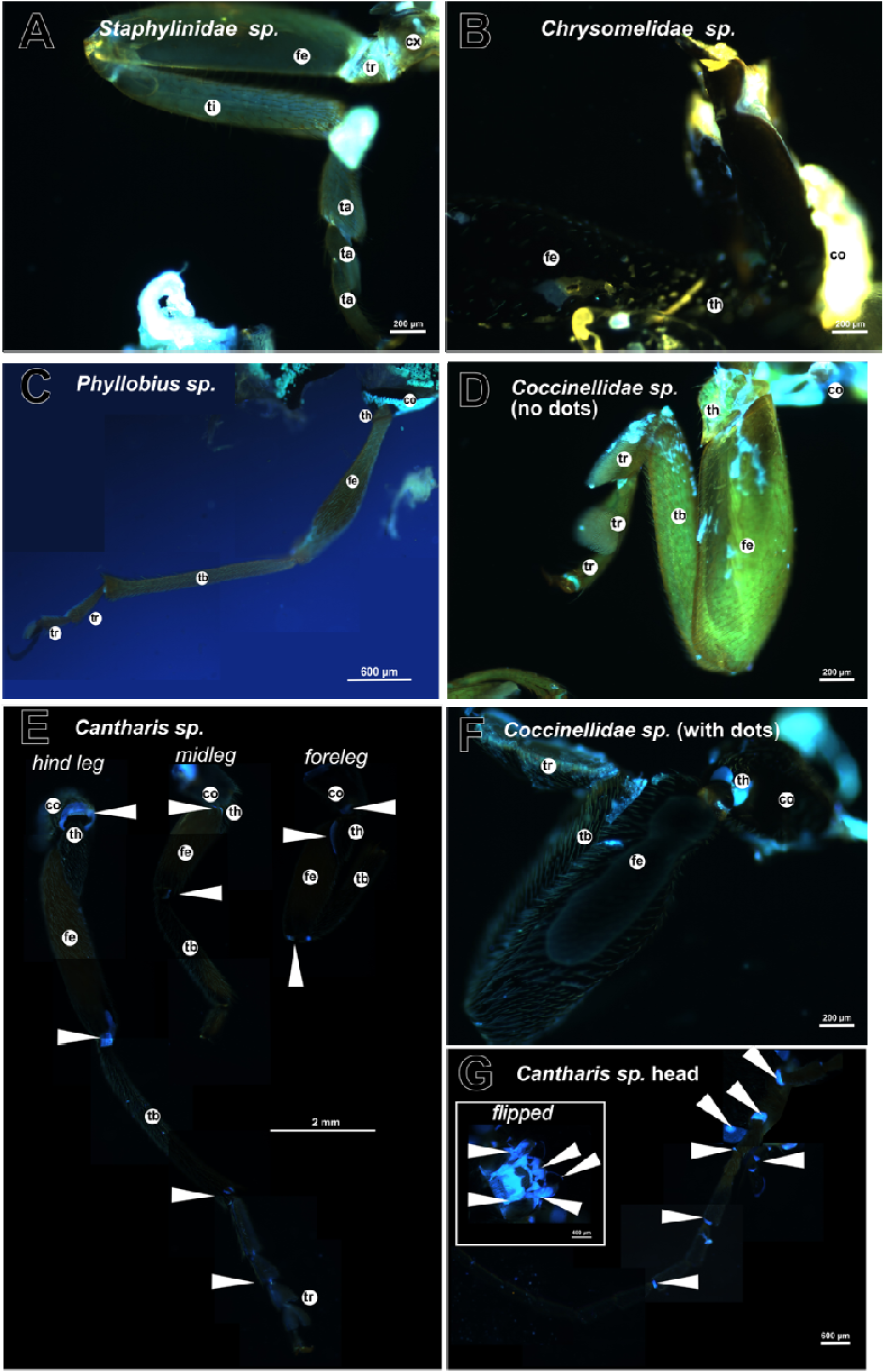
Coleoptera samples. Legs of different Coleoptera species are shown in this figure. The samples of *Staphylinidae sp.* (A), *Chrysomelidae sp.* (B), *Phyllobius sp.* (C), *Coccinellidae sp.* (no elytral dots) (D) and *Coccinellidae sp*. (with elytral dots) (F) exemplify the lack of measurable DT without opening the cuticle. Only the sample of *Cantharis sp.* (E and F) shows DT signals in the three leg pairs (E) and the head (G and the white box of G showing flipped head). The white arrowheads point to the DT signals The underlying measurements are shown in sfigure s12. The images were done by the inverse Axio Observer Z1 (Zeiss) with an Axiocam Mono camera. co… coxa, fe… femur, tb… tibia, th… trochanter, tr… tarsi

**figure 12.**
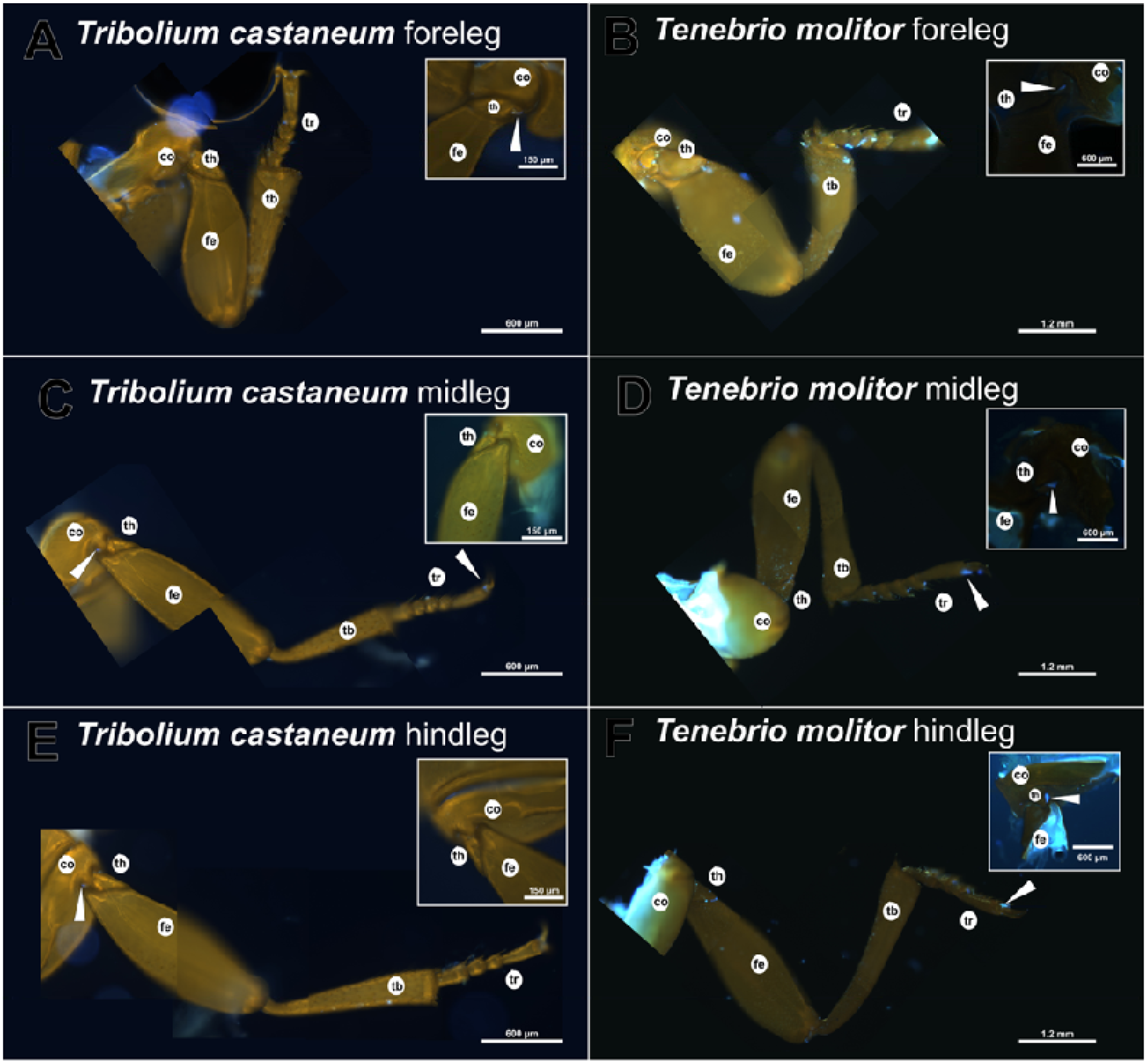
*Tenebrio molitor* and *Tribolium castaneum* leg. In comparison to other Coleoptera, the leg of *Tribolium castaneum* (A, C and E) and *Tenebrio molitor* (B, D and F) emit a DT signal in the area of the coxa to trochanter joint and some praetarsi. To demonstrate that these signals are variable, in the white box, an alternate image of the same coxa-trochanter area is added. The white arrowheads ‘point to the DT locations. White arrowheads point to DT locations. The underlying measurements are shown in sfigure s13. The Axiocam Mono with the inverse Axio Observer Z1 (Zeiss) microscope was used. co… coxa, fe… femur, tb… tibia, th… trochanter, tr… tarsi

### DT detection in legs of Lepidoptera

To work on the cuticle of the legs, firstly we had to remove the scales from the surface gently in all in Lepidoptera samples. In these we did not detect any DT signal in the whole leg except for two regions: a spot on the tibia (mostly in one leg in *E. kuehniella* and *P. interpunctella* or the other samples) and a weak signal over 500 counts in the coxa. The tibia-tarsal joint had a spherical shape and lied close to structures named spur. The coxa signal was often shaped as a half-moon (see figure 13 and sfigure s18 to s21). Inspired by our finding in *B. dubia*, we tried to verify an inner DT signal in *E. kuehniella* as well. In opened the legs of *E. kuehniella* undefined DT signals were present in the area of the trochanter (see sfig. s22).

**figure 13.**
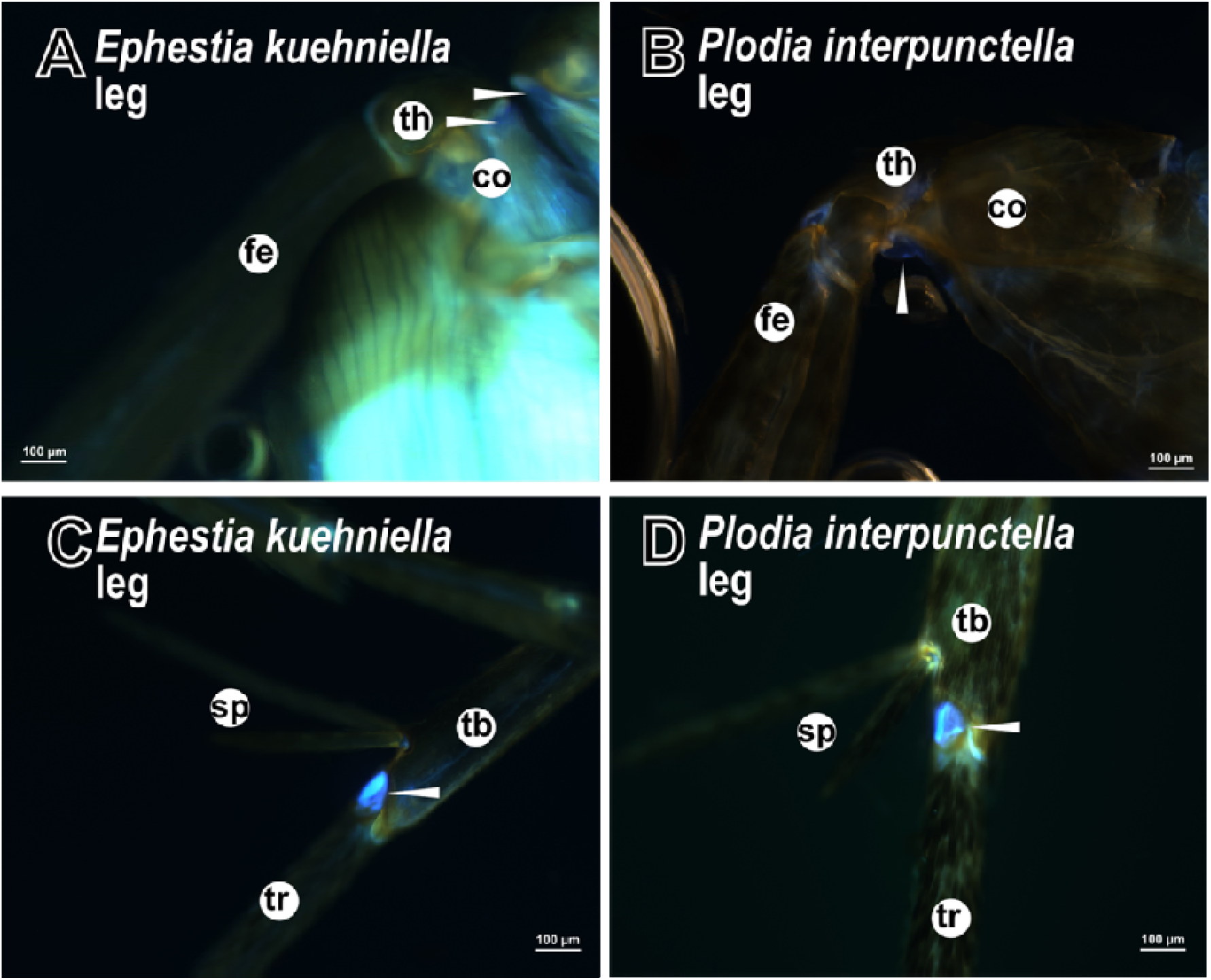
Overview of *Ephestia kuehniella* and *Plodia interpunctella.* The pests (A, C) *Ephestia kuehniella* and (B, D) *Plodia interpunctella* are two moths selected and represent the highest amount of collected species of Lepidoptera. Both species show DT in the same locations, like in wing hinge and the proboscis (see sfigures s41 and s42). No trochanter to femur signal is visible from the outside, but in the joint of coxa and trochanter, we find a signal (A and B). Another leg position with DT is the (C and D) tibia and tarsi connection. For all samples, the removal of the scales from the surface was necessary. All Resilin areas are highlighted by a white arrowhead. The underlying measurements are shown in sfigure s18. The images were obtained with the inverse Axio Observer Z1 (Zeiss) microscope and the Axiocam Mono camera. ac… antenna cleaner, co… coxa, fe… femur, sp… spur, tb… tibia, th… trochanter, tr… tarsi

### DT detection in legs of Hemiptera

In the Hemiptera, the appearance of DT signal is incoherent. DT incidences were found in the legs of *Pyrrhrocoris sp*., probably *Pyrrhrocoris apterus* and other samples of undefined species. Signals in the leg including the coxa-trochanter, the femur-tibia, the tibia-tarsus or between the segments of the tarsi including the praetarsus were found in *Pyrrhocoris* sp. and, in the case of the signal between tarsal segments also in a *Palomena* species. These spots were often triangle-shaped (see figure 14 and sfigure s23). However, these signals were not unambiguously found in all samples (see sfig. s24 and s25). For example, representing two coxae from the same animal, one coxa displayed a DT signal, while the other one did not. The refuted signals may hence represent false-negative incedences. Variation of the DT signal was consistently encountered in Hemipteran species. The intensity variation of these samples is very high, resulting in an unsatisfying identification in the leg.

**figure 14.**
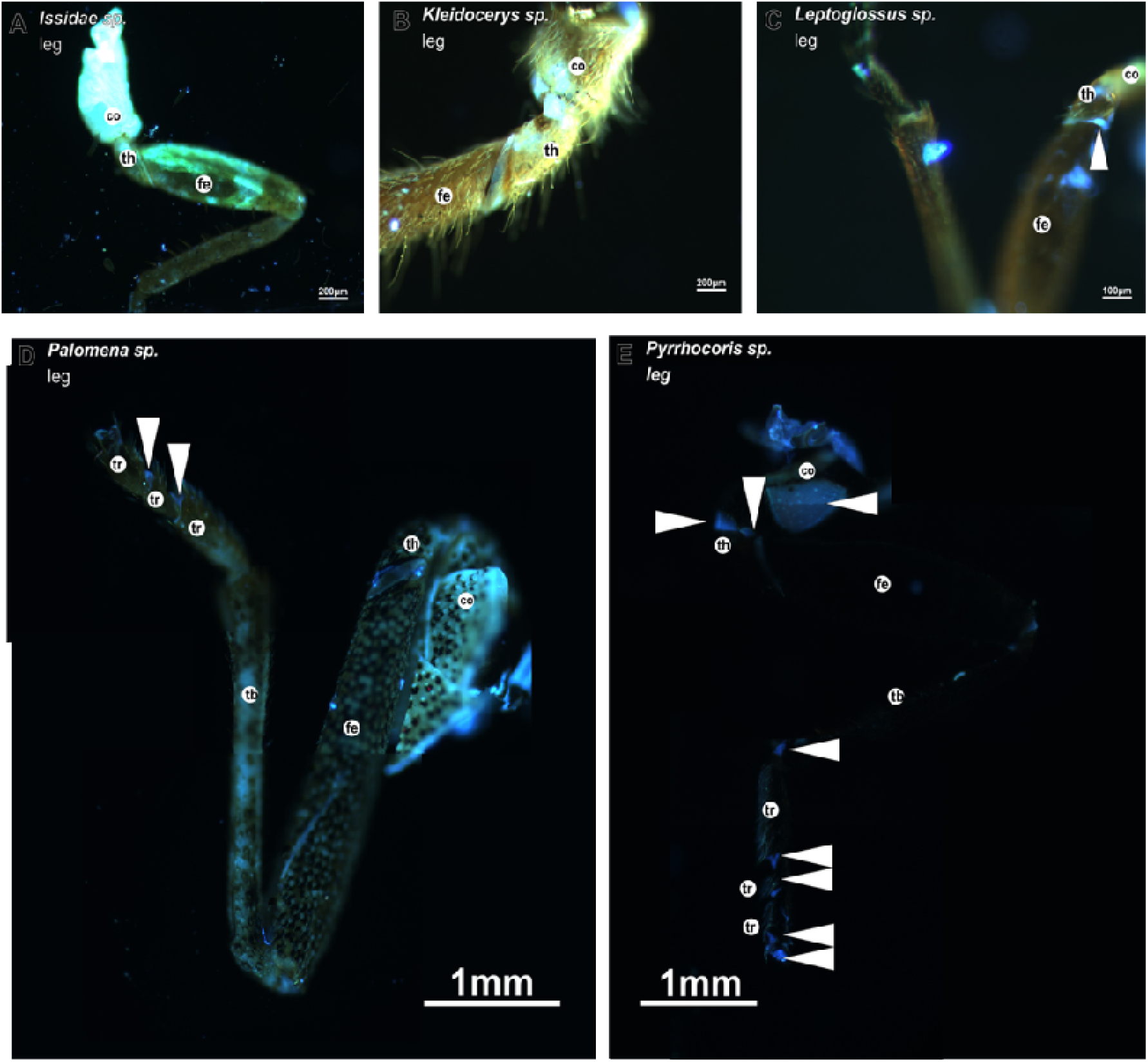
Hemiptera samples legs. The Hemiptera are an inconsistent Resilin containing group. We do not see any DT in (A) *Issidae sp.* and (B) *Kleidocerys sp*. legs. DT in the joint of trochanter to femur in (C) *Leptoglossus sp.* or (D) *Plaomena sp.* tarsal joints is visible. (E) *Pyrrhocoris sp.* occasionally show DT in the coxa, coxa to trochanter, the tibia to tarsi joint and also tarsal joints. These results were difficult to redo. We do not know the reason for this. Consistent DT findings are areas in the wing hinge (see sfigures s43 and s44). The white arrowheads show the Resilin containing locations. The underlying measurements are shown in sfigure s23. All images were done by an inverse Axio Observer Z1 (Zeiss) microscope with a black and white sensitive camera (Axiocam Mono). co… coxa, fe… femur, tb… tibia, th… trochanter, tr… tarsi The DAPI and the DT filters are named in German by the software: “Kanal_1_DAPI. Mittlere Intensität” and “Kanal_5_AF405.Mittlere Intensität”, respectively.

**Figure 15.**
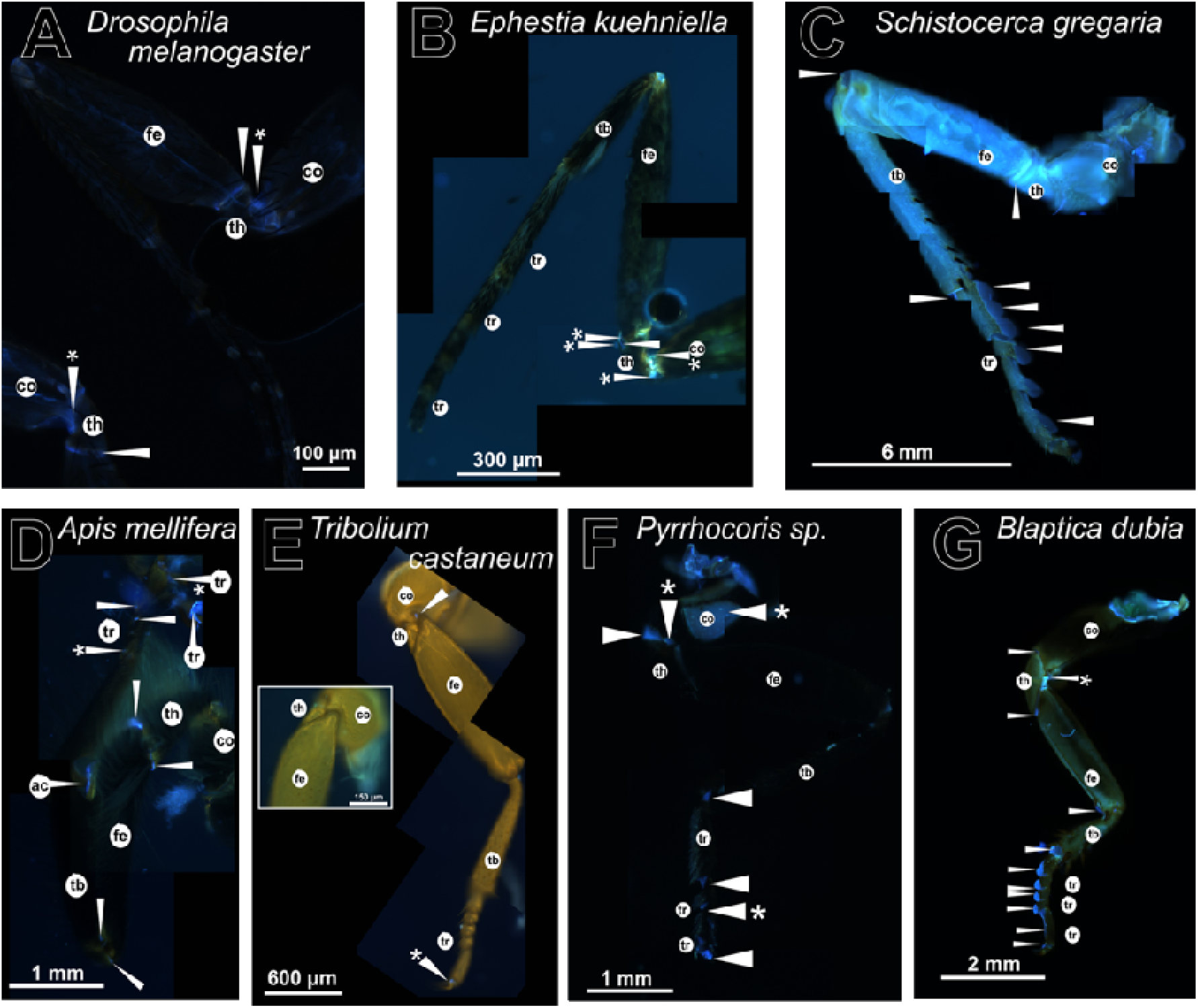
compilation of the Orders – leg. The compilation of images shows the similarities of the morphological structure of the leg. It gives a short overview of common Resilin locations and their differences in area size, intensity and shape. (A) The *D. melanogaster* leg (selected from figure 6 and sfigure s1) has a strong trochanter-femur signal (Lerch *et al*. 2020; 2022) and a weak coxa-trochanter signal. (B) From outside, the leg of *E. kuehniella* shows a coxa-trochanter DT signal (from sfig. s19), but, besides the epiphysis-related signal, no other signal is visible. From the inside, by opening the leg (the crack), different signals with strong and weak components at the area of trochanter-femur joint appear but are difficult to clearly assign. (C) The foreleg of *S. gregaria* (from fig. 7) contains Resilin signals in nearly all joints with higher intensities, except in the coxa-trochanter joint. (D) The honeybee foreleg shows weaker DT signals only in tarsal joints, all other signals are strong (see fig. 9). (E) The midleg of *T. castaneum* has no trochanter-femur signal, this is typical for Coleoptera. As an exception, there are DT signals in the coxa-trochanter and praetarsal segments, which are, however, not consistent (from fig. 12). (F) Inconsistent Resilin locations are shown for the legs of *Pyrrocoris sp.* (Hemiptera). There are strong and weak signals along all connective elements of the leg, but they do not occur in all samples (from fig. 14). (G) The foreleg of *Blaptica dubia* reveals mostly strong signals of DT in all leg joints (from fig. 8). Only the connection of coxa-to-trochanter signal is weaker than 1000 counts between the DAPI and DT filter. At this position, in Diptera, Lepidoptera and *Blaptica* a similar distribution of Resilin and morphology is found. In honeybees and Orthoptera, due to leg polymorphism, Resilin distribution differs accordingly (see the original figures and results). In Coleoptera and Hemiptera, this signal is variably detected. The white arrowhead points to the DT signal, the white asterisk marks the areas of weak signals between 500 and under 1000 counts detected with the DAPI and DT filters. The images were obtained with the inverse Axio Observer Z1 (Zeiss) microscope and the Axiocam Mono camera. co… coxa, fe… femur, tb… tibia, th… trochanter, tr… tarsi The DAPI and the DT filters are named in German by the software: “Kanal_1_DAPI. Mittlere Intensität” and “Kanal_5_AF405.Mittlere Intensität”, respectively.

### Summary of DT in insect legs

In general, we observe that for legs, a stereotypic Resilin or DT signal distribution does not exist in the insect orders examined (see sfigure 15). At some joints, however, like the coxa-trochanter joint, DT is detectable frequently, seen in six out of seven orders. The variability of DT distribution, in general, suggests that DT distribution depends on the mode of locomotion and not on the phylogeny alone. A clear relation between the DT occurrence, phylogeny and moveability of the insect should be further studied to test this hypothesis.

### DT detection in other insect body parts

For all tested orders, we searched for DT signals in other body regions and checked (MISM) with other reference samples, too (see supplementary data s-head, s-wing and other areas). Diversity of DT distribution was also visible in the head (see supplementary s-head data and sfigure s26). A comprehensive analysis of insect heads with respect to DT incidence is missing to date. The only DT-bearing body part that appears to be generally shared by different insect species are the wing hinges (see supplementary s-wing data and sfig. s27).

## Discussion

Considering the number of publications on this matter, we find that the identification of Resilin is a central question in the field of entomology. Two problems have commonly been neglected. First, the incidence of Resilin through microscopic DT detection is indirect and therefore uncertain. Second, non-DT auto-fluorescence signal derived from other components of the cuticle may yield false positive identification of Resilin. This might be especially a problem for weak DT signals. Here, we have developed a new microscopic approach (MISM) to reliably identify Resilin in different insect species. Our strategy is to firstly use the genetic and histological information on Resilin in the fruit fly *D. melanogaster* to specify microscopic settings for powerful DT identification. This involves the subtraction of potentially background signals from a DT specific signal applying two filter sets. Two thresholds were defined by this measure. Based on this, we next applied MISM to different species from different orders including species that had previously been extensively examined on this matter and were, therefore, used as secondary references in this work. Finally, with growing experience, we analysed DT incidences in species that had not been characterized in this field to date. We are aware of the problem of exclusion of weak DT signals by our method. We propose the use of additional imaging methods including time-resolved measurements (Reinhardt *et al*. 2017; Mouchet *et al*. 2023) and Raman spectroscopy (Woodrow et al., 2024) for an accurate analysis of these false-negative signals. We would like to emphasize that identification of Resilin in different insect species by a single method, as presented here, facilitates conclusions as compared to the approach of comparisons across publications and respective methods (see figure 16).

**figure 16.**
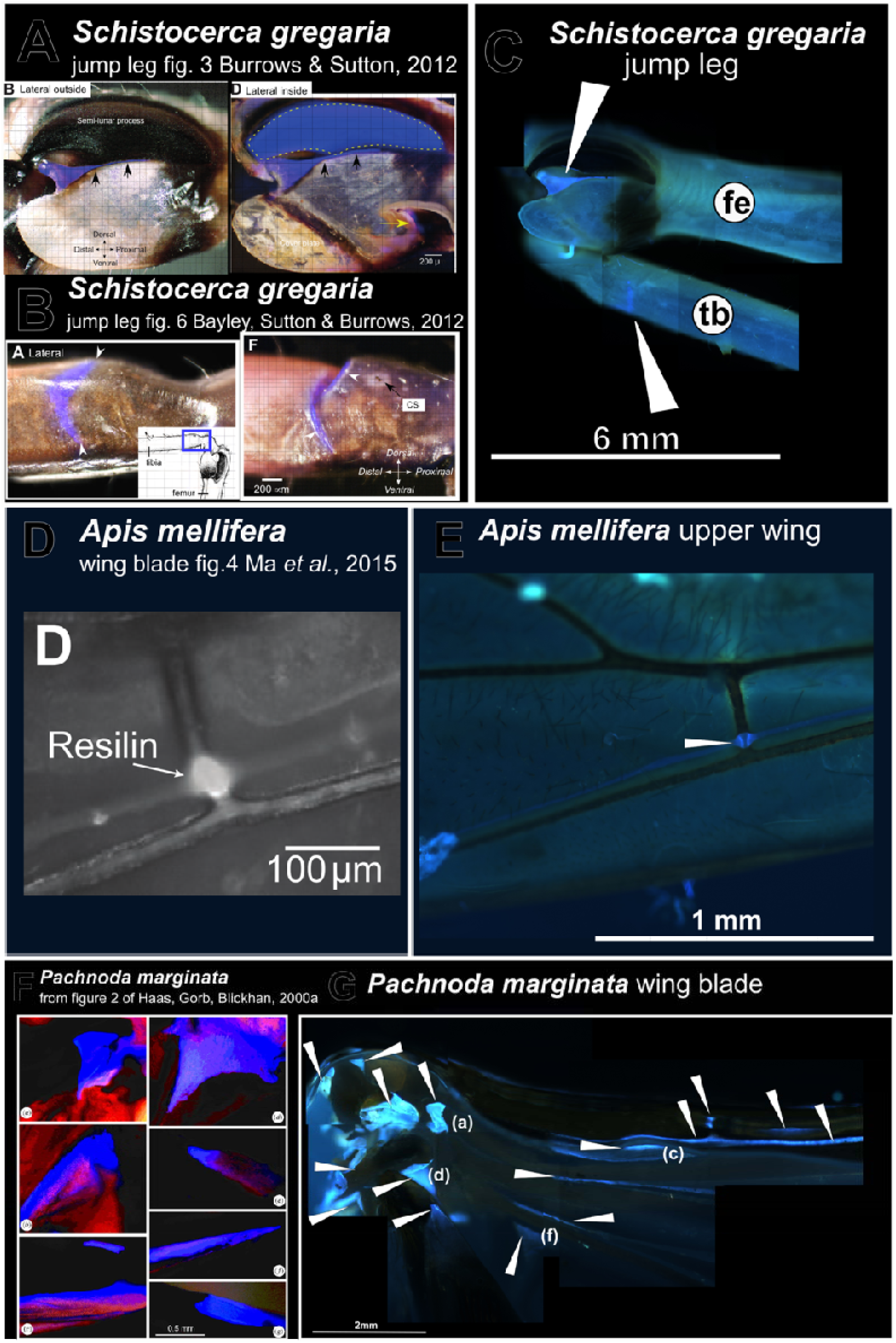
Comparison of reference data of published images with data measured in this work. To validate our method and to define the signal acceptance threshold, published examples were used as references. (A) The semi-lunar process and (B) the buckling region of *S. greagera* (Sutton and Burrows 2012, Bayley, Sutton and Burrows 2012) have been described in the literature; these two areas were choosen as references (see fig. 3 A Sutton and Burrows 2012). (C) In our measurements, both areas show high intensities with over 1000 counts (from figure 7). (D) In the wing vein of *A. mellifera,* DT is present (Ma *et al*. 2015) and (E) and is used as references in our work showing high intensities of over 1000 counts (from fig. 9). (F) In the wing blade of *P. marginata,* different DT areas have been described (Haas, Gorb, Blickhan 2000). (G) We were able to replicate them in our approach (from sfigure s36, a, c, d and f). These DT areas had over 1000 counts. The white arrowheads point to the DT locations. The images (C, E, G) were obtained with the inverse Axio Observer Z1 (Zeiss) microscope and the Axiocam Mono camera. co… coxa, fe… femur, tb… tibia, th… trochanter, tr… tarsi

### Demarcation of the problem

The *D. melanogaster* Pro-Resilin polypeptide has 36 tyrosines (Ardell & Andersen 2001) that potentially form DT or TT bonds with neighboring Pro-Resilin polypeptides. These bonds are responsible for the auto-fluorescence of Resilin matrices and their physical properties. The actual number of DT or TT per Pro-Resilin unit is unknown. Assuming that Resilin matrices consist of other proteins such as Cpr56F, the DT and TT composition of a Resilin matrix becomes very complex (Andersen 2010). For silk moths (Dong *et al*. 2025) or mosquitos (Ohkubo *et al*. 2023), similar diversity of *pro-resilin-*like sequences was reported. In principle, the amount of deposited Pro-Resilin as well as the number of DT and TT bonds may serve to define the physical properties of a given Resilin patch reflected by its auto-fluorescence intensity. Confirming this view, a correlation of the intensity and physical properties of Resilin has been reported using synthetic Resilin (Elvin et al. 2005). In addition, differences in DT and TT amounts and their ratio in different body parts of the locust *S. gregaria* were determined by HPLC (High Performance Liquid Chromatography) indicating different physical properties in the respective regions (Andersen 2004). Quantification of DT and TT is hence crucial to understand the function of a Resilin containing cuticle. To improve DT and TT network quantification, the general overview of Resilin in legs in insect orders requires standardized methods. MISM should serve to open the field for more insects beyond *D. melanogaster*.

### Non-Resilin DT signal

Since, in our initial approach, we had started with the correlation of the DT signal with the presence of a GFP-tagged versions of Pro-Resilin in *D. melanogaster*, identification of DT signals was biased when we refuted DT signals that did not overlap with the GFP signal. Therefore, we sought to revise our method to avoid Pro-Resilin-GFP focused DT localization. In Lerch et al. (2020), we already presented observations hinting in this direction: first, we showed that there is no strict correlation between Resilin-GFP and DT intensities. Second, in *pro-resilin* knock-down flies, the DT signal was not abolished and was significantly higher than the background signal. Together, these findings argue that non-Pro-Resilin proteins may be part of DT matrices thereby determining the region-specific amount of DT. A second explanation for this observation is, in the extreme case, that Pro-Resilin is not necessarily associated with DT. Working on the topic, we saw that in tarsi of *D. menalogaster*, Resilin-GFP occurred without measurable DT. Today, with the new approach of identifying Resilin-networks by DT detection, we are confident of the finding of the coxa-trochanter signal in Diptera. This new area of DT has counts between 500 and 1000 and did not correlate neither with Resilin-GFP nor with Cpr56F-GFP in *D. melanogaster*. This coxa area occurred in different other Orders (e.g. Blattodea and Lepidoptera), as well. This is a very important finding of a non-Resilin DT signal as it indicates that the term Resilin might be misleading. Inversely, there are Pro-Resilin-GFP patches in *D. melanogaster* without a convincing DT signal: the dot in the femur, the tibia-tarsal joint or the tendon in tarsi (Lerch *et al*. 2020). In conclusion, the coincidence of DT and Resilin is not absolute; each case should be analysed separately.

### Identification of Resilin: Coxa-trochanter

The coxa and the trochanter together constitute the insect hip necessary to move the leg with respect to the thorax. Usage of the hip differs between orders, possibly impacting Resilin/DT localization. In Diptera and Lepidoptera a broad halfmoon like shape of DT signal is detected in the coxa-trochanter joint. As in *Apis mellifera* (Hymenoptera), in *B. dubia* (Blattodea) and *Cantharis sp.* (Coleoptera, in the foreleg), the DT signal forms a fingernail shaped domain at this position. In Tenebroids, a respective triangle shaped signal was observed in all leg pairs of adults and juveniles. We noticed a tendency of increase of signal intensity from the fore- to the hindleg. Triangular DT signals were also detected in the coxa-trochanter joints in Hemiptera, however, not in all samples. Finally, Orthoptera in general do not have any DT in the coxa-trochanter joint region. In summary, the DT signal in the coxa-trochanter region is variable between and within orders. This is in contrast to signal areas in the wing hinges, where we always found DT signals (see supplementary s-wing data) The differences may reflect differences in locomotion. Honeybees and cockroaches move in narrow habitats, while flies and moths rather fly and walk only short distances. In Orthoptera, the coxa seems to be made of multiple sclerites joint to the trochanter without an arthrodial membrane. This arrangement may maximize the energy output of the jump. The differences may also be due to the age of the individuals (Lerch *et al*., 2020, Kristensen 1968, Sutton, Burrows 2012) or on pigmentation reducing the fluorescence signal by dimming the DT signal (see supplementary discussion on the dimming problem). Tests on the mechanics of this structure could add more data on this issue supporting the assignment of Resilin. According to the literature, some Hemiptera including planthoppers have a specific signal close to the trochanter that forms a protrusion that becomes visible by opening the leg (Burrows 2010). This signal is not shown in our samples.

### Identification of Resilin: Trochanter-femur

Looking in the next joint between the trochanter and the femur, the DT signal incidence in different Diptera samples including *Drosophila* species and *M. domestica*, the winter crane flies (Tipulidae) and hoverflies (Syrphydae) was similar. All leg pairs showed a trochanter signal. This confirms data published by Lerch et al. (Lerch *et al*. 2020 & 2022). We would like to note that in smaller Diptera (e.g. Drosophila species or Tipulidae) signal variability within a species was low, while in bigger species (e.g. *M. domestica*), it was rather high. Other unidentified Diptera showed a range of the DT signal, not depending on the size, but maybe on melanization of the leg. The crescent shape in *D. melanogaster* was very common (Lerch *et al*. 2020, 2022).

Similar results were obtained for the trochanter-femur joint in Hymenoptera. All leg pairs contained DT, often in a band-line form. The signal is not consistent in size and shape in the three pairs of legs. This may be due to the massive morphological adaptation of them in many Hymenoptera. Comparing ants and bees, we mark that the pigmentation of the cuticle leads to different intensities. It is interesting that in *A. melifera*, the areas with DT signal were unpigmented while they were melanized in the legs of the other samples (mostly ants).

Orthoptera species contain Resilin in trochanter to femur in the fore- and midleg pairs. The data from Hayot et al. (Hayot *et al*. 2013) with *Gryllus firmus* and our data with *Gryllus assimilis* confirm this statement. We also found that the jump legs did not contain Resilin on the edge to the femur.

In the Blattodea sample of *B. dubia* the trochanter was visible as a dorsal dot with linear lateral extensions somehow comparable to the broader signal found in *D. melanogaster*. This finding contradicts Zill *et al*. 2000. These authors observed the trochanter of *Periplenata americana* but did not find any DT signal. These are different species; however, for the other leg parts (see later) the signal is similar. The internal shape of this DT structure in *B. dubia* is slightly broader than viewed from the outside. The manipulated membrane containing DT may be stretched, whereas, when unmanipulated, it may be folded. Alternatively, this area is covered by pigmented sclerites that may dim the signal (see also supplementary discussion on the dimming problem). Both interpretations indicate that the cuticle composition in DT areas should be investigated in detail in order to evaluate DT incidence.

In most Coleopteran species there is no signal in the trochanter-femur joints findable. An outsider is the *Cantharis sp.*, which shows DT at this position. By contrast, in this location, the Lepidopteran species do not show any DT signal. For both Coleoptera and Lepidoptera, we opened the legs in representative samples (*P. marginata* and *E. kuehnella*). Like in *B. dubia*, a DT signal occurs, which is not visible from the outside. This could question the reliability of MISM for strong pigmented or very thick cuticles. We discussed the loss of signal by reduced intensity in the supplementary data on the dimming problem.

For the trochanter as an element between the coxa and the femur, we adhere to the hypothesis that Resilin provides an energy storage for walking and jumping supporting the idea of a general dampening system in the leg for vertical movement (Snodgrass 1935; Frantsevich and Wang, 2009). The trochanter itself is not needed for the movement, but the connection to the thorax via the coxa and the tarsi via the femur-tibia system seems to be important for locomotion. Deora *et al*. (Deora *et al*. 2017) showed in Diptera that the thorax itself does not contain Resilin for flying mechanism. So maybe besides the wing hinge, the trochanter is assisting by the distribution of forces for this process. In general, Resilin in the trochanter has no common shape. We reckon that the shape of this area corresponds to the functional need that in turn depends on the size and the weight of the insect. This may explain that insects like Lepdioptera, which mainly fly and do not walk, do not need an uncovered trochanter Resilin area in the outside. This speculation remains to be investigated in detail.

### Identification of Resilin: Femur-tibia

DT incidence in the femur-tibia joint is highly variable in insect orders. To date, there is no systematic and comprehensive study published about this issue. In *D. melanogaster* Resilin as visualized by a GFP-tagged version of Pro-Resilin, was detected in this region; however, it was not associated with DT-auto-fluorescence. MISM did not indicate DT incidence at this position. In other samples, for example in Coleoptera species like *Lethrus apterus* and *Pentodon idiota* (Frantsevich *et al*. 2019), the leg had to be dissected to reveal DT. In *S. greagaria*, it was shown that a major part of DT is masked by the semi-lunar process (Burrows and Sutton 2012). This exemplifies the problem of signal dimming (discussed in a supplementary discussion on the dimming problem). Likewise, the DT-positive tendon in the femur and tibia could also be identified only upon dissection of the tissue (Burrows 2016). By contrast, the buckling region on the tibia was identifiable without preparation (Bayley *et al*. 2012). Besides, we were able to confirm the triangular region in the jump leg of *S. gregaria* (Burrows and Sutton 2012). Additionally, we can show that other Orthoptera may also have a semi-lunar process represented by the triangle signal in *G. assimilis*. This suggests an evolutionary conserved jumping mechanism in some Orthopteran species. The other leg pairs of *S. gregaria* show DT in the femur-tibia connective, too. For Blattodea, represented by *B. dubia*, the first appearance of DT in this joint of all leg pairs is shown in contrast to the findings in *P. americana* (Neff *et al*. 2000).

DT is inconsistent in Hemiptera; a discussion on this matter would drift off to speculations and is, therefore, omitted here. Lepidoptera, Diptera and most Coleoptera did not show any DT signal in this joint. As an exception for Coleoptera, *Cantharis sp.* showed a signal at this position. In Hymenoptera, in the leg of *Apis mellifera*, the respective signal is traceable. Commonly, as for the trochanter-femur joint, the variability of the DT signal in the femur-tibia connective may reflect differences in usage. In the femur of *G. assimilis*, the oval dot of both sides presented on the midleg is probably the sound productive organ named tympana as described in Ball *et al*. 1989 (see for comparison sfig. s4 and for *G. bimaculatis* in figure 13.2 A and B of Ball *et al*., 1989).

### Identification of Resilin: Tibia-tarsus

In the tibia-tarsi joint region of *Apis mellifera*, Blattodea, both samples of Orthoptera and Lepidoptera and the Coleopteran *Cantharis sp.*, we detected DT signals. For Blattodea these locations fit with the findings in *Periplaneta americana* (Frazier *et al*. 19999 and Neff *et al*. 2000). In ants (except for the antenna cleaner), most Diptera and other Coleopterans, we failed to find this signal. In Hemiptera, again, the signal is inconsistent. For *D. melanogaster*, ResGFP and Cpr56F-GFP localizations support the presence of Resilin in this area (Lerch *et al*. 2020). In the Coleoptera references (Nadein & Betz 2016, 2018 or Frantsevich *et al*. 2019), tibial-tarsal tendons with Resilin have been described, in our study, however, by opening legs of P. marginata we failed to uncover such tendons. We observed other joint related DT signals in the sun beetle and found in T. molitor in the inside of the coxa-trochanter area signals as well. The presence of inlay DT signals in P. marginata not visible from outside solidified the point of the dimming problem (see supplementary discussion about dimming). Lepidopteran species show a DT signal in this region on the foreleg, which we assume to be connected to the epiphysis on the tibia, which, in turn, is probably used for antenna cleaning (Robbins 1989 and Perini *et al*. 2019). It is described that the behaviour of cleaning of the antenna differs between species and the antenna cleaner or tibial epiphysis are present on the foreleg or the midleg or miss altogether (Robbins 1989). In any case, DT in combination with the epiphysis was not mentioned prior to our work. Antenna cleaners are well described in Hymenoptera. In the honeybee, the antenna cleaner is formed by the tibia and the basitarsus of the foreleg and contains DT that is continuous with the joint DT signal. A similar structure is absent in the other pairs of legs (see also Beutel *et al.,* 2020), which, however, do have a tibial-tarsal DT patch. Likewise, in most if not all species examined, at the tibial-tarsal joints, we detected a DT signal. We hypothesize, therefore, that the antenna cleaner and the epiphysis in Hymenoptera and Lepidoptera, respectively, derive both from the tibial-tarsal joint. Another possibility is an origin from the hair bases. Indeed, the hair bases of the grooming device in *Ischnura elegans* (Piersanti *et al*. 2024) and in *D. melanogaster* contain Resilin that might adopt a new function (shown with Resilin-GFP in Lerch *et al*. 2020). Besides the Resilin patch of the antenna cleaner, we detected DT in all joints in the honeybee leg, which are missing in the tested ant species, except for the trochanter-femur joint. These distinct incidences in the honeybee may be associated with locomotion differences. The specific mode of locomotion may explain leg DT differences in the other orders (Orthoptera and Blattodea), as well.

### Identification of Resilin: Tarsi

In case studies, in many different species, tarsi have been shown to contain DT at different positions including pulvilli and joints. However, again tarsal Resilin incidence was not studied systematically and in detail. Early investigations revealed Resilin in the pulvilli of *Calliphora erythrocephal* (Bauchhenß 1979) and *Episyrphus balteatus* (Gorb 1998) between claws and adhesive pads and tarsomer joints in *Musca domestica* and *Eristalis pertina* (Niederegger & Gorb 2003) and were found in tarsal joints and praetarsa of *Eristalis tenax* (Michels and Gorb 2012). The arolium in *Bombyx mori* contains Resilin and is essential for the adhesive function (Dong *et al*. 2025) and shows the occurrence of tarsal Resilin for a Lepidopteran species. With Pro-Resilin-GFP and Cpr56F-GFP lines, we identified these proteins in the tarsal tendons and pretarsa in a previous work (Lerch *et al*. 2020). Hymenoptera tarsi were also reported to contain Resilin (Frantsevich & Gorb 2002; 2004 and Endlein & Federle 2008 and Beutel *et al*. 2020). As an example of Blattodea, *Periplaneta americana* (Frazier *et al*. 1999; Neff *et al*. 2000) was shown to have Resilin in the joints of tibia and tarsi, and for Orthoptera species, within the adhesive pads of tarsi of *L. migratoria* and *Tettigonia viridissima* (Perez Goodwyn *et al*. 2006). In these cases, Neff and colleagues could show DT signals without dissecting their sample, while in the other two cases the insects were opened for analysis. The evidence for DT in tarsal segments of Coleoptera is described in *Coccinella septempunctata* tarsal setae (Pikser *et al*. 2013) and the tarsomeres of females (Michels and Gorb 2012). Hemipteran DT is often shown in adhesive structures (Rebora *et al*. 2018; Reinhardt *et al*. 2019; Rebora *et al*. 2021) or praetarsa (Voigt *et al*. 2007).

In this study, for *S. gregaria*, *B. dubia, A. mellifera* and *Cantharis sp.*, it was possible to stably detect DT in the tarsi. Tenebroids and Hemiptera, by contrast, showed inconsistent DT signals. For other Coleoptera, Diptera and the Lepidoptera no such signals were identified. The adhesive pads on the tarsi of *S. gregaria* and *B. dubia* both show DT and look identical. In other locusts, such adhesive pads have already been described (Perez Goodwyn *et al*. 2006). In all other samples, such kind of structures were absent. Commonly, DT function of the tarsal joints to praetarsa may be dampening, while softness of the adhesive pads is often connected to substrate attachment (Frazier *et al*., 1999; Perez Goodwyn *et al*. 2006; Voigt *et al*., 2017; 2019).

### Conclusion on the identification of DT and Resilin in the insect leg

Summarizing all localizations of DT in the leg in the tested orders, we cannot draw a model of a common DT or Resilin pattern in insect legs. Despite the conserved overall morphology of legs (Snordgass 1935; Boxshall 2004), the physical composition of the leg segments, represented by DT and Resilin, differs between the species conceivably reflecting different gene expression patterns during leg development (Angelini and Kaufman 2005). We reckon that it is possibly dictated by variations in the specific mechanical needs. Similarly, a high degree of morphological diversity and respective DT and Resilin distribution is also seen in the head (see supplementary data s-head). By contrast, DT patches in the wing hinge region are found in all tested orders; however, we did not analyse the extent of homology between these patches (see supplementary data s-wing).

MISM discloses questions about the definition of Resilin. Foremost, we show the incidence of DT in the coxa-to-trochanter joint without the presence of Resilin-GFP in *Drosophila* (Lerch *et al*. 2020). This may suggest that other Pro-Resilin-like proteins form DT and occupy these locations. Shall they be called Resilin? Assigning these signals over the threshold of 500 counts as false-positive, i.e. non-Resilin, would signify a loss of information in many insect species where the Pro-Resilin protein is not histologically identified. Moreover, in most cases where direct Pro-Resilin visualization is not possible, the example of non-Resilin DT signals implies that no DT incidence can be named Resilin unambiguously. Therefore, it is rather reasonable to us to apply the term Resilin-like to DT areas that possibly contain other visco-elastic proteins than Pro-Resilin as found in *Drosophila*, *B. mori* (Dong *et al*. 2025) and other holometabolous insects (Ohkubo *et al*. 2023). The only example of a Resilin-like structure is, hence, the coxa-to-trochanter joint in *D. melanogaster*; in all other cases, this distinction is, strictly spoken, not possible. This problem becomes evident in some insect species such as the bedbug *Cimex lectularius* six Pro-Resilin-like proteins are present (Flaven-Pouchon *et al*. 2024) complicating the assignment of a bona fide orthologue, in turn making it impossible to relate DT incidences to Pro-Resilin matrices alone.

### Co-evolution of Resilin and locomotion

Resilin incidence may be related to the mode of locomotion in insects. In view of the different types of locomotion in insects – crawling, jumping, and walking, this would imply that Resilin distribution is under evolutionary control. The mechanical properties that may be involved in locomotion are energy storage, dampening and protection against fatigue.

Resilin-dependent energy storage has been described in jump mechanisms involving the pleural arch of Siphonoptera (supplementary data other areas; Bennet-Clark and Lucey 1967; Rothshild *et al*. 1975; Lyon *et al*. 2011) or Hemiptera (supplementary data other areas; Rothshild *et al*. 1975; Burrows and Morris 2002; Burrows 2003; Gorb 2004; Burrows 2007, Burrows *et al*. 2008; Burrows 2010; Burrows *et al*. 2011; Sutton and Burrows 2011; Burrows *et al*. 2014; Siwanowicz and Burrows 2017 and Burrows and Dorosenko 2014a), the semi-lunar process in locusts (supplementary data sleg; Bennet-Clark 1975; Burrows and Sutton 2012; Rogers *et al*. 2025), inner leg elements of Coleoptera (supplementary data sleg; Nadein and Betz 2016; 2018; Frantsevich *et al*. 2019) and the furca of Collembola (Nickerl *et al*., 2014; Oliveira 2022). Dampening and stress reduction have been often cited in flight systems but not in legs (Haas *et al*. 2000a; b; Haas *et al*. 2006; Appel and Gorb 2011; Rajabi *et al*. 2016a; b; Kovalev *et al*. 2018; Bäumler and Büsse 2019; Appel *et al*. 2022; Appel *et al*. 2023; see more in supplementary data). Finally, protection against material fatigue has been shown to depend on Resilin as reducing the amount of Resilin enhances the possibility of breaking the wing hinge in flies or the hindleg in locusts (Lerch *et al*. 2020, 2022; Rogers *et al*. 2025) or influences the adhesive capacity of the leg (Grote *et al*. 2024; Dong *et al*. 2025). To what extent the reduced walking efforts in genetic manipulated Resilin flies may be linked to any of the functions of Resilin remains to be investigated (Lehmann *et al*. 2024). In any case, non-full penetration of the phenotype was probably due to compensatory proteins such as Cpr56F in the fruit fly (Lerch *et al*. 2020) or CPR139 in *Bombyx mori* (Dong *et al*. 2025) and seems to be a common theme for holometabolous insects (Ohkubo *et al*. 2023).

The implication of Resilin in leg mechanics i.e. walking, crawling or jumping is evolutionary old as DT signals have been identified in the flightless Collembola and Crustaceans. The predominant Dipteran walking mode is spring based applying a tripod gait with a cycling spring energy storage in three legs (Lehmann *et al*. 2024). This gait is also described for cockroaches, locusts and ants (Chunt *et al*. 2021; Labonte and Holt. 2022). However, the energy distribution of kinematics is not even in the legs of insects based on the diversity of the geometry of the tripod stances, resulting in a variation of Resilin shape and localization (based on Chunt *et al*. 2021). In locusts, for instance, the coxa-trochanter and the trochanter-femur segments of the third leg do not contain Resilin, but they perform the tripod gait. In any case, correlation of DT and Resilin distribution with the tripod gait or other gaits remains to be studied in detail. Indeed, to date, a comprehensive analysis of mechanical behaviour and the function of Resilin is still missing.

The diversity of Resilin incidences in the insect legs is contrasted by the uniformly detected DT patches in the wing hinges of all species studied in this work. This may reflect the evolutionary new establishment of the flight system in insects, whereas leg-based locomotion i.e. walking is ancestral. Indeed, again, DT signals have been found in Collembola (Nickerl *et al*., 2014; Oliveira 2022), a sister-group of insects and Crustaceans (Andersen and Weis-Fogh 1964; Burrows 2009; Michels *et al*. 2012; 2015; 2016; Michels and Gorb 2012; Appel *et al*. 2022). As stated above, the DT patches in insect wing hinges, however, are unclassified to date and need a closer inspection for a solid conclusion on this matter.

The insect trochanter provides an example for how we propose to address the biological function of Resilin in a comparative approach based on our data. Let us speculate that the trochanter-femur Resilin incidence is related to a mode of walking. Flying and occasionally walking insects like bees, moths, and flies with similar leg pair morphology may require a different function of the Resilin incidence in the trochanter than predominantly jumping and walking insects like locust or beetles. By examining the trochanter, shape and location of Resilin on the trochanter-femur contact site appear to be similar in Diptera and Hymenoptera, but not in Lepidoptera where no Resilin signal was detectable at this position. Coleoptera species except the soldier beetle (Cantharidae), also lack a trochanter-femur Resilin signal. Only the first leg pairs in Orthoptera show a respective signal for Resilin. Thus, the distribution of Resilin in the trochanter of species examined here does not allow a conclusion on the incidence of the trochanter Resilin patch and its impact on the type of locomotion. Obviously, one reason for this failure is that our understanding of locomotion may, despite the great number of publications, be rather limited. A more drastic explanation would be that the Resilin patch in this particular case is not involved in locomotion per se but another type of motion like one applied during mating. Another possible evolutionary responsivity of Resilin is exemplified by the presence of the antenna cleaner and the associated DT signal in Lepidoptera and Hymenoptera and their absence in the Diptera, Orthoptera, and Coleoptera samples.

Besides the localization, the different composition of Resilin and the cuticle need to be focused in to understand biomechanics of insect locomotion. An example are visible signals from the inside and not the outside, what we summarized as the problem (see supplementary discussion), these auto-fluorescence fluctuations were also addressed by other authors (Preuß *et al*. 2024; Roze *et al*. 2024). Future studies on Resilin and its function in locomotion should be accompanied by genetic manipulation of Resilin matrix protein expression. This combination added with evolutionary context will allow a higher understanding of movability and how to overcome mechanical stress most efficiently. In this sense, the recent molecular analyses of Resilin in *D. melanogaster* and *S. gregaria* are promising.

## Acknowledgements

We thank the Bloomington Stock Center at the Indiana State University (USA) for stocks. We are grateful to the Microscope facility of the Chair of Systems Biology and Genetics of Professor Dahmann, Institute of Genetics, Faculty of Biology, Technische Universität Dresden.

