## Supplementary material for "Diversity of Resilin incidence in the insect leg": Suppl Data

### Diversity of Resilin incidence in the insect leg - Supplementary Material

### Supplementary Figures

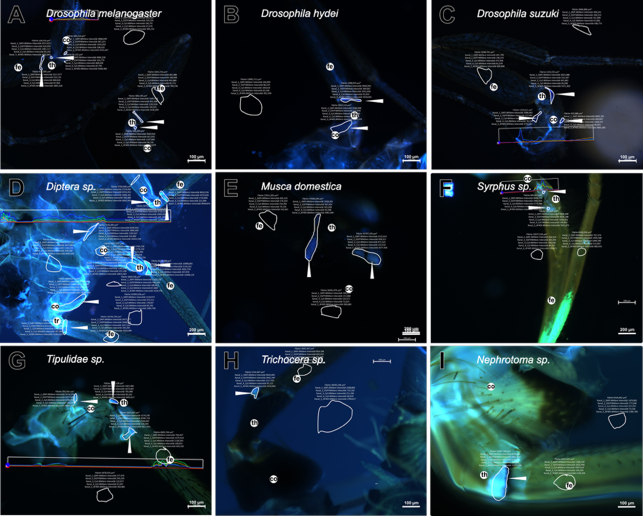

#### sfigure s1 - Diptera leg data.

This figure is a copy of figure 6 including, in addition, the areas of the measurement generated with the profile tool (by ZenBlue) and drawn by hand to include the DT signals.

The DAPI filter is named of the software: Kanal_1_DAPI.Mittlere Intensität. The DT filter is named by the software: Kanal_5_AF405.Mittlere Intensität.

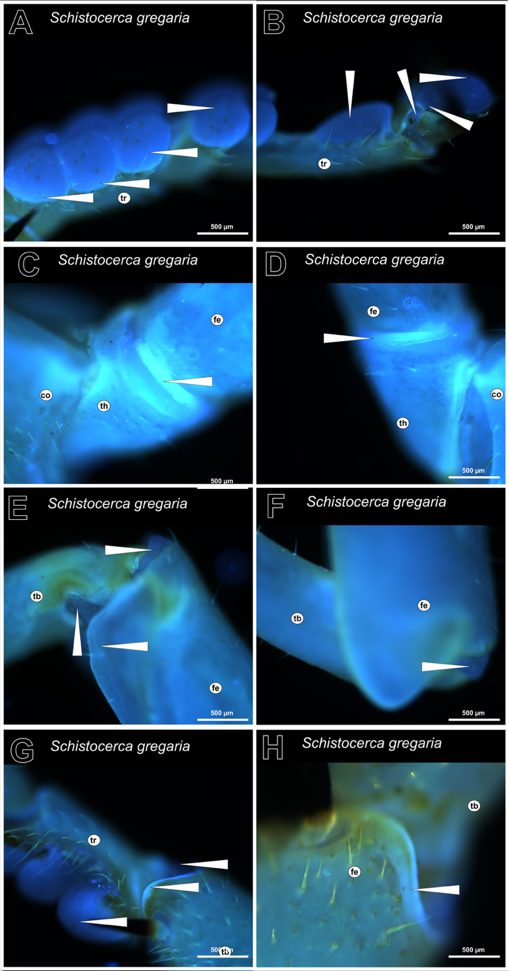

#### sfigure s2 - S. gregaria leg samples.

To show the DT positions in more detail, we present our new findings in this compilation (see figure 7).

(A) DT signal in adhesive pads and tarsi as also shown in (B) including the praetarsus. Focus on the trochanter of the first leg (C) and the midleg (D). (E, F and H) The femur to tibia joint contains DT, especially in the anterior tip, on the femur edges and the lateral connection area between tibia and femur. This is similar in the region between the tibia and the tarsi (G). The signal of the adhesive pads is shown.

The white arrowheads point to the DT locations. The underlying measurements are shown in sfigure s3. The images were obtained with the inverse Axio Observer Z1 (Zeiss) microscope and the Axiocam Mono camera.

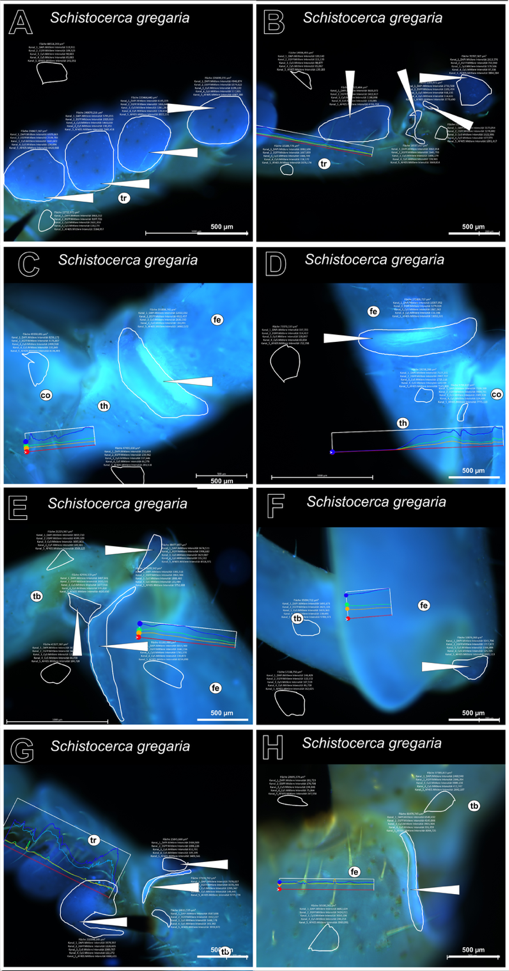

#### sfigure s3 – S. gregaria leg samples data.

This figure is a copy of sfigure s2, including, in addition, the areas of the measurement generated with the profile tool (by ZenBlue) and drawn by hand to include the DT signals.

The DAPI and the DT filters are named in German by the software: “Kanal_1_DAPI. Mittlere Intensität” and “Kanal_5_AF405.Mittlere Intensität”, respectively.

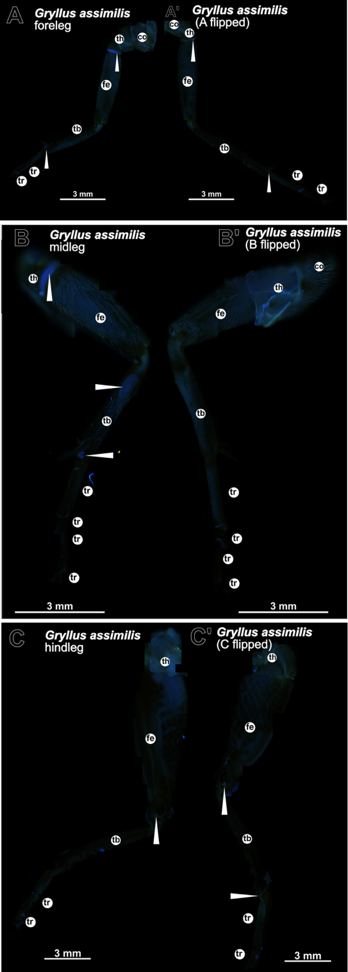

#### sfigure s4 - Gryllus assimilis legs.

The leg has a general blue fluorescence signal, with some darker areas and many hairs covering and darkening the cuticle. Images are compositions of single images.

The left and the right sides of all three leg pairs, the foreleg (A and A'), the midleg (B and B') and the jump leg (C and C'), were analysed for DT detection. Both, the foreleg (A, A') and the midleg leg (B, B') show a trochanter to femur signal. The jump leg (C, C') contains a femur to tibia signal representing the same signal known in the *S. gregaria* jump leg. The joint signal of the tibia and tarsi is visible in all three legs.

The white arrowhead points to the DT signal. The underlying measurements are shown in sfigures s5 and s6. The images were obtained with the inverse Axio Observer Z1 (Zeiss) microscope and the Axiocam Mono camera.

co... coxa, fe... femur, th... trochanter, tr... tarsi

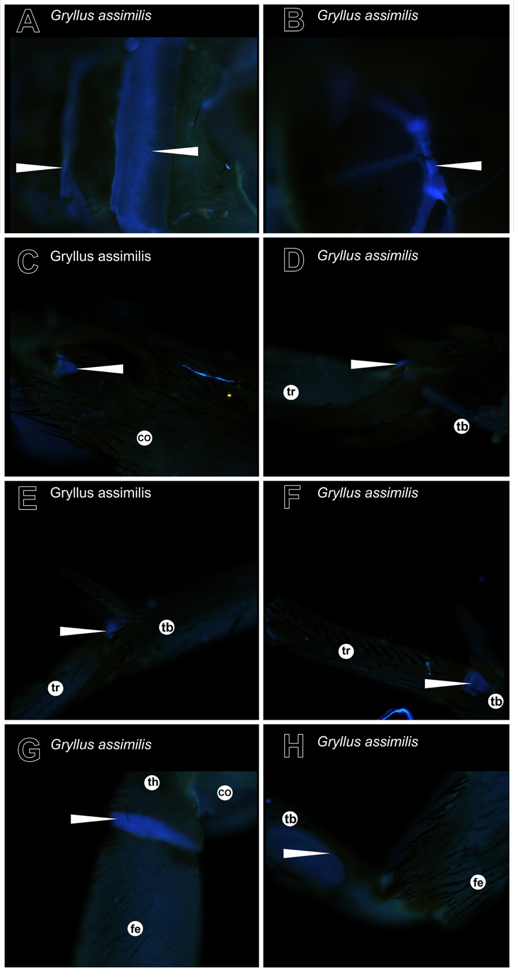

#### sfigure s5 - Gryllus assimilis head and leg samples.

Here, we show the signals in *G. assimilis* in detail, focussing on the head (A and B) or leg joints (C-H). (A) The dorsal side of the head has two band-like areas of DT signal. (B) There is only one broad line on the ventral side of the head. (C) The jump leg shows the typical triangle structure with DT in the femur-tibia joint. The DT signal of the connective between the tibia and tarsi is visible in all three leg pairs (D jump leg, E foreleg and F midleg). The trochanter-femur signal occurs only in the foreleg (G) and the midleg. (H) We noted a unique signal in the upper region of the midleg tibia that may represent a sound productive organ.

The white arrowhead points to a DT signal. The underlying measurements are shown in sfigure s6. The images were obtained with the inverse Axio Observer Z1 (Zeiss) microscope and the Axiocam Mono camera.

co... coxa, fe... femur, tb... tibia, th... trochanter, tr... tarsi

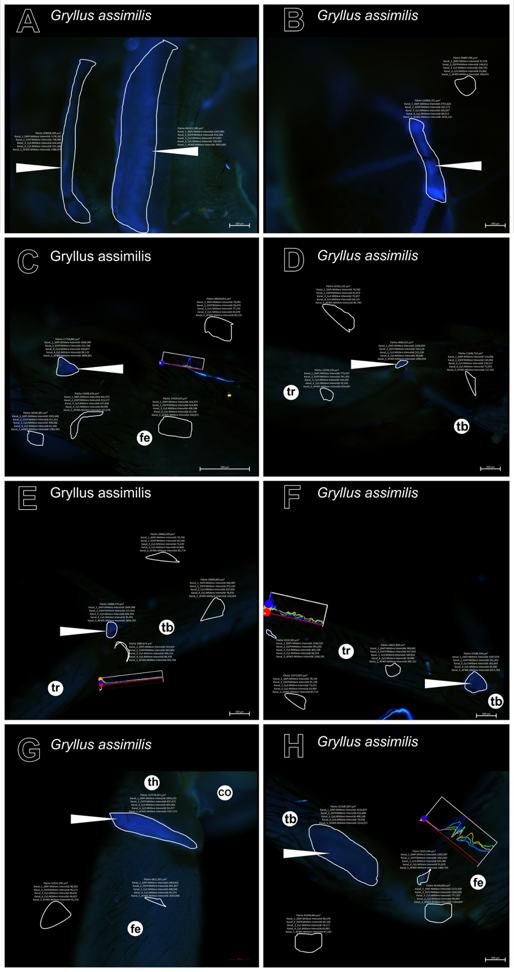

#### sfigure s6 - Gryllus assimilis head and legs samples data.

This figure is a copy of sfigure s5 including, in addition, the areas of the measurement generated with the profile tool (by ZenBlue) and drawn by hand to include the DT signals.

The DAPI and the DT filters are named in German by the software: “Kanal_1_DAPI. Mittlere Intensität” and “Kanal_5_AF405.Mittlere Intensität”, respectively.

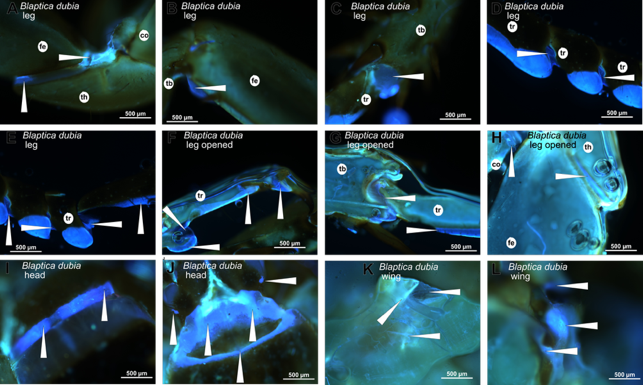

#### sfigure s7 - Blaptica dubia sections of the leg, wing and head.

These images of the legs (A-H), of the head (I and J) and of the wing hinge (K and L) of *Blaptica dubia* show close-ups of the DT pattern. DT locations measured are pointed to by a white arrowhead. (A) The trochanter is marked by the bulge shaped tip on the ventral side of the leg with the dorsal side attached to femur and coxa. (H) From the inside, the bulge in the trochanter is visible, but a signal line to the coxa is more prominent than from the outside. (B) At the femur to tibia connection, the dorsal area is flatter than the bulge joint part of tibia to the first tarsal podomere (C). This bulge is well visible also from the inside (G) of this connection, where a small DT structure connects tibia and tarsi. The tarsal connections bear DT in the joint membrane, the adhesive pads from the outside did not emit DT (E, D). (F) The opened segment of the adhesive pads shows DT by observing the area, it seems that only a partial layer of adhesive pads contains DT. The reason of the dimming of that signal from the outside is not known. I and J are samples of the head with strong incidences of DT. The wing hinge DT signal are shown in K and L.

The white arrowhead points to the signal of DT. The underlying measurements are shown in sfigure s8. The images were obtained with the inverse Axio Observer Z1 (Zeiss) microscope and the Axiocam Mono camera.

co... coxa, fe... femur, tb... tibia, th... trochanter, tr... tarsi

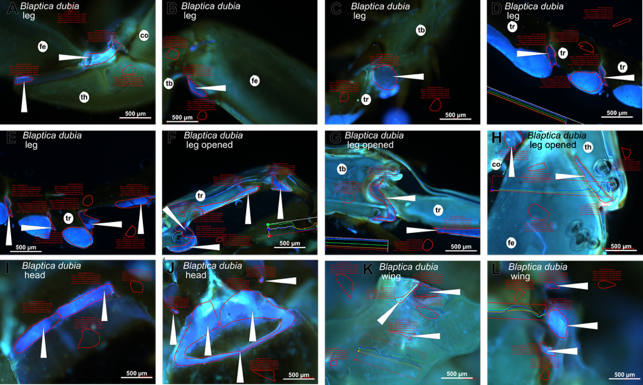

#### sfigure s8 - Blaptica dubia sections of the leg, wing and head data.

This image is a copy of sfiugre s7 including, in addition, the areas of the measurement generated with the profile tool (by ZenBlue) and drawn by hand to include the Dityrosine signals.

The DAPI and the DT filters are named in German by the software: “Kanal_1_DAPI. Mittlere Intensität” and “Kanal_5_AF405.Mittlere Intensität”, respectively.

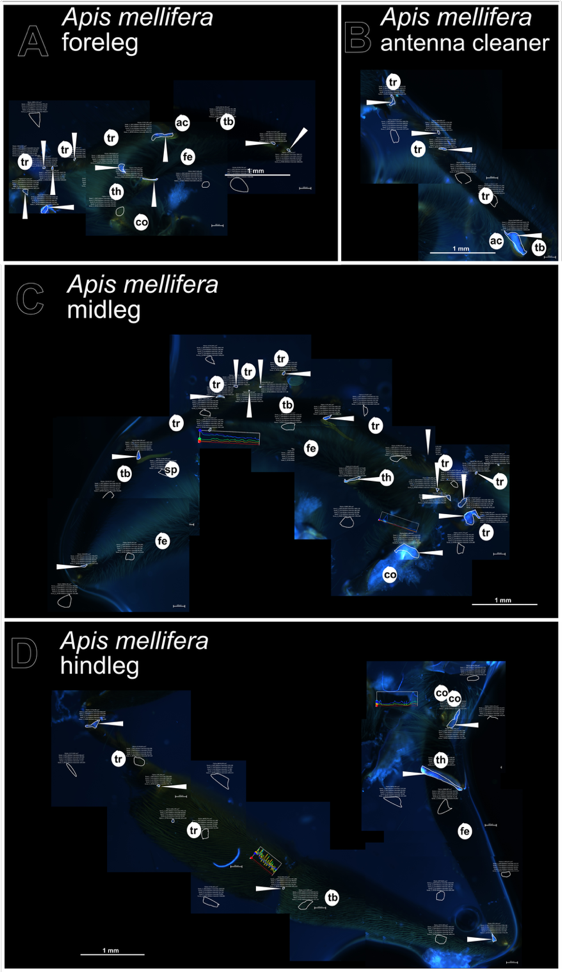

#### sfigure s9 - Apis mellifera wing and leg data.

This image is a copy of figure 9 including, in addition, the areas of the measurement generated with the profile tool (by ZenBlue) and drawn by hand to include the Dityrosine signals.

The DAPI and the DT filters are named in German by the software: “Kanal_1_DAPI. Mittlere Intensität” and “Kanal_5_AF405.Mittlere Intensität”, respectively.

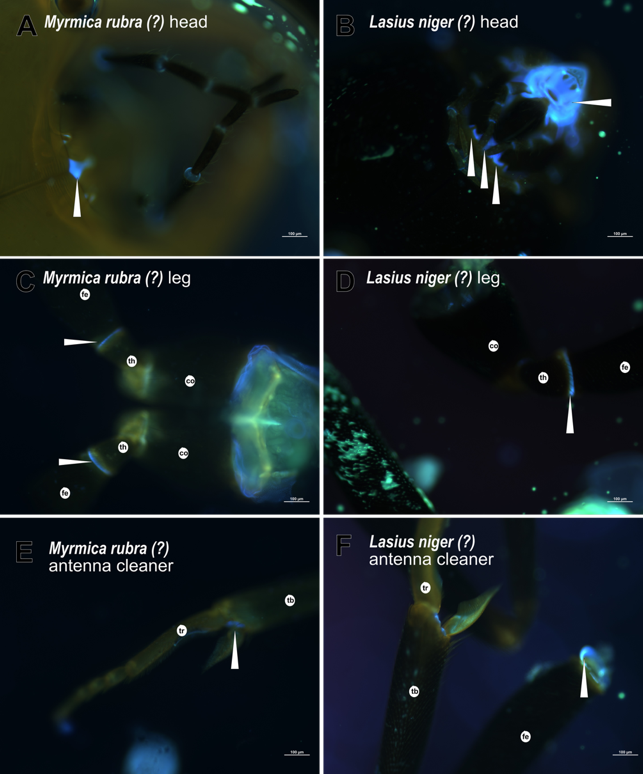

#### sfigure s10 – Ant samples.

DT is present in ants in different body parts.

The ant head contains areas of DT (A and B), and we found DT in the trochanter to femur joint (C and D). In the typical antenna cleaner of the Formicidae, we were able to detect DT (like Beutel et al. 2020) in (E). From the outside, viewed from the distal end, no measurable signal is detected; however, after opening, the leg shows a trochanter/femur signal. The colour of the cuticle seems not alter the signal intensity in the two different ant species.

White arrowheads point to the DT locations which there visualized by using a black and white sensitive camera (Axiocam Mono) in combination with the inverse Axio Observer Z1 (Zeiss) microscope. The underlying measurements are shown in sfigure s11.

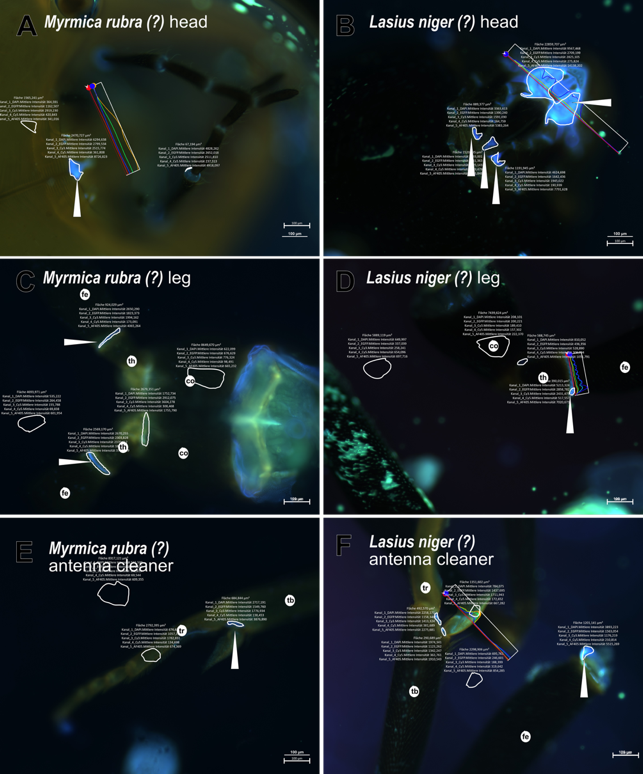

#### sfigure s11 – Ant sample data.

This image is a copy of sfigure s10 including, in addition, the areas of the measurement generated with the profile tool (by ZenBlue) and drawn by hand to include the DT signals.

The DAPI and the DT filters are named in German by the software: “Kanal_1_DAPI. Mittlere Intensität” and “Kanal_5_AF405.Mittlere Intensität”, respectively.

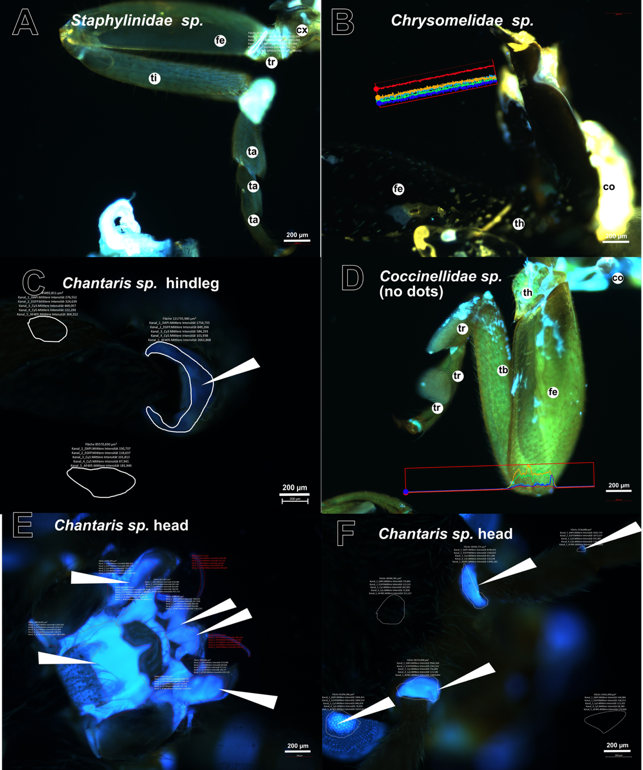

#### sfigure s12 - Coleoptera samples.

This figure shows the measurement data of figure 11 including the DT-lacking samples of *Staphylinidae sp.* (A), *Chrysomelidae sp.* (B) and *Coccinellidae sp.* (no dots) (D). For *Cantharis sp.*, samples are shown including the trochanter of the hindleg (C) and areas of the head (E and F).

The measured DT locations are pointed to with white arrowheads. Images were obtained with the inverse Axio Observer Z1 (Zeiss) microscope and the Axiocam Mono camera. The white arrowheads point to DT.

co... coxa, fe... femur, tb... tibia, th... trochanter, tr... tarsi

The DAPI and the DT filters are named in German by the software: “Kanal_1_DAPI. Mittlere Intensität” and “Kanal_5_AF405.Mittlere Intensität”, respectively.

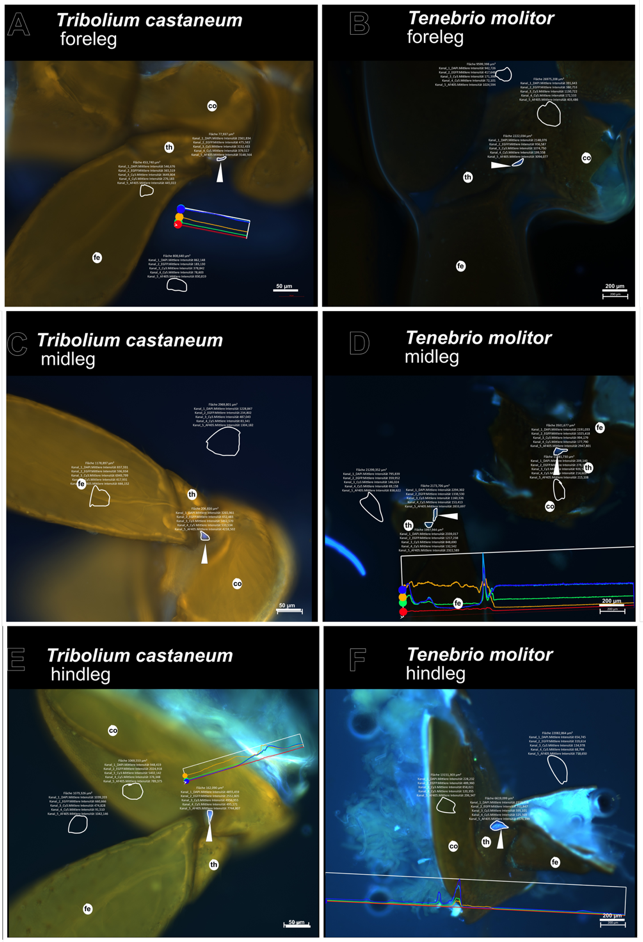

#### sfigure s13 -Tenebrio molitor and Tribolium castaneum leg data.

Here, we show the source data in *Tenebrio molitor* (B, D and F) and *Tribolium castaneum* (A, C, and E) for the DT signal occurring in the region of coxa to trochanter (see figure 12). To show the difference, areas of no cuticle and no DT containing areas were also measured. The white arrowheads point to DT locations.

The Axiocam Mono with the inverse Axio Observer Z1 (Zeiss) microscope were used to obtain the images.

co... coxa, fe... femur, th... trochanter

The DAPI and the DT filters are named in German by the software: “Kanal_1_DAPI. Mittlere Intensität” and “Kanal_5_AF405.Mittlere Intensität”, respectively.

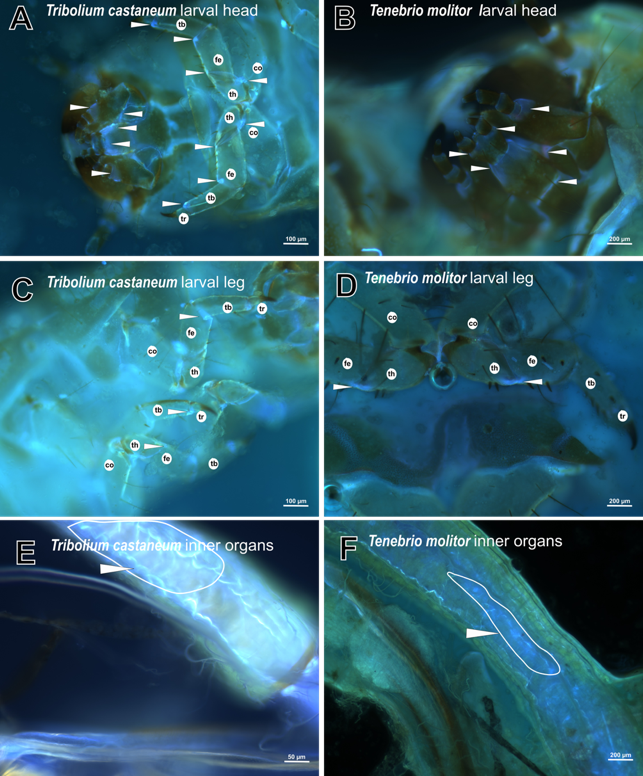

#### sfigure s14 – Larval samples of Tribolium castaneum and Tenebrio molitor.

Here, we show larval stages of *Tribolium castaneum* and *Tenebrio molitor* including inner organs (see Happ & Happ 1975 and Jaloszynski & Ruta 2021) for DT presence. (A, B) In the larval head and legs (C, D) of *Tribolium castaneum* and *Tenebrio molitor* DT is measurable. Both larvae are more transparent than adults; this allows detection of some signals in different locations. (E and F) In the inner organs DT specific signals are detected. The DT areas are pointed to with white arrowheads. The underlying measurements are shown in sfigure s15.

Images are done by an inverse Axio Observer Z1 (Zeiss) microscope with a black and white sensitive camera (Axiocam Mono).

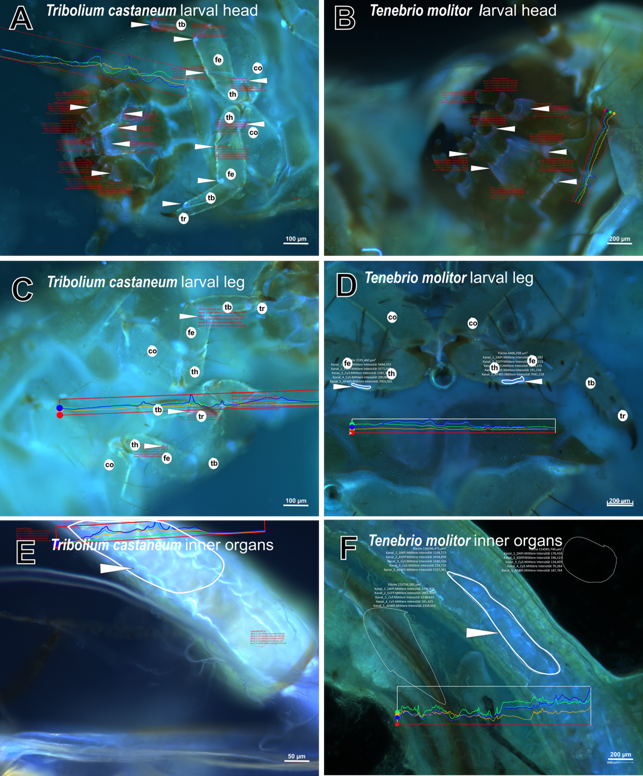

#### sfigure s15 - Data of larval samples of Tribolium castaneum and Tenebrio molitor.

This image is a copy of sfigure s14 including, in addition, areas of the measurement generated with the profile tool (by ZenBlue) and drawn by hand to include the DT signals.

The DAPI and the DT filters are named in German by the software: “Kanal_1_DAPI. Mittlere Intensität” and “Kanal_5_AF405.Mittlere Intensität”, respectively.

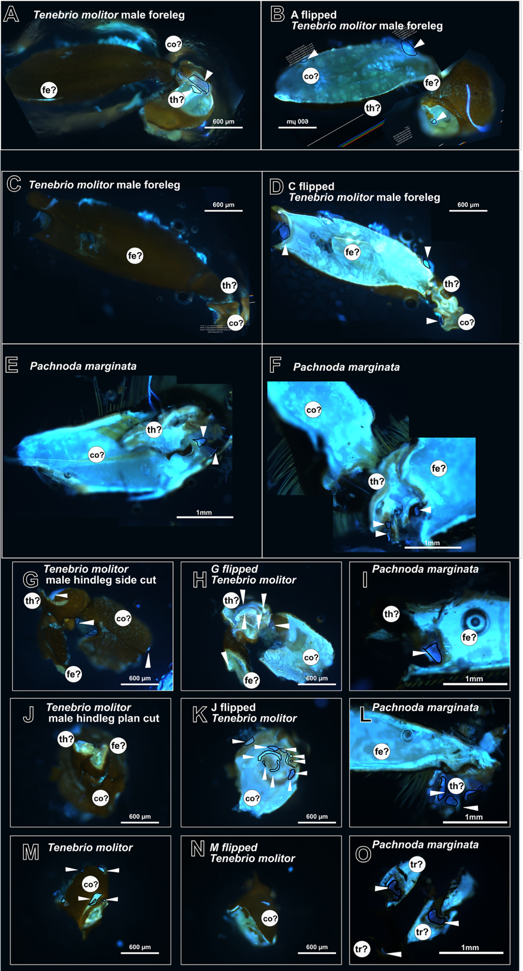

#### sfigure s16 – Dissection of Coleoptera samples.

Following information of different authors (like Nadein and Betz 2016 and 2018 or Burrows and Sutton 2012) that the dark stiff cuticle could mask DT, samples by *Pachnoda marginata* (E, F, I, L and O) and *Tenebrio molitor* (A-C, G, H, J, K, M, N) were dissected and opened.

The inner structures show a high auto-fluorescence also covering blue (mostly DAPI) and present areas where the DT-to-DAPI ratio is over 500 and 1000 counts. DT areas are framed with a black line and pointed to with a white arrowhead. Possibly because of the dissection process, some areas are deformed, or false-false areas occurred. Alternatively, this signal may emanate from inner structures formed by the soft membrane cuticle of the legs.

In general, a common shape or pattern is not visible, but DT is present.

The measured DT signals are pointed to with white arrowheads. The underlying measurements are shown in sfigure s17. All images were observed by an inverse Axio Observer Z1 (Zeiss) with an Axiocam Mono camera in false coloured images.

co?... suspected coxa, fe?... suspected femur, th?... suspected trochanter, tr... suspected tarsi

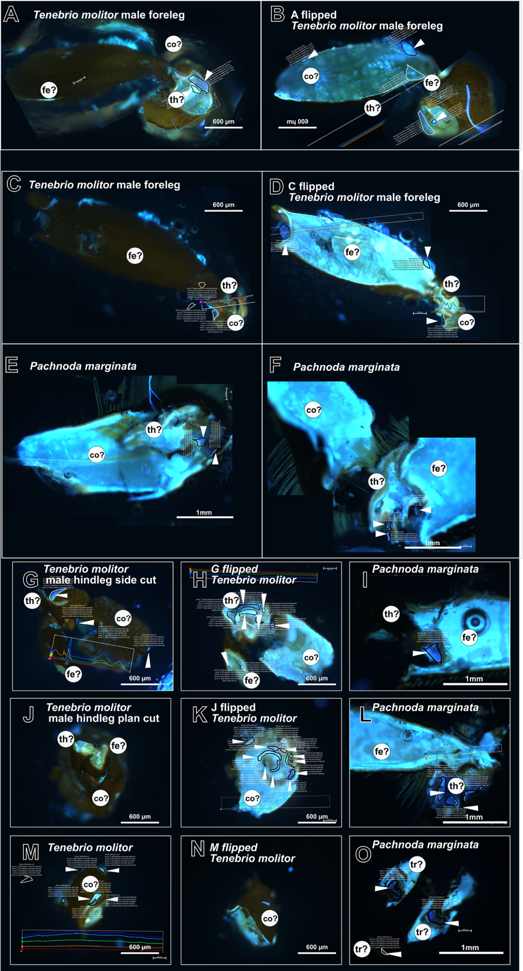

#### sfigure s17 - Data of Coleoptera sample dissections.

This image is a copy of sfigure s16 including, in addition, the areas of the measurement generated with the profile tool (by ZenBlue) and drawn by hand to include the Dityrosine signals.

The DAPI and the DT filters are named in German by the software: “Kanal_1_DAPI. Mittlere Intensität” and “Kanal_5_AF405.Mittlere Intensität”, respectively.

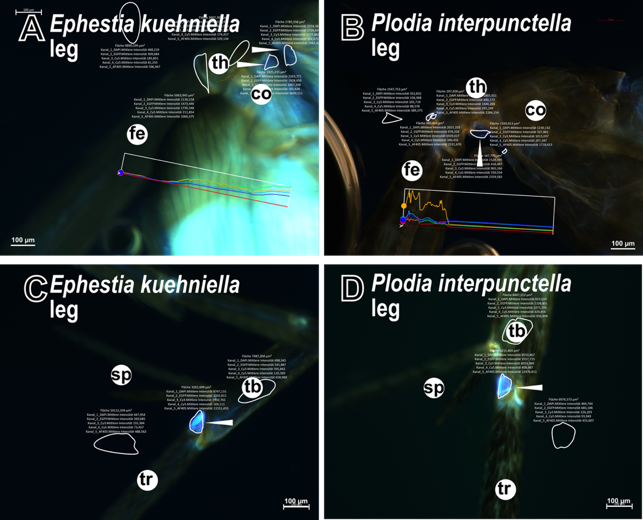

#### sfigure s18 - Overview of the Ephestia kuehniella and Plodia interpunctella data.

This image is a copy of figure 13 including, in addition, the areas of the measurement generated with the profile tool (by ZenBlue) and drawn by hand to include the DT signals.

The DAPI and the DT filters are named in German by the software: “Kanal_1_DAPI. Mittlere Intensität” and “Kanal_5_AF405.Mittlere Intensität”, respectively.

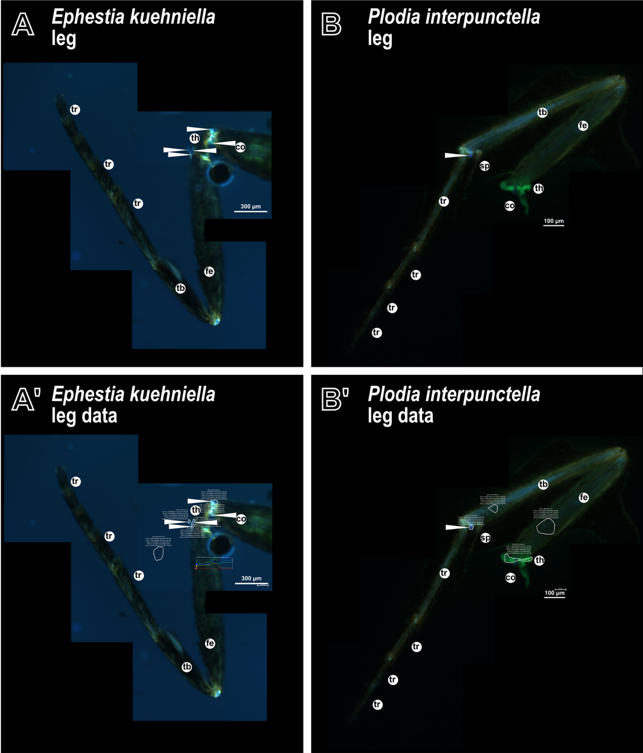

#### sfigure s19 – Whole leg of Ephestia kuehniella and Plodia interpunctella.

We present the whole leg of both moths of (A and A') *Ephestia kuehniella* and (B and B') *Plodia interpuctella*. The results of the measurements are shown in (A' and B').

(A and A') Inside the leg (opened by cracking), a DT signal in the coxa-trochanter region is detected. In this sample, the signal of tibia and tarsi is not visible. (B and B') In the *P. interpunctella* leg, the coxa and partly the trochanter are missing. The tibia and tarsi joints show a DT signal. For all samples, the removal of the scales from the surface was necessary.

The DT locations are pointed to with white arrowheads.

The images are composed of false coloured images generated with the software ZenBlue using an inverse Axio Observer Z1 (Zeiss) microscope with a camera (Axiocam Mono camera) and gained with the same settings.

co... coxa, fe... femur, sp... spur, tb... tibia, th... trochanter, tr... tarsi

The DAPI and the DT filters are named in German by the software: “Kanal_1_DAPI. Mittlere Intensität” and “Kanal_5_AF405.Mittlere Intensität”, respectively.

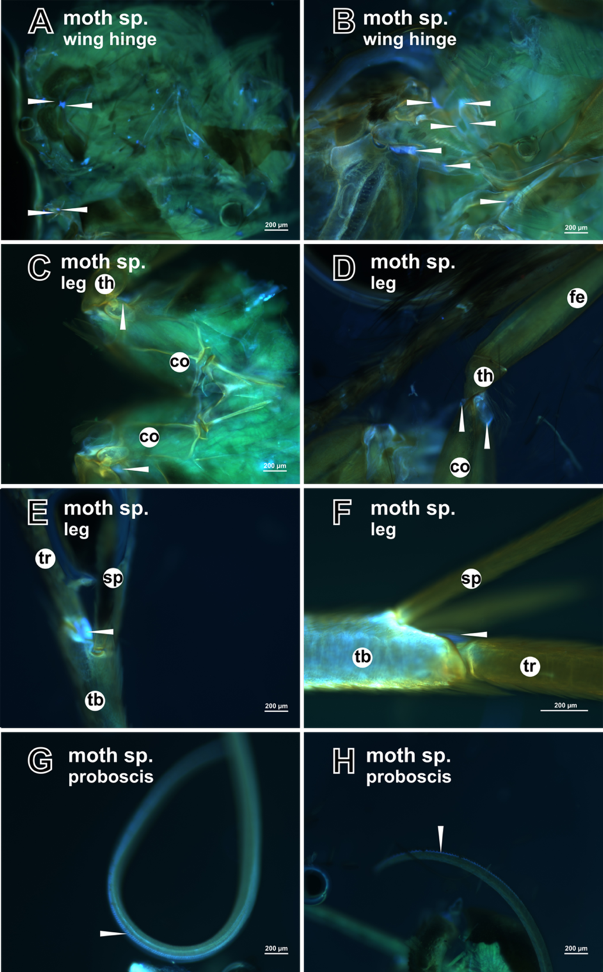

#### sfigure s20 – Lepidoptera samples.

Other moth species than *Epheastia kuehniella* and *Plodia interpunctella* were analysed, and some of these samples are shown in this figure.

(A and B) In these unidentified moth species, we detected different DT signals. (C and D) In comparison to other insects, no DT is detected in the trochanter, but the coxa to trochanter area contains DT. (E and F) An area between tibia and tarsi shows DT signals, it is unclear whether they constitute a structure like an antenna cleaner. (G and H) The proboscis of moths in general contains DT.

For all samples, the removal of the scales from the surface was necessary.

The DT locations are marked with white arrowheads. The underlying measurements are shown in sfigure s21. The images are composed of false coloured images produced by the software ZenBlue using an inverse Axio Observer Z1 (Zeiss) microscope with a camera (Axiocam Mono camera) and gained with the same settings.

co... coxa, fe... femur, sp... spur, tb... tibia, th... trochanter, tr... tarsi

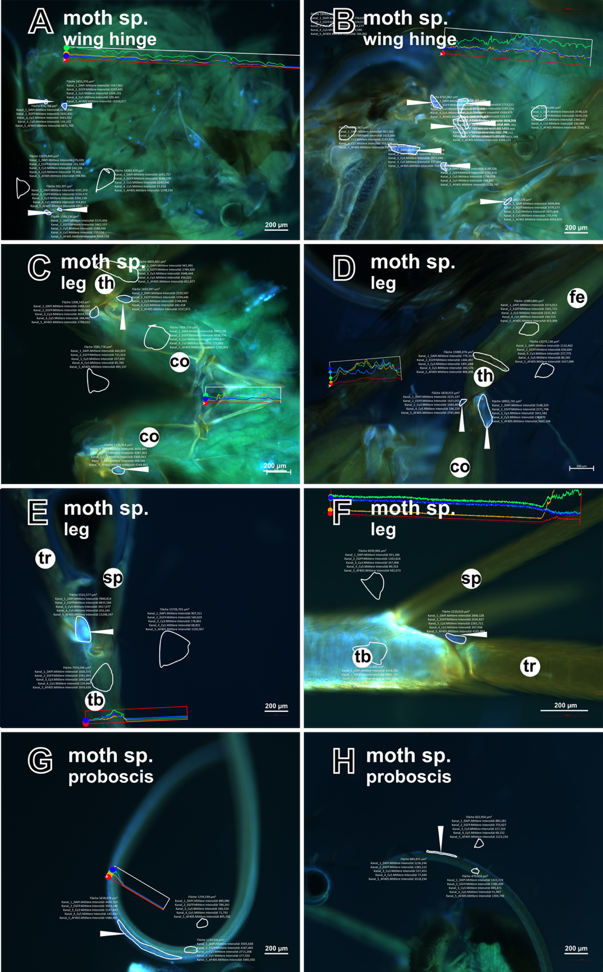

#### sfigure s21 – Lepidoptera sample data.

This image is a copy of sfigure s20 including, in addition, the areas of the measurement generated with the profile tool (by ZenBlue) and drawn by hand to include the Dityrosine signals.

The DAPI and the DT filters are named in German by the software: “Kanal_1_DAPI. Mittlere Intensität” and “Kanal_5_AF405.Mittlere Intensität”, respectively.

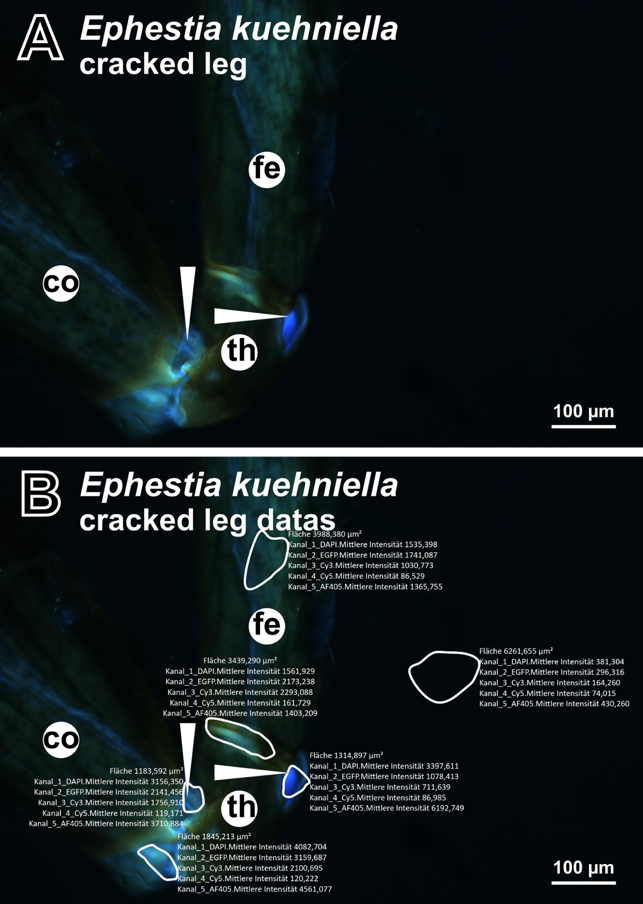

#### sfigure s22 – Dissection of Ephestia kuehniella.

Lepidoptera did not show any DT signal in the trochanter using our detection method when viewed from the outside. (A) In a first approach, we opened the respective area and found in the inside of the trochanter joint a DT signal. (B) The measured area is framed by a white line and the measurement results as total counts of the area is given to the frame. This shows that the signal in the upper area in the coxa to trochanter has a DT-to-DAPI value of over 500 counts, while the opened trochanter to femur area has a DT-to-DAPI value of over 1000 counts.

The removal of the scales from the surface was necessary.

All DT areas are pointed to by white arrowheads. The images were done by the inverse Axio Observer Z1 (Zeiss) microscope and the Axiocam Mono camera.

co... coxa, fe... femur, th... trochanter

The DAPI and the DT filters are named in German by the software: “Kanal_1_DAPI. Mittlere Intensität” and “Kanal_5_AF405.Mittlere Intensität”, respectively.

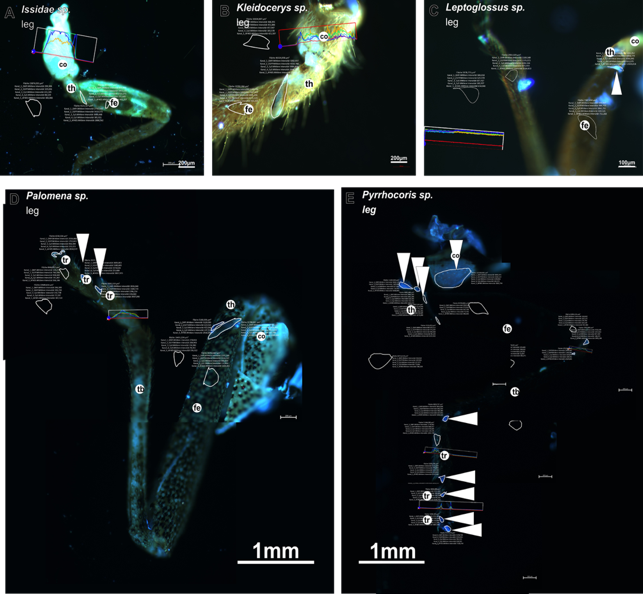

#### sfigure s23 -Hemiptera sample data.

This image is a copy of figure 14 including, in addition, the areas of the measurement generated with the profile tool (by ZenBlue) and drawn by hand to include the Dityrosine signals.

The DAPI and the DT filters are named in German by the software: “Kanal_1_DAPI. Mittlere Intensität” and “Kanal_5_AF405.Mittlere Intensität”, respectively.

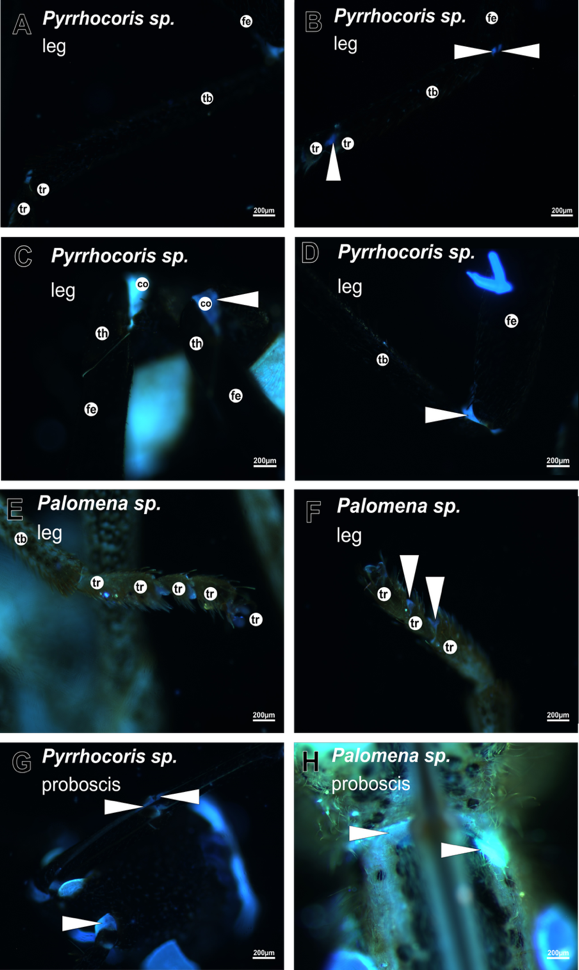

#### sfigure s24 – DT detection in Hemiptera.

In Hemiptera, DT detection in the cuticle outside the wing hinge is sporadic. That is, the fluorescence signal is probably dimming by the tanning degree of the cuticle.

DT is detected in the same leg area in *Pyrrhocoris sp.* only in the sample shown in (B) but not in (A). In (C) the *Pyrrhocoris sp.* sample of the same animal is sporadic. (D) shows a femur and tibia joint signal. In *Palomena sp.* (E and F), the tarsal DT signals are variable.

Whether the head i.e. proboscis signals (G and H) are every time visible, is unpredictable.

DT containing locations are pointed to with white arrowheads. The underlying measurement data are shown in sfigure s25.

An inverse Axio Observer Z1 (Zeiss) microscope with a black and white sensitive camera (Axiocam Mono) were used for the imagine.

co... coxa, fe... femur, tb... tibia, th... trochanter, tr... tarsi

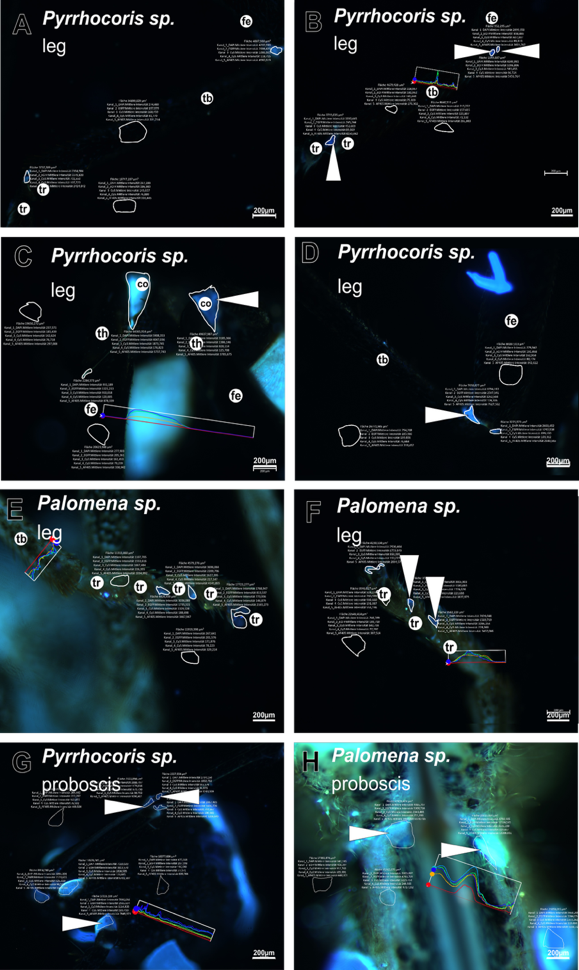

#### sfigure s25 – Data of DT detection in Hemiptera.

This image is a copy of sfigure s24, including, in addition, areas of the measurement generated with the profile tool (by ZenBlue) and drawn by hand to include the Dityrosine signals.

The DAPI and the DT filters are named in German by the software: “Kanal_1_DAPI. Mittlere Intensität” and “Kanal_5_AF405.Mittlere Intensität”, respectively.

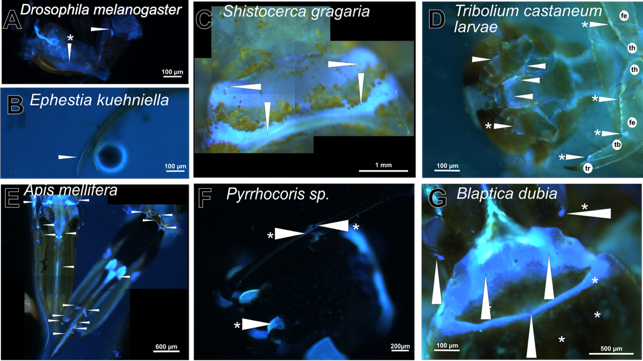

#### sfigure s26 Compilation of all orders - head

The insect head and feeding apparatus are divers. The variety is reflected by the DT distribution. In general, a unique localization pattern for each species is shown. With the applied method, there are no consistent DT signals in Coleoptera heads except for *T. molitor* and *T. castaneum* larvae and the adult of *Cantharis sp.*; in Hemiptera, DT localization in the head is not replicable, neither.

(A) The proboscis of *D. melanogaster* (from sfigure s28 and s29) has a strong DT signal in the labellum and a weaker one in the cibarium. (B) The proboscis of *E. kuehniella* has a strong DT signal (DAPI-DT filter, from sfig. s41 and s42), which is consistent with the assumption of Hepburn (1971). (C) The head of *S. gregaria* (from sfig. s32 and s33) has different strong DT patches; here, we show the upper head region with its intense bandlike areas. (D) In *T. castaneum*, head signals were detected only in larvae (from sfig. s14 and s15). Typically, in adult heads, no signals were detected (E). The honeybee proboscis was suggested by Rehder (1987) and Groneberg et al. (1993; 1997) to contain Resilin (from sfig.s30 and s31). (F) As an example for inconsistent DT signals in *Pyrrocoris sp.* (Hemiptera), the DT signals were always weak and occasionally fall below the DAPI-DT filter 500 count threshold (from sfig. s24 and s25). (G) The head of *B. dubia* reveals mostly strong DT signals (from sfig. s7 and s8), which resemble the pattern in Orthoptera samples.

The white arrowhead points to the DT signal, the white asterisk marks the areas of weak signals between 500 and under 1000 counts detected with the DAPI and DT filters. The images were obtained with the inverse Axio Observer Z1 (Zeiss) microscope and the Axiocam Mono camera.

co... coxa, fe... femur, tb... tibia, th... trochanter, tr... tarsi

The DAPI and the DT filters are named in German by the software: “Kanal_1_DAPI. Mittlere Intensität” and “Kanal_5_AF405.Mittlere Intensität”, respectively.

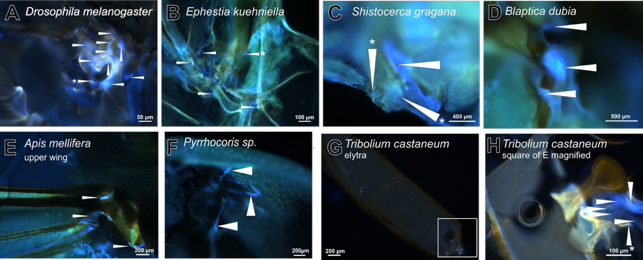

#### sfigure s27 compilation of all orders - wing hinge.

The insect wing hinge shows DT incidences in all analyzed samples. The numbers and intensities between the species are, however, different. A correlation of this information to the flight system needs further studies. By trend, smaller insects like Diptera, Lepidoptera and some Coleoptera species (like *T. castaneum*) showed rather dot-like signals, while bigger samples like Hemiptera, Orthoptera, Blattodea, Hymenoptera and bigger Coleoptera (*P. Marginata*) have a more coherent composition of DT signals.

(A) The wing hinge of *D. melanogaster* (from sfigure s34 and s35) has several signal patches with an overall intensity over 1000; these locations overlap with Res-GFP expression (Lerch *et al.* 2020). (B) There are several strong dot-like signals in the wing hinges of both *E. kuehniella* wings (from sfig. s41 and s42)*.* (C) In *S. gregaria,* DT is detected in patches with different shapes, we found DT signals also in the wing blade (from sfig. s32 and s33). These areas resemble those identified in the *L. migratoria* wing blade (Michels and Gorb 2012; Kovalev *et al.* 2018). (D) The wing hinge of *B. dubia* shows differently shaped DT signals with strong intensities (from sfig. s7 and s8). (E) DT signals in the honeybee wing have been described previously (Ma *et al.* 2015) and also by Nachtigall (*et* *a**l**.* 1998). We recapitulate these DT-positive areas and show DT signals in the upper wing hinge (from sfig.s30 and s31). (F) The Hemiptera *Pyrrocoris sp.* has strong, mostly small band-like DT signals in the wing hinge (from sfig. s43 and s44). (G, H) The hinge of the elytra of *T. castaneum* contains DT areas (from sfig. s38 and s39).

The white arrowhead points to the DT signal, the white asterisk marks the of weak signals (DT-to-DAPI difference of below 500). The images were obtained with the inverse Axio Observer Z1 (Zeiss) microscope and the Axiocam Mono camera.

co... coxa, fe... femur, tb... tibia, th... trochanter, tr... tarsi

The DAPI and the DT filters are named in German by the software: “Kanal_1_DAPI. Mittlere Intensität” and “Kanal_5_AF405.Mittlere Intensität”, respectively.

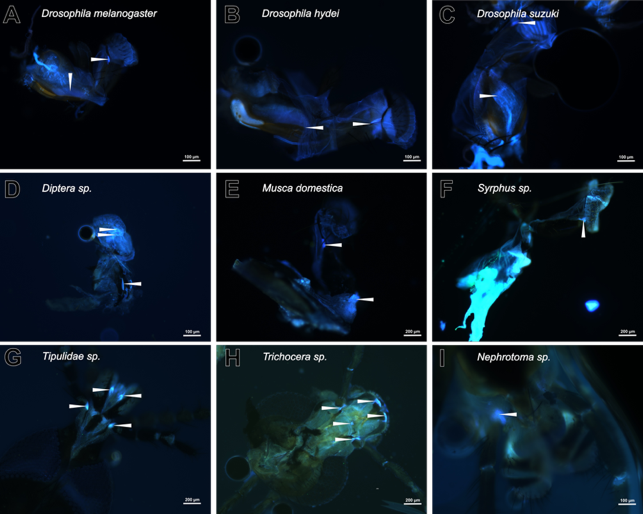

#### sfigure s28 -Diptera proboscis.

DT signal in the labrum and labellum in (A) *D. melanogaster*, (B) *D. hydei* and (C) *D. suzuki*. In flies with similar morphology, these signals are detectable in (D) *Diptera sp.* and (E) *Musca domestica*. In (H) *Syrphus sp.* the signal is very strong. In all *Drosophila* species the relation of the stronger labellum signal to the weaker labrum signal is shown. Other Diptera (G-I) do have a DT signal in the proboscis, but their morphology is different.

The white arrowheads point to the DT containing locations. The underlying data are shown in sfigure s29. The images were obtained with the inverse Axio Observer Z1 (Zeiss) microscope and the Axiocam Mono camera.

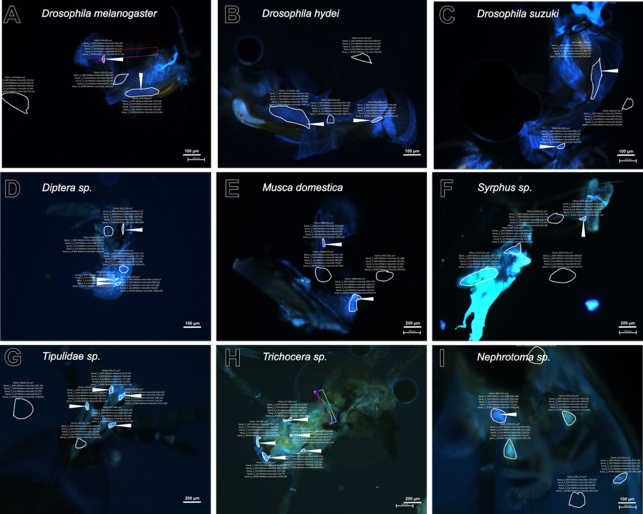

#### sfigure s29 -Diptera proboscis data.

This image is a copy of sfigure s28 including, in addition, the areas of the measurement generated with the profile tool (by ZenBlue) and drawn by hand to include the DT signals.

The DAPI and the DT filters are named in German by the software: “Kanal_1_DAPI. Mittlere Intensität” and “Kanal_5_AF405.Mittlere Intensität”, respectively.

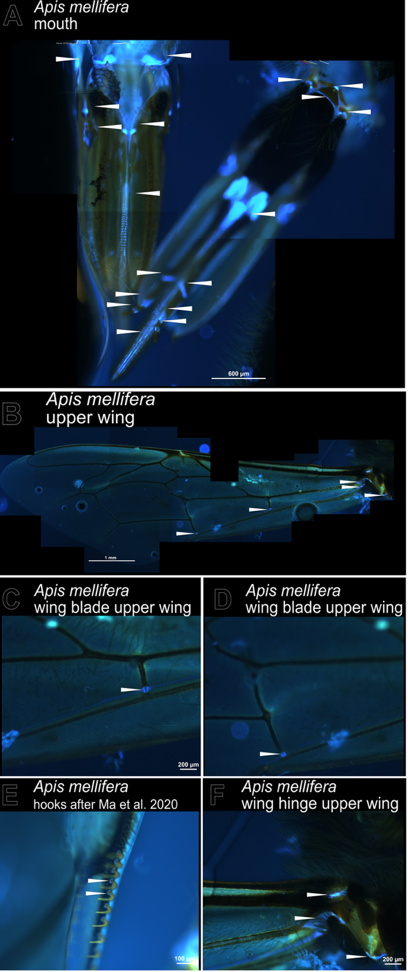

#### sfigure s30 -Apis mellifera mandible and wing

Prominent regions with DT in *A. mellifera* are the (A) mandible and (B) the wing. (B-E) In the anterior wing, we found DT signals in vein junctions. In C and D, we see closeups of DT in the veins (Ma *et al.* 2015). In E, we see that DT is present in the lateral hooks. (F) DT patches are present in the wing hinge of the upper wing (like Nachtigall *et al.* 2008).

White arrowheads point to the measured DT locations. The underlying raw data is shown in sfigure s31. The images were obtained with the inverse Axio Observer Z1 (Zeiss) microscope and the Axiocam Mono camera.

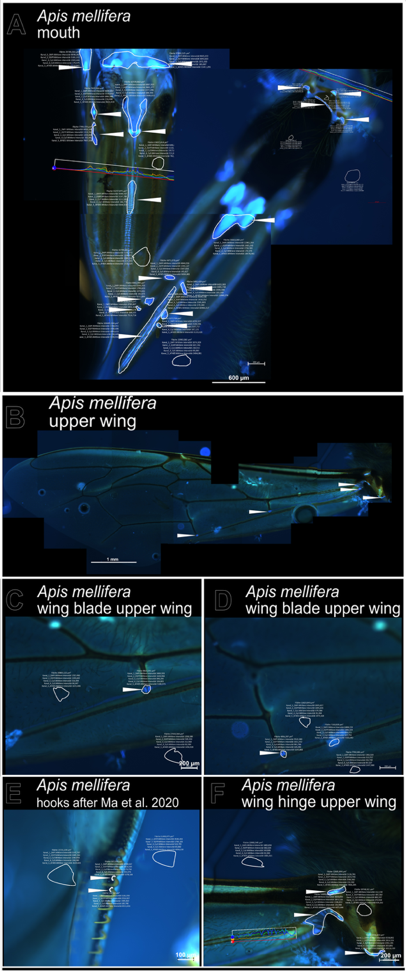

#### sfigure s31 -Apis mellifera mandible and wing data

This image is a copy of sfigure s30 including, in addition, areas of the measurement generated with the profile tool (by ZenBlue) and drawn by hand to include the DT signals.

The DAPI and the DT filters are named in German by the software: “Kanal_1_DAPI. Mittlere Intensität” and “Kanal_5_AF405.Mittlere Intensität”, respectively.

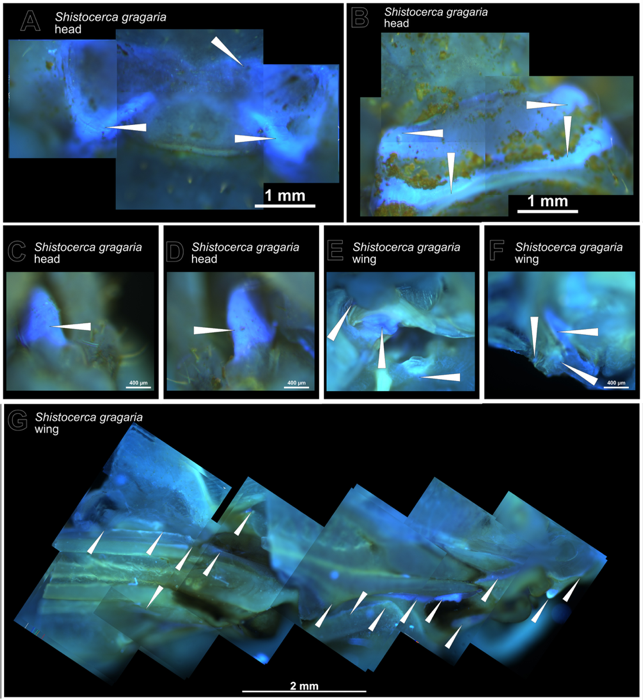

#### sfigure s32 – S. gregaria head and wing.

The *S. gregaria* shows Resilin in the (A-D) head (mandible) and the (E-G) wing. (A) The ventral side of the head shows two extended triangular shaped signals; the (B) dorsal head side has two broad DT lines. (C and D) On both lateral sides of the head, there is a tooth-like DT structure. (E and F) These images show different focused areas of the wing hinge. (G) The compilation of the cut wing close to the wing hinge shows that the wing blade contains DT.

White arrowheads point to DT containing locations. Underlying raw data is shown in sfigure s33. The imaging was done with an inverse Axio Observer Z1 (Zeiss) microscope and the Axiocam Mono camera.

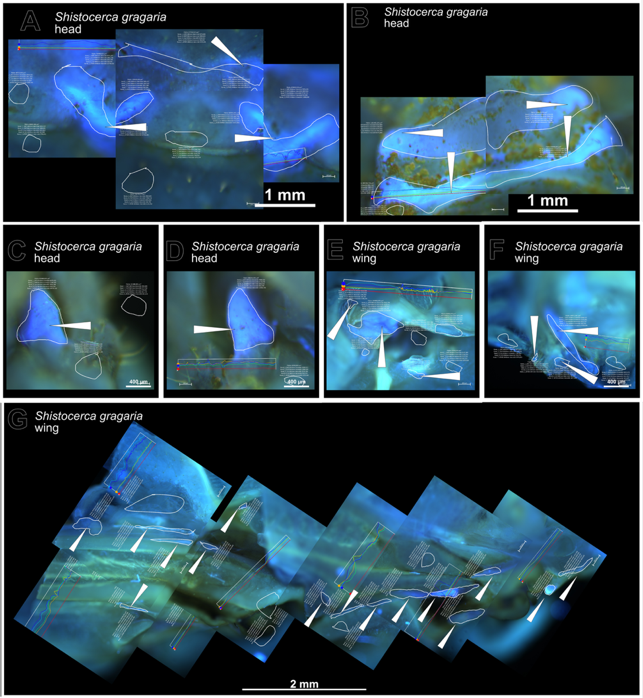

#### sfigure s33 -S. gregaria head and wing data.

This image is a copy of sfigure s32 including, in addition, the areas of the measurement generated with the profile tool (by ZenBlue) and drawn by hand to include the DT signals.

The DAPI and the DT filters are named in German by the software: “Kanal_1_DAPI. Mittlere Intensität” and “Kanal_5_AF405.Mittlere Intensität”, respectively.

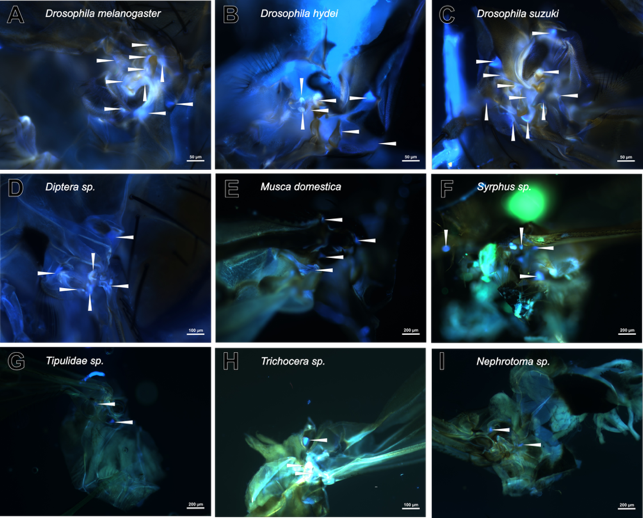

#### sfigure s34 -Diptera wing hinge.

The wing hinge of Diptera shows DT spots in all samples. The intensities vary in all samples and lie between 500 to over 1000 DT-DAPI counts. It seems that in some fly-related species including (A) *D. melanogaster*, (B) *D. hydei*, (C) *D. suzuki*, (D) *Diptera sp.*, (E) *Musca domestica* and (F) *Syrphus sp.* more DT dots are present than in the other *Nematocera*-related samples including (G) *Tipulidae sp.*, (H) *Trichocera sp.* and (I) *Nephrotoma sp.*

The white arrowheads point to the DT containing locations. Underlying raw data is shown in sfigure s35. Images are done by an inverse Axio Observer Z1 (Zeiss) microscope with a black and white sensitive camera (Axiocam Mono).

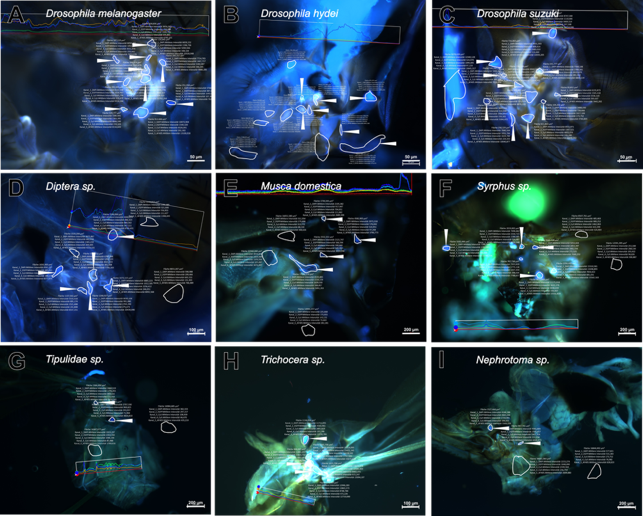

#### sfigure s35 -Diptera wing hinge data.

This image is a copy of sfigure s34 including, in addition, the areas of the measurement generated with the profile tool (by ZenBlue) and drawn by hand to include the DT signals.

The DAPI and the DT filters are named in German by the software: “Kanal_1_DAPI. Mittlere Intensität” and “Kanal_5_AF405.Mittlere Intensität”, respectively.

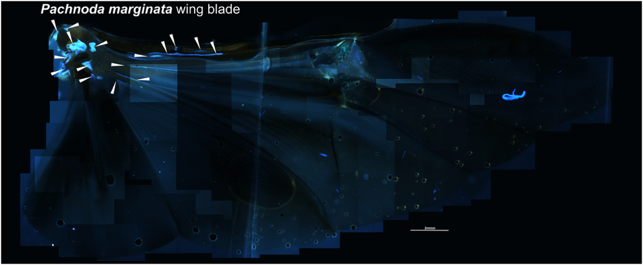

#### sfigure s36 - Pachnoda marginta wing blade.

The whole compilation of the wing blade to compare to the figures of Haas, Gorb and Blickhan (2000) shows similar locations of DT. Not all published locations were recapitulated, probably because of the different microscope settings and image quality. A recent publication of Appel et al. 2023 on the *P. marginata* wing blade, however, shows the same pattern as found in our work. Detailed images of spots with DT are in the sfigure s37. The DT containing areas are pointed to by white arrowheads. An inverse Axio Observer Z1 (Zeiss) microscope was used with a black and white sensitive camera (Axiocam Mono).

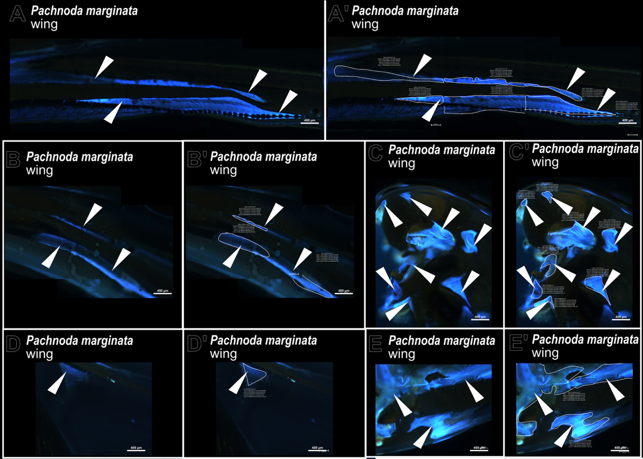

#### sfigure s37 - Pachnoda marginata wing sample data.

This image is a detailed composition of the wing blade signals of *Pachnoda marginata*. Images with dashed letters are those with added measurement areas generated with the profile tool (by ZenBlue) and drawn by hand to include the DT signals in comparison to the respective images with non-dashed letters. The white arrowheads point to the DT locations shown in sfigure s36.

The DAPI and the DT filters are named in German by the software: “Kanal_1_DAPI. Mittlere Intensität” and “Kanal_5_AF405.Mittlere Intensität”, respectively.

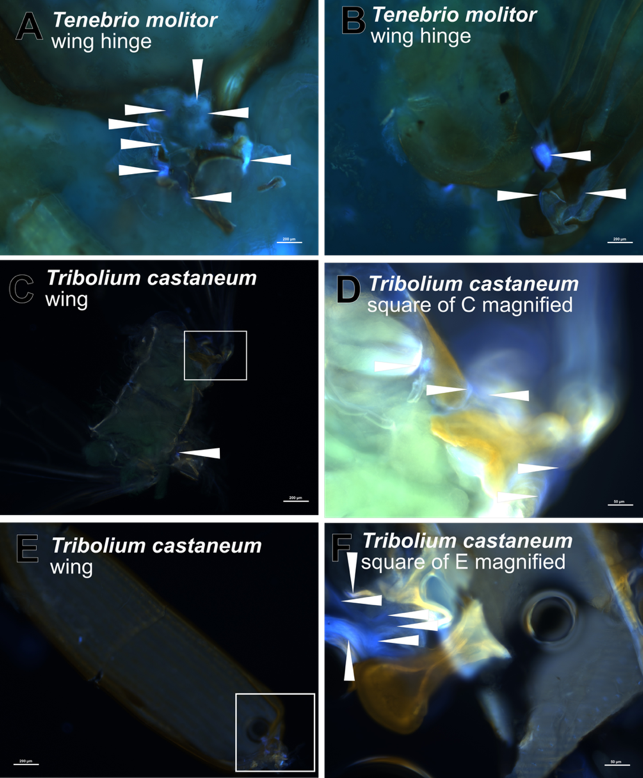

#### sfigure s38- Tenebrio molitor and Trobolium castaneum wing.

DT is detectable (A, B) in the wing hinge of *T. molitor* and (C-F) of *T. castaneum*. Higher magnification of the boxed area of the inner wing (C) and the elytra (E), are shown in D and F. White arrowheads point to the measured DT locations. Underlying raw data is shown in sfigure s39. For imaging, the inverse Axio Observer Z1 (Zeiss) microscope was used with a black and white sensitive camera (Axiocam Mono).

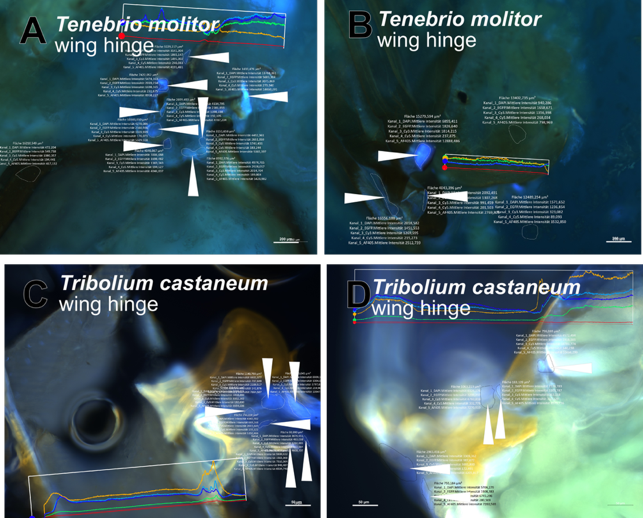

#### sfigure s39 – T. molitor and T. castaneum wing data.

This image is a copy of sfigure s38 including, in addition, the areas of the measurement generated with the profile tool (by ZenBlue) and drawn by hand to include the DT signals.

The DAPI and the DT filters are named in German by the software: “Kanal_1_DAPI. Mittlere Intensität” and “Kanal_5_AF405.Mittlere Intensität”, respectively.

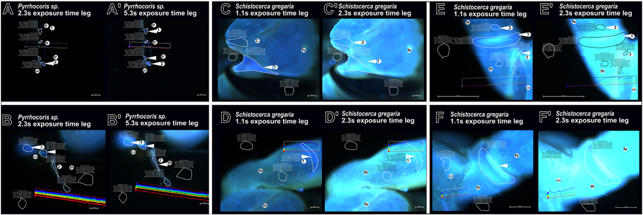

#### sfigure s40 Exposure time.

To show the effect of the exposure time for the measurement of DT or DAPI signal intensity, here, we analyse two samples. (A and A') In the *Pyrrhocoris sp.* leg, we observe two effects: a signal spot (1) changed the status from a no-signal state of 300 DT-to-DAPI difference counts at 2.3s exposure time to a signal state of 800 counts with 5.3s exposure time. A stronger signal (2) at the lower exposure time is also a signal at the longer exposure time (differences of 790 counts at 2.3s to 1400 counts at 5.3s). The *Pyrrhocoris sp.* (B, B') shows strong signals at both exposure times in the coxa (1) (difference of 1100 counts at 2.3 to 2400 counts at 5.3s), while in the small area of the trochanter to femur zone (2), we observe a signal only at the strong exposure time (difference of 300 counts at 2.3s to 525 counts at 5.3s).

The triangle shape in the joint of the (C, C') *S. gregaria* jump leg described by Burrows and Sutton (2012) displays at both exposure times a valid signal. Signal values are inconsistent because of the effect of oversaturation of the DT signal (from 3100 counts at 1.1s to 3000 counts at 2.3s). By consequence, in other areas of the joint (2) a false positive signal could occur, or a false negative signal may rise depending on the exposure time (difference of 400 counts at 1.1s to 750 counts at 2.3s). Another reference in *S. gregaria* is the buckling area in the tibia, which appears as a strong signal at both exposure times (D, D'). The oversaturation effect occurs also in other areas such as the trochanter-femur joints of the (E, E’) midleg or of the (F, F') first leg.

The trochanter-to-femur signal ratio (1) is high and is reduced at the higher exposure time (difference of 3000 counts at 1.1s and 550 counts at 5.3s). Signal (2) in E and E' is almost unchanged despite of rising the exposure time (difference of 1100 counts at 1.1s to 1400 counts at 2.3s), while signal (3) vanishes by rising exposure time (difference of 920 counts to 17 counts). In F and F', the trochanter signal is masked by oversaturation in comparison between the exposure times (difference of 2900 counts at 1.1s, down to 300 counts at 5.3s). The oversaturation (see C' 1, E' 1 or F) is visible in the measured values; the technical upper limit of measurement occurs at around 16.000 intensity counts and is the reason for the reduced difference when both filters yield oversaturated signals. Because of the oversaturation problem, we recommend testing first imaging at lower exposure times, before using the maximum possible exposure time avoiding the oversaturation effect, accepting that one may lose weak signals as false negative results.

As described in the materials and methods section, the differences were calculated by DT-DAPI subtraction. We find that there is no linear correlation between intensity of the areas and the exposure time. The signals of B, B' (2), C, C' (2) and (3) and E, E' (2) occurred only in this image samples and where not replicable in other images of this kind and therefore do not occur in the results of this species.

The white arrowhead points to the DT signal. The images were obtained with the inverse Axio Observer Z1 (Zeiss) microscope and the Axiocam Mono camera.

co... coxa, fe... femur, tb... tibia, th... trochanter, tr... tarsi

The DAPI and the DT filters are named in German by the software: “Kanal_1_DAPI. Mittlere Intensität” and “Kanal_5_AF405.Mittlere Intensität”, respectively.

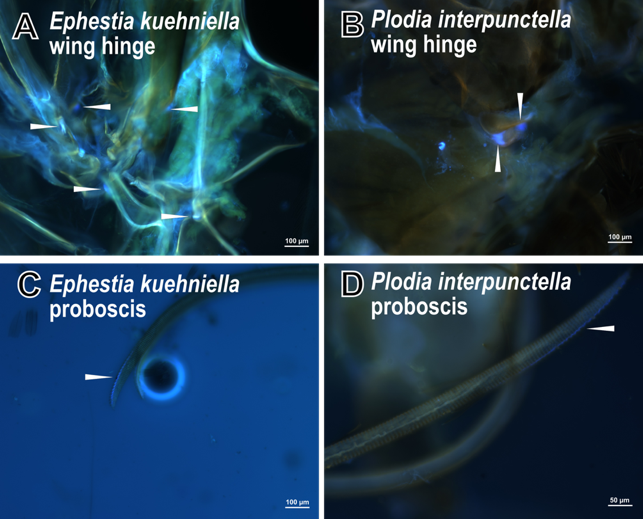

#### sfigure s41 - Overview of E. kuehniella and P. interpunctella wing and proboscis.

The storage pests (A, C, E, G) *E. kuehniella* and (B, D, F, H) *P. interpunctella* were the reference Lepidoptera species of this work.

Both species show DT in the same locations, like the (A and B) wing hinge and the (G and H) proboscis. No trochanter to femur signal is visible from the outside, but in the joint of coxa and trochanter, we find a signal (C and D). Another leg position with DT is the (E and F) tibia and tarsi connection.

White arrowheads point to the measured DT locations. Underlying raw data are shown in sfigure s42. The images were obtained with the inverse Axio Observer Z1 (Zeiss) microscope and the Axiocam Mono camera.

ac... antenna cleaner, co... coxa, fe... femur, sp... spur, tb... tibia, th... trochanter, tr... tarsi

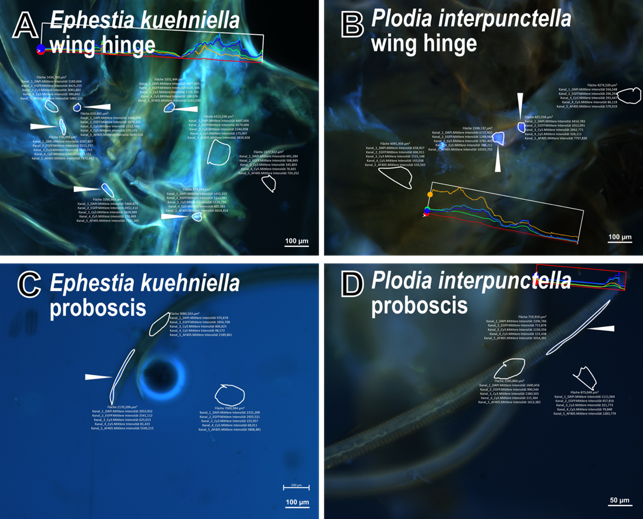

#### sfigure s42 - overview of Ephestia kuehniella and Plodia interpunctella wing and proboscis data.

This image is a copy of sfigure s41 including, in addition, the areas of the measurement generated with the profile tool (by ZenBlue) and drawn by hand to include the DT signals.

The DAPI and the DT filters are named in German by the software: “Kanal_1_DAPI. Mittlere Intensität” and “Kanal_5_AF405.Mittlere Intensität”, respectively.

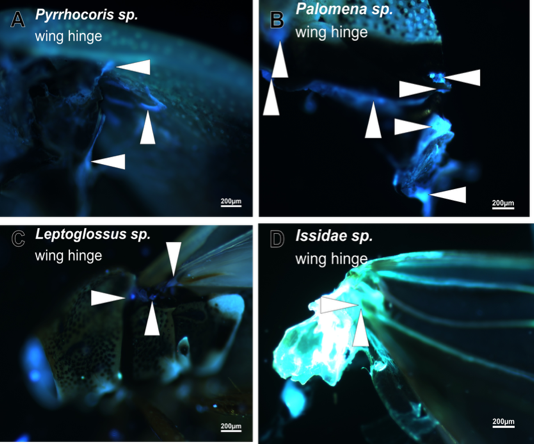

#### sfigure s43 - Hemiptera samples wing hinge.

In contrast to the sporadic findings in the legs, DT signals in the wing hinges are consistent. (A) *Pyrrhocoris sp.*, (B) *Leptoglossu specs.*, (C) *Palomena sp.* and (D) *Issidae sp.*.

White arrowheads point to the measured DT locations. Underlying raw data are shown in sfigure s44. The images were obtained with the inverse Axio Observer Z1 (Zeiss) microscope and the Axiocam Mono camera.

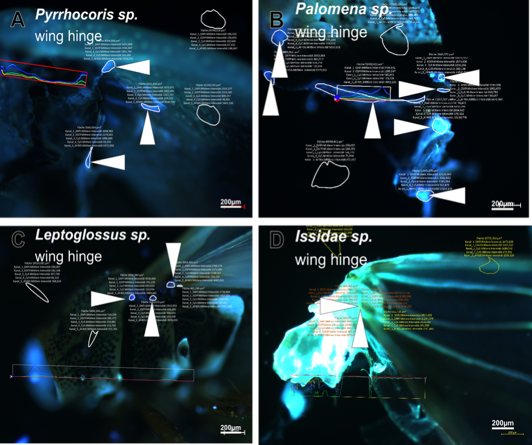

#### sfigure s44 - Hemiptera samples wing hinge data.

This image is a copy of sfigure s43 including, in addition, the areas of the measurement generated with the profile tool (by ZenBlue) and drawn by hand to include the DT signals.

The DAPI and the DT filters are named in German by the software: “Kanal_1_DAPI. Mittlere Intensität” and “Kanal_5_AF405.Mittlere Intensität”, respectively.

**Supplementary Results**

#### Resilin in different leg segments and in specific leg adaptations

We observed a complex pattern of Resilin-GFP and DT signal in the legs of the fruit fly *D. melanogaster*. This genetic method showed Resilin content in all joints, but we should note that DT and Resilin-GFP signal intensities were not proportional. In the following, we are presenting and discussing data not included in the main text.

##### Jumping apparatus

The morphology of the insect leg and its implication especially in jumping is well-studied. Years of research focused on the source of the jump power and the underlying mechanism in fleas or locusts. Bennet-Clark suggested the involvement of Resilin in the jump mechanism of the hindleg in fleas (Bennet-Clark and Lucey 1967). Initial investigations by fluorescence microscopy failed to substantiate this assumption (Heitler 1977). Rentz suggested, without actually showing it, that all Orthoptera have Resilin in their jump legs (Rentz 1996). In general, in early cases, the presence of Resilin was mentioned but not documented, for example in the hindleg of froghoppers (Burrows 2003) or *Sipyloidea sp.* (Burows and Morris 2002). Burrows and colleagues demonstrated first that Resilin is an important element in the jump apparatus of *S. gregaria* by fluorescence microscopy (Bayley *e**t* *al**.* 2012; Burrows & Sutton 2012). Thereafter, Bennet-Clark showed the energy storage capability in the semi-lunar process (Bennet-Clark 1975). They were able to show Resilin in the semi-lunar process element of the jump leg and a triangle patch of Resilin in the femur to tibia joint (Burrows & Sutton 2012).

Another area of interest is on the tibia described by Bayley et al. (Bayley *e**t* *al**.* 2012) and called “buckling region” visible as a stripe. Burrows showed later that this region is connected by a Resilin-containing inner tendon of the tibia muscle (Burrows 2016). The influence of Resilin was also demonstrated recently in a genetic approach (Rogers *et al*. 2025).

For Blattodea, Resilin was found in the joints of tibia of *Periplaneta americana* (Neff *et al**.* 2000) and in the jump mechanism of *Saltoblattella montistabularis* (Picker *e**t* *al**.* 2012). Resilin in leg segments has been reported in different hymenopteran species including ants and bees (Federle *et al.*, 2001) and wasps (Frantevich & Gorb 2002; 2004), but these incidences have not been correlated with jumping.

Jumping Coleoptera were also suggested to have Resilin in their legs (Furth 1980), but later the authors rejected that idea based on missing evidence (Alticinae, Furth *et al.* 1983 and Furth 1988). Resilin was shown by opening the hindlegs of such Coleoptera species like *Sphaeroderma testaceum, Podagrica* *fuscicornis* (Alticinae, Nadein & Betz 2016) and *Orchestes fagi* (Cucujiformia, Nadein & Betz 2018), and in the inside of the femur-tibia joint of *Lethrus apterus* and *Pentodon idiota* (Scarabaeidae, Frantsevich *e**t* *a**l**.* 2019). Resilin was finally detected in jump apparatus of Coleoptera, too*.* The presence of Resilin in elastic parts of the legs of *P. marginta* (Busshardt *et* *a**l**.* 2012) or of the tenebrionid beetles (Ichikawa *e**t* *a**l**.* 2016) were suggested but were not demonstrated. By contrast, in the inside of tibia of *Carausius morosus* (Phasmatodea), Resilin was visualized, without, however, mentioning if its visible from the outside as well (Schmitt *et al.* 2018). In summary, the femur and tibia are differentially equipped with Resilin, conceivably reflecting its functional need in different insect species.

##### Tarsi and praetarsi

For the praetarsi of a number of insect species like *Chironomus riparius, Leptinotarsa decemlineata, Aeshna mixta; Sympecma annulata, Bembix rostrata, Notonecta glauca, Ranatra linearis, Graphosoma italicum, Leptinotarsadecemlineata* and *Lethrus apterus*, Resilin presence was suggested using electronic microscopy and staining with methylene blue (Gorb 1996). Later, Resilin in the tarsi and adhesive pads was observed in the Blattodea species of *Periplaneta americana* via DT signals (Frazier *et al*. 1999) and in the joints of tibia and tarsi (Neff *et al**.* 2000). In the Orthoptera species *Locusta migratoria* and *Tettigonia* viridissima, Resilin in the adhesive was reported (Perez Goodwyn *et al.* 2006). For Hemiptera, Resilin was revealed in praetarsi of *Dicyphus errans* (Voigt *et al.* 2007) or in attachment structures, adhesive pads and setae in *Nezara viridula*, *Coreus marginatus* or *Cimex lectularius* (Rebora *et al.* 2018; Reinhardt *et al.* 2019; Rebora *et al.* 2021a). In *Platycnemis pennipes* (Odonata), Resilin was shown in the praetarsi (Gorb 1996). In *Anax imperator*, it was shown that Resilin occurrence was dependent on age and tarsal cuticle tanning (Preuss *et al.* 2024). For Hymenoptera, Resilin was found in the pretarsi and tarsi as well as the antenna cleaner on the foreleg (Frantsevich & Gorb 2002; 2004 and Endlein & Federle 2008 and Beutel *e**t* *a**l**.* 2020). In Coleoptera Resilin occurred in small structures like the tarsal setae (Peisker *e**t* *a**l**.* 2013) and tarsomeres of female *Coccinella septempunctata* (Michels and Gorb 2012). The tarsi of *Ischnura elegans* (Odonata) used for grooming bearing hair bases containing Resilin (Piersanti *et al.* 2024). In *Mantophasma kudubergense* (Mantophasmatidae) (Eberhard *et al.* 2009), the DT auto-fluorescence was found in the praetarsi; and in *Sungaya aeta* (Phasmatodea), the age and water dependent fatigue of Resilin in the adhesive pads was analysed (Grote *et al.* 2024). Via genetic modification in *Bombyx mori* the role of Resilin as material for adhasive areas was demonstrated, revealing the influence and occourence in the arolium in this Lepidopteran species. That Diptera contain Resilin in their tarsi was demonstrated first in the praetarsi of *Calliphora erythrocephal* (Bauchhenß 1979)*,* the pulvilli of *Episyrphus balteatus* (Gorb 1998) between claws and adhesive pads and the tarsomere joints in *Musca domestica* and *Eristalis pertina* (Niederegger & Gorb 2003), *Braula coeca* (Büscher *et. al* 2022). Later, Resilin was detected in tarsal joints and praetarsi of *Eristalis tenax* (Michels and Gorb 2012). Resilin was mentioned in the tarsal segments of *Aedes albopictus* but not analysed in detail (Croce and Scolari 2022b; Scolari *et al.* 2022). the In the fruit fly *D. melanogaster*, Resilin incidence in the tarsi was shown (Lerch *et al.*, 2020), however, a detailed analysis was not presented. Taken together, tarsal Resilin is a common trait in insects; its tarsal localisation, however, varies with species.

##### Trochanter

There are four articles dealing with Resilin in the trochanter. Focusing on the trochanter, a band-like structure of Resilin-GFP was identified overlapping with a distinct DT signal (Lerch *et al.* 2020). This shape of DT auto-fluorescence was described also for *D. suzukii* and *D. hydei* (Lerch *et al.* 2022). Likewise, in *Gryllus firmus* (Hayot *et al.* 2013), Resilin was shown, however, in *P.* americana, presence of Resilin was refuted (Zill *et al.* 2000).

*Resilin in head and feeding apparatus.*

The feeding mechanism of insects is highly divers, so even between species of the same order it is difficult to compare this region. Early works on Resilin were, for example, published by Rice, who described the cuticle composition in Diptera (Rice 1970). He focussed on the feeding mechanism including the cibarium of tsetse flies applying histological methods for Resilin staining and fluorescence detection, and mechanics to highlight Resilin. In other publications, Resilin presence was suggested in the proboscis of *Rhodnius prolixus* (Hemiptera) or the labium of *Brevicoryne brassicae* (Hemiptera) (Bennet-Clark 1963; Wensler 1977), as well as in the food pumps of reduviid bugs (Hemiptera) in general (Edwards 1983). In *Bactrocera oleae* blue fluorescence was shown in the head, but Resilin was not mentioned for these areas (Rebora *et al.* 2021b). The mouthparts of *Haplothrips verbasci* larvae (Heming 1978) and the head region of adults (Chent *et al.* 2020) suggested to use Resilin in the folding mechanism. For Lepidoptera Hepburn (1971) detected a flexible region in the whole dorsal part of the cuticle wall of the sucking organ in *Adela viridella*, *Pieris brassicae*, *Danaus plexippus* and *Deilephila elpenor*, but did not provide any fluorescence images in this article (Hepburn 1971). The presence of Resilin in the honeybee proboscis was suggested by Rehder (1987); consistently, Groneberg et al. (1993; 1997) stated that ant mandible shells also contain Resilin.

The identification with fluorescence microscopy alone in the head started after 2010 with *Periplaneta americana* (Blattodea) including the maxillary palp, membranous areas of stipes, galea and lacinia (Schmitt *et al.* 2014). Büsse and colleges showed Resilin in the hunting tools of the whole mouth in larvae of *Anax imperator* (Odonata) (Büsse *et al.* 2021). The age dependent occurrence of Resilin was demonstrated in the mandible of *Anax imperator* (Preuss *et al.* 2024). For two species of Mantodea, *Sphodromantis lineola* and *Gongylus gongylodes,* Resilin was demonstrated in their mandibles (Roze *et al*. 2024). *Culex pipiens* and *Aedes albopictus* Resilin was found in the head and in the antenna (Croce and Scloari 2022 a; b, Scolari *et al.* 2022). Resilin was identified in *D. melanogaster*, *D. suzukii* and *D. hydei* in the labellum and cibarium (Lerch *et al.* 2020, 2022).

With the usage of MISM, we refer to the findings of Hepburn (1971), showing for Lepidoptera, especially for *P.* *interpunctella* and *E. kuehniella,* Resilin in the proboscis (sfigure s41 and s42). For Diptera like Drosophilidae (Lerch *et al.* 2020, 2022), we found a labellum and cibarium signal in all studied samples with a proboscis (see sfig. s28 and s29). In Lepidoptera, the proboscis signal was over 1000 counts, in Diptera, the proboscis signal exceeded the high threshold consistently only in the labellum, while their the cibarium signal passed the 500 counts threshold. This intensity difference in the proboscis was also described in Lerch (*et al.* 2020, 2022). The other Diptera samples with a different mouth structures revealed Resilin with high intensities (over 1000 counts), too. Positive signals for Resilin in the head were visualised in Hymenoptera with high values (see sfig. s10, s11, s30 and s31). The head of *S. gregaria* showed strong signals (over 1000 counts) with different shapes, like the band-like shapes in the dorsal head part or a tooth like signal on the lateral edge (see sfig. s32 and s33). These band-like shapes occur also in the *G. assimilis* samples (see sfig. s5 and s6). Similar shapes of Resilin were also found in the head of *B. dubia* (see sfig. s7 and s8). These data confirm Resilin in the head of Blattodea. Indeed, mouthparts of *P. americana* including Resilin have been described in detail (Schmitt *et al.* 2014), however, homologies of Resilin incidences between *B. dubia* and *P. americana* need to be analysed in detail.

In the heads of different Hemipteran species, we were unable to distinguish a clear DT signal that was significantly higher than the DAPI signal except for two samples (sfig. s24 and s25). The bending part of the proboscis is usually a confident structure to assume Resilin presence. In two specimens (*Pyrrhocoris* sp. and *Palomena* sp.), we found signals, which we could not repeat based on the inconsistency between samples (see s-table 1).

For Coleoptera adults, no signal was found except for *Cantharis sp.* (see figure 11 and sfig. s12), which were measured with values over 1000 counts. The larval heads of *T. molitor* and *T. castanaeum* contained Resilin with a range of 500 to over 1000 counts difference in different spots (see sfig. s14 and s15).

Compiling these measurements, the heads of insects with their variety of feeding mechanisms are an interesting organ to locate Resilin in order to analyse the mechanics of food uptake and processing involving extreme forces.

*Resilin in the wing blade and hinge*

The first data of Resilin in these structures were presented by Weis-Fogh (1960) in *Schistocerca gragaria* prelar arm and wing hinge and a dragonfly. Then, other authors first described the necessity of Resilin in other insects, too, for example, Miyan and Ewing (Miyan & Ewing 1985) who mentioned the elastic energy involving up- and downstroke of the wing in Diptera. Further Pfau (1987) discussed that Resilin elements may function during wing stroke. Resilin in wing hinges were assumed over the changes in cuticle flexibility by temperature in *Aedes aegypti* (Costello 1974) or *Chrysops beameri* (Drees 1980). Some authors claimed the presence of Resilin in the wing hinge of Diptera without showing Resilin at all (Wisser and Nachtigall 1984; Trimarchi and Schneiderman 1994; Dickinson 2003). Resilin detection by fluorescence microscopy was firstly described in the cross veins of the wings of the damselfly *Enallagma cyathigerum* (Gorb 1999). Later, detection of Resilin in the flight system of dragonflies and damselflies such as *Epiophlebia superstes*, *Sympetrum striolatum*, *S. vulgatum* and *Matrona basilaris* was extensively analysed (Donoughe *et al.* 2011; Appel and Gorb 2011; Michel and Gorb 2012; Rajabi *et al.* 2016a; b; Appell *et al.* 2015; Michels, Appell and Gorb 2016; Fauziyah *et al.* 2014). In the bullae of wings of mayflies, it was only shown for *Siphlonurus lacustris* (Domínguez *et al.* 2023). In other orders such as Hymenoptera (*Apis* mellifera, Ma *et al.* 2015; 2020; *Bombus impatiens*, (Nachtigall *e**t* *a**l**.* 1998; Mountcastle & Combes 2014) or Blattodea (*Periplaneta* americana, Neff *et al.* 2000), the presence of Resilin in the wing blade or the wing hinge has been reported, as well. Even other Orthoptera were shown to contain Resilin in the wing hinge and prelar arm: *Locusta migratoria* (Michels and Gorb 2012; Kovalev *et al.* 2018). Moreover, wing movement and folding were proposed to require Resilin in Coleoptera such as *Pachnoda marginta* and *Coccinella spetempunctata* (Haas 1994 and Haas *e**t* *a**l**.* 2000a). For *P. marginta*, it was also imaged (Haas *e**t* *a**l**.* 2000a). Li and colleagues showed Resilin also in the blade of the h and folding areas of the ind wing of the bamboo weevils, Cy*rtotrachelus buquet* (*C*oleoptera) (Li e*t* *a**l**.* *2*019). In his master thesis, Li reported the mRNA coding for pro-resilin in the wing of *Tribolium castaneum*, but failed to detect a DT-signal in the wing blade (Li 2013). In Dermaptera, Resilin was found in the wing blade needed for wing folding (*Forficula auricularia* Haas *et al.* 2000b; *Forficula auricularia* and *Labia minor* Deiters *et al.* 2014; 2016). Resilin has been mentioned in the basal sclerite of the forewing attachment site of *Eumaeus atala* without, however, actually documenting this localisation it was mentioned too (Koi and Daniels 2015). In an earlier publication, we visualized Resilin and Resilin-like proteins in the wing hinge in D. *melanogaster* and *D. suzukii* (Lerch e*t al.* *2*020, 2022). We also detected Resilin a small DT area in the wing blade of *D. hydei* (Lerch *et al.* 2020; 2022). An evolutionary context of Resilin was discussed in Haas (2006) or Donoughe (*et al.* 2011). Donughe and colleagues show the species-specific distribution of Resilin in the wing vein joints, Haas proposed to include Resilin incidences as a marker, but not as dispersal point of groups. A modern review showed the diversity and need of Resilin in wings in a variety of insect orders (Appel *et al.* 2023).

Applying MISM, we examined the presence of Resilin in the wing hinges and blades. In Diptera, especially for *D. melangaster, D. suzukii* and *D. hydei* (see sfigure s34 and s35), we were able to reproduce formerly detected Resilin spots (Lerch *et al.* 2020, 2022). To evaluate our microscopic approach, we referred to the results in *P. marginta* (Haas *e**t* *al.* 2000a). For the same areas, we allocate signals in the wing that show a Resilin specific intensity over both value thresholds: 500 and 1000 counts (see figure 10, sfig. s36 and sfig. s37). We were able to recapitulate the signals reported by Ma (*e**t* *a**l**.* 2015) ) for the wing vein areas of the fore and hind wings and the basal structure of the wing (see fig. 9, sfig. s30 and s31). However, the lateral stripes could not be recognized at a threshold over 500. As a different reference, we were also able to repeat the findings by Nachtigall (*et* *a**l**.* 1998) in the wing hinge. The counts of intensity in *A. mellifera* were in general over 1000, only the hooks were measured between 500 and 1000 counts. In addition, we provide new results of other samples. In cases of Diptera, Resilin is identified in several dot-like areas in the thorax. The only variability is the total amount of these areas, with on average 5 to 7 spots in Drosophilae, and the 2 to 4 spots in Tipulidae.

For *S. gregaria*, we found Resilin in the wing (see sfig. s32 and s33) and in *B. dubia* at the wing hinge (sfig. s7 and s8). The intensity variety found in *P. marginata* is also observed in all Diptera. Similarly, we found DT fluorescence in hinges of the elytra and wing in *T. molitor* and *T. castaneum* (sfig. s38 and s39). In Lepidoptera, especially in *P.* *interpunctella* and *E. kuehniella*, DT distribution in the wing hinges is as in other insect species described above (see sfig. s41 and s42). Finally, DT signals in the wing hinges in Hemiptera are comparable to those in other species (see sfig. s43 and s44).

Together, in all wing hinges analysed here, DT is detectable. For the wing blade, DT incidences depend on the size of the insect. In many cases, bending and folding of wing blades possible necessitate DT at crucial points. By consequence, the intensity and shape of the DT areas varies between species.

*Other body parts of insects with Resilin*

After the first identification of Resilin as a protein matrix (Weis-Fogh 1961a; b; Andersen 1964; Andersen and Weis-Fogh 1964; Jensen and Weis-Fogh 1962) Resilin was studied in all kinds of body elements. In the following, we give a brief summary of these studies:

In a seminal work, it was shown that Resilin was deposited in the locust tibial cuticle following the circadian rhythm (Neville 1963a; b; 1967). Resilin in insect genitals was suggested in ovipostor of thripiden (Bode 1975) or *Iridophora clarki* (Disney 1986). Applying imaging and molecular techniques, Resilin was detected in the ovipositor in a number of different hymenopteran species: Eupleminae (Gibson 1986), *Apis mellifera* (Hermann and Willer 1986), *Cr**yptocheilus versicolor* (Kumpanenko and Gladun 2018), *Leptopilina heterotoma (*Sampalla *e**t* *a**l**.* 2018), *Diachasmimorpha longicaudata* (Cerkvenik 2019), *Vespa crabro* (Stetsun and Matushkina 2020) and in Aculeata: *Evania sp., Epyris sp., Vespula germanica, Dasypoda hirtipes, Bombus terrestris, Cryptocheilus versicolor, Sapyga similis, Scolia sexmaculata* and *Mutilla europaea* (Kumpanenko *et al.* 2019). In the literature, it was also claimed that the spermatheca of *Hebrus pusillus* consisted of Resilin layers (Heming-van Battum and Heming 1986). Ilango (2005) postulated the presence of Resilin in spermatheca in *Drosophila*. We confirmed this with GFP-tagged Pro-Resilin (Lerch *et al.* 2020). In agreement with this, for Coleoptera, Conti (*e**t* *a**l.* 1972) showed Resilin in the tubulus of spermatheca in *Dytiscus margin**alis*. Moreover, spermathecal ducts of the mealworm beetle *T. molitor* emitted UV-excited blue signal indicative of Resilin (Happ and Happ 1975). However, (Frenk and Happ 1976) did not detect any Resilin signal in the spermatophores of *T. molitor*. Even in smaller moveable context DT of Resilin was spotted like in the flagellum and the genital wall of *Cassida rubiginos* (Filippov *e**t* *a**l**.* 2016; Matsumura *et* *al.* 2017*)* or in the sperm pump of *Monotoma picipes* (Jaloszynski and Ruta 2021). In the cuticle segments of the penis in *Brachythemis lacustris* (Miller 1982), presence of Resilin was suggested; later Miller postulated it for the whole group of Libellulidae in the Odonata (Miller 1991). In Lepidoptera, the presence of Resilin was suggested in the genital region and pheromone secretion system without, however, providing fluorescence microscopy images (Wunderer 1990; Egelhaaf *e**t* *a**l**.* 1992). An overview, hence, cannot be drawn here. Michels and colleagues identified Resilin in the spermalege of female *Cimex lectularius* (Michels *et al.* 2015).

Resilin is also found in the lens cuticle of the firefly *Photinus pyralis* (Sannasi 1970), in *Mesogomphus lineatus* (Viswanathan and Varadaraj 1985) or in *Monomorium pharaonis* and *Chrysoperla carnea* (Michels and Gorb 2012).

The occurrence of Resilin was often suggested in sensilla in a variety of insect species like in *Apis mellifera* (Thurm 1964), Thripids (Bode 1975), *Tetrastichus hagenowii* (Barlin *et al.* 1981), *Calliphora vicina* (Grünert and Gnatzy 1987) or *Rhagoletis pomonella* (Stoffolano *et al.* 1987). In *Eristalis tenax* and *Acheta domesticus* (Michels and Gorb 2012; Michels, Appel and Gorb 2016), the Resilin content of sensilla were demonstrated by confocal microscopy.

That sound organs (tymbals) use Resilin was shown for cicada such as *Agalmatium bilobum* (Young and Bennet-Clark 1995). For tiger moths and woolly bears, Resilin was mentioned in the tymbal organ by referring to non-published personal communication with Frank Coro (Conner 2009).

In Siphonaptera and Hemiptera, Resilin was found in the pleural arch that is needed for a jump mechanism located in the thorax.First description based on images with molecular staining of Resilin was done for Siphonaptera: *Spilopsyllus cuniculus* (Bennet-Clark and Lucey 1967), for *Xenopsylla cheopis* (Rothshild 1975; Rotschild and Schlein 1975; Rothschild *et al.* 1975)*, Nosopsyllus fasciatus, Ceratophyllus styx* (Rothschild *et al.* 1975) and in *Tunga* *trimamillata (*Pampiglione *et al.* 2005). That Resilin is present also in thoracic structures, which lack a true pleural arch, was shown for *Isochopsyllus octactenus* and *Spilopsyllus cuniculi* (Rothschild *et al.* 1975). Later Burrows cited Resilin in the thoracic jump mechanism for froghoppers (Burrows 2003) or *Sipyloidea sp*. (Burrows and Morris 2002). Visualisation with fluorescence microscopy and antibody staining in the pleural arch was done in *Ctenocephalides felis* (Lyon *et al.* 2011). For Hemiptera, Rothschild and colleagues showed Resilin with staining in *Javesella dubia* (Rothschild *et al.* 1975); its presence was reported but not shown in *Cercopis vulnerata* (Gorb 2004). Later, by fluorescence microscopy, Resilin was also identified in the pleural arch of *Philaenus spumarius,* *Aphrophora alni* (Burrows 2007, Burrows *et al.* 2008), *Issus coleoptratus* (Burrows 2010), Delphacodes *sp.* (Burrows *et al.* 2011), *Archaeopsyllus erinacei* (Sutton and Burrows 2011), *Metcalfa pruinosa* (Burrows *et al.* 2014), *Thionia bullata* and *Lepyronia quadrangularis* (Siwanowicz and Burrows 2017). In *Boreus hyemalis* (Burrows 2011) and *Micromus variegatus* (Burrows and Dorosenko 2014a), Resilin was shown in the thorax, where it is needed for jumping. In Collembola, Resilin was suggested (Andersen and Weis-Fogh 1964) and later shown by fluorescence microscopy, in the furca, a structure specific for the jump mechanism (Nickerl *et al.*, 2014; Oliveira 2022).

The valves of the pygidial defence system in the bombardier beetles *Brachinus elongatulus* (Arndt *et al.* 2015) or *Brachinus sclopeta* (Giulio *et. al.* 2015) contain Resilin; in pygidial glands, a spraying defence system of *Harpalus pensylvanicus*, Resilin was detected (Rork *et al.* 2019).
In *Orgilus Lepidus*, Resilin was noted in the sting apparatus (Hawke *e**t* *a**l.* 1974) ; or in poision apparatus of *Pogonomyrmex badius* (Varman 1982). With staining and fluorescence, Varman (1981) showed the presence of Resilin in the integument of the abdomen for physogastry in *Odontotermes obesus*. Elvin (*et al.* 2005) compared the Resilin containing tendon of *Zyxomma* *sp*. with a structure formed by recombinant Resilin. With confocal microscopy, the content of Resilin in the neck of *Libellula depressa* was shown (Michels and Gorb 2012). In the fan of larval tail of *Chaoborus crystallinus* (Diptera) (Burrows and Dorosenko 2014b) or in the hair of *Malacosoma castrensis* larvae (Lepidoptera) (Kovalev *e**t* *a**l**.* 2020), Resilin was demonstrated by UV microscopy. More recent studies showed the presence of Resilin in spermathecal ducts, the hair bases and tracheal endings of *D. melanogaster* (Lerch *et al.* 2020). In *Bactrocera oleae* (Rebora *et al.* 2021b) and *Aedes albopictus* (Croce *et al.*2023), Resilin was reported in the tracheal system.

In the present work, we demonstrated Resilin in the inner organs of *T. molitor* and *T. castaneum* (see sfigures s14 and s15) confirming the work published by Happ and Happ (1975).

### Supplementary discussion

#### DT signal versus DAPI signal

For robust identification of Resilin, we adhere to the definition of auto-fluorescence (natural fluorescence of the sample) and background (any detectable and interfering light of sample) of (Reichman 2000). We designed a method avoiding sample (section, buffer) and image manipulation (adjusting gain or saturation) by excluding the background and other auto-fluorescence by a broad DAPI filter differentiated by the DT-specific filter (MISM).

Trivially, using the DT-specific filter for detection improves the sensitivity. Obviously, this facilitates quantification. Thus, in an optimal microscopic setup, intense DT regions are easily detectable allowing unambiguous identification and localisation. The problem arises for regions with weak DT signal. Generally, localisation of especially weak DT areas by fluorescence microscopy may yield false positive or false negative results. Raising the exposure time would worsen image quality (see below). Likewise, the use of broad-range filters like the DAPI-specific filter to enhance signal detection would deteriorate the ratio of true and background signals. Indeed, commonly used DAPI filters have a detection maximum at or beyond 420nm excluding the DT emission maximum of 415nm. Our DAPI filter, for instance, collects light in the range of 420 to 470nm. To gain data on Resilin localization in *P. americana*, Neff (2000) used a filter with a detection lower limit of 420nm, while Ma’s (2015) filter for DT localisation in honeybees detected signals beyond 425nm. We have to be aware of cuticle elements, which emit auto-fluorescence in the settings of the DAPI filter but not of the DT filter. This was nicely shown for the envelope of *D. malonagaster* that does not emit light around 415nm, but around 470nm (Zuber *et al.* 2018 figure 4f). To illustrate the problem with our samples, we compared the data of the adhesive pads in *S. gregaria* and *B. dubia*. In the locust, the adhesive pad signal was visible without any further manipulation; observations with a DAPI filter showed the same results. This is expected for strong signals. By contrast, the cockroach adhesive pads showed a distinct signal detected with the DAPI filter, while the DT-specific filter did not reveal any DT in the adhesive pads. Thus, the DAPI generated signal represents a false positive localization; as shown with the DT filter, the DT signal resides inside the structure. Moreover, the common DAPI specifications would also suggest that large parts of the leg in *S. gregaria* contain DT, consistent with the supplementary data of Burrows and Sutton (Burrows & Sutton 2012 see supplementary data movie 3) showing a bluish leg. Using the DT-specific filter that catches the DT maximum wavelength of 415nm, the leg of the locust is free of any background DT signal. False negative signals may also be present in Hemiptera samples, in *Ephestia kuehniella* or in Coleoptera (see *P. marginta* cut and not cut specimens). These signals were conceivably caused by interference of the cuticle itself (see the dimming problem). To exclude false positive measurements, we established thresholds to assign DT/Resilin in an image.

#### *The threshold*

The thresholds to accept or refute a signal were set at 500 DT-DAPI and 1000 DT-DAPI counts difference as the higher limit. This setting relies on data used from the trochanter intensity mean of 70 individuals from D. melanogaster and D. suzukii (unpublished). The threshold of 1000 DT-DAPI counts difference was chosen based on the DT signal detected in the distal edge of the fly trochanter overlapping with a strong Resilin-GFP signal (Lerch et al. 2020). We noticed, however, that this threshold was too high to assign DT incidences to Resilin-GFP signals in the wing hinge of the fly, therefore, we defined a second threshold with 500 DT-DAPI counts difference to include the detection of rather weak signals. We argue that this adjustment may allow detection of DT signals in species where a Resilin-GFP signal reference is not present.

The higher threshold was sufficient to recapitulate DT signals of reference specimens including the S. gregaria jumpleg triangle (figure 6, sfigure s4 and s5), the P. marginata wing blade signal (sfig. s34 and s35), the D. melanogaster trochanter (fig. 5 and sfig. s1) and the A. melifera wing blade (sfig. s30 and s31). Although the two thresholds were useful in identifying DT signals in many different species, in some cases the detection was still ambiguous. Augmenting specimen number as in the case of *T. castaneum* and *E. kuehniella* was needed to clarify the situation. Overall, the thresholds defined in *D. melanogaster* are useful to identify and quantify DT incidences in many insect species.

The positive of the thresholds to identify and qualify Resilin localisations is shown and is leading to more objective analyse of membranous areas and DT containing cuticles, which lead in this paper to expand knowledge of cuticle composition in a well known model organism (see Results Diptera and discussion non-Resilin DT signal).

#### The exposure time

During this study, we examined the influence of exposure time that depends on the magnification - on the quality of the signal. These two parameters depend, of course, on the size and colour of the specimen. For bigger insects, 5x magnification was necessary. The exposure time varied form 2.3 seconds for brighter insects to often 5.3 seconds for darker ones. For unambiguous signals, as expected, the exposure time did not play a decisive role. In S. gregaria, P. maginata and the Tenebroids, for instance, areas with a strong signal, for example within wing hinges, did not change with exposure time. Likewise, the jump leg triangle in S. gregaria gave a DT-DAPI threshold signal of over 1000 with both exposure times. The influence of the exposure time was obvious in areas with DT-DAPI signals with a value of less than 1000. This point is exemplified in the foreleg of *Pyrrhocoris sp.* and the fore- and hindleg of *S. gregaria* shown in sfigure s40. In the tarsi of *Pyrrhocoris sp.*, low exposure time reveals one reliable signal, while at higher exposure time an additional reliable signal appears. In the trochanter, a signal is detected only by higher exposure times, while the signals in the coxa remain reliable at lower and higher exposure times. In *S. gregaria*, the problem of exposure time becomes critical as in this species the background fluorescent signal of the cuticle is very high causing often an over-saturation of the signal at higher exposure times. A signal appears, nevertheless, when the femur is over-exposed (what would usually not be done during standard microscopy). In the trochanter of one sample, the DT-DAPI signal intensity difference diminishes just to a point of acceptance when over-exposed; in another sample this actually happens, and the presence of DT is refuted. In conclusion, trivially, exposure time that depends on magnification and the tissue tanning defines signal intensity. In the same magnification, there is no linear correlation of exposure time and intensity. Based on this conclusion, we recommend testing single species for magnification and exposure time to minimize oversaturation against the visibility of the structure or use, like for the filter set up, two different exposure times to validate a threshold for the exposure times.

To minimise subjectivity, we used the profile function of ZenBlue (Zeiss), which could be more objectively done by a software scanning the image instead of a human being.

#### The dimming problem

Another problem with fluorescence microscopy is the effect of signal reduction or loss on the specimen, what we refer to as dimming. Light could be lost for different reasons: it may be optically blocked or reflected. Dimming included also the term quenching, which is used in fluorescence microscopy when other molecules in the specimen have the ability to reduce the intensity of the emitted light and thereby mask fluorescence signal (Losi *et al.*, 1993; Reichman, 2000). The insect cuticle is composed of chitin, proteins, lipids and catecholamines that form melanin. Melanin absorbs UV light of a broad range. This composition also defines physical properties including UV protection. The cuticle tanning effect potentially distorts the DT to DAPI ratio. For the localisation of DT signals in different types of tissues, we assume that the presence of quenching molecules influences detection. Inconsistently, our sample of black ants show signals also with a dark or black cuticle in the leg. There consistently, the tanned cuticle of *P. marginata* leg did not show any signal; only by opening the leg structures, we found signals inside the joints. This is consistent with published data. Nadein and Betz (Nadein & Betz 2016, 2018), for instance, had to open the leg of *Sphaeroderma testaceum, Podagrica fuscicornis* and *Orchestes fagi*  to identify a signal. Other instances where DT in leg structures was only detected upon dissection were: the femur-tibia joint of *Lethrus apterus* and *Pentodon idiota* (Frantsevich *et al.* 2019), *P. marginta* (Busshardt *et al.* 2012) or the tenebrionid beetles (Ichikawa *et al*. 2016); inside of tibia of *Carausius morosus* (Schmitt *et al.* 2018), the tarsi and adhesive pads of *P. americana* (Neff *et al.* 2000; Frazier *et al.* 1999), the tarsal pads of *Locusta migratoria* and *Tettigonia viridissima* (Perez Goodwyn *et al.* 2006), the jump apparatus of fleas (Bennet-Clark and Lucey 1967; Rothshild 1975; Rotschild and Schlein 1975; Rothschild *et al.* 1975) and froghoppers (for example: Rothschild *et al*. 1975; Burrows 2007, Burrows *et al.* 2008; Burrows 2010; Burrows *et al.* 2011; Sutton and Burrows 2011; Burrows *et al*. 2014; Siwanowicz and Burrows 2017). Not to mention inner organs like the pygidal glands of bombardier beetles (Arndt *et al.* 2015; Giulio *et al.* 2015).

Burrows and Sutton (Burrows & Sutton 2012) described that the DT signal in the hindleg of locusts at different developmental stages faded from the outside layer because of the darkening of the cuticle. In Mantodea samples, the authors described different degrees of auto-fluorescence that they attributed to sclerotisation and melanisation suggesting their influence on imaging (Roze *et al.* 2024). In the Odonata *Anax imperator*, quenching of the Resilin signal was shown to be related to the mechanical importance of cuticle darkening (Preuss *et al.* 2024). We identified a DT signal in a region with dark cuticle in *E. kuehniella*, that deserves a closer examination with regard to the quenching problem.

Besides of melanin, other molecules such as lipids and proteins could also affect the quenching properties of the cuticle. These is maybe the case for the adhesive pads of *B. dubia*, which revealed Resilin after opening and not being tanned. In comparison in *S. gregaria* is different where the same located adhesive pads show DT without opening.

Thus, in conclusion, detection of DT should take into account that cuticle tanning may influence intensity. Therefore, we suggest, by this kind of cuticles to open the leg and to study further.

#### Comparison of our method with other methods

The usage of other fluorescence microscopy methods like confocal microscopy (Michels and Gorb 2012), two-photon microscopy (Chien et al., 2011; Rabasovic 2015; Mouchet et al. 2023) or even atomic force microscopy (Qin et al., 2009; Dutta et al., 2009; Wanasingha et al., 2025) to study structural elasticity yield a detailed and rich set of data. These methods require expensive hardware and special training and are not common tools in most laboratories. The usage of a simple filter set up is comparably easier especially for bigger samples. We recommend for broad overviews and high-throughput studies of diverse species also the use of standard fluorescence microscopes. Additionally, by our experiences, data acquisition with confocal or two-photon microscope minimizes but does not exclude the background problem with the DAPI filter set up. Therefore, the MISM method might also allow a more precise and objective measurement to identify DT and Resilin or Resilin-like proteins in the cuticle by more sophisticated microscopical methods.

For localisation resilin in living tissues, different classical histological methods have been employed. Abundant resilin has been detected applying histological staining of proteins with dyes like tolidine and methylene blue that mark negatively charged molecules (Weis-Fogh 1961a). Specific antibodies (Burrows *e**t* *a**l**.* 2011; Wong *e**t* *a**l.* 2012) or the expression of tagged versions of Pro-Resilin (Lerch *e**t* *a**l**.* 2020) were also successfully used in the field. Weis-Fogh and Andersen (Andersen & Weis-Fogh 1964) also showed that DT is sensitive to pH-value changes, very often this parameter is used to identify resilin (Gorb 2001; Burrows 2012). Techniques like HPLC (high pressure liquid chromatography) have also been used to detect DT and TT amounts in different body parts of *S. gregaria* (Andersen 1964).

Together, these methods show the presence of DT or resilin in a cuticle structure by, however, often interfering with sample integrity. Dyes like methylene blue or toluidine blue for histological staining are not specific enough to only highlight resilin (Rice 1970; Hermann & Willer 1986). The molecular methods require changes of pH-values or dehydrations stages and sectioning. Sample extraction is a prerequisite for HPLC analyses. Moreover, we believe that these methods are excellent for the identification of cuticle structures with abundant DT and Resilin, but may fail to identify regions with low DT and resilin content. Using two filters with overlapping spectra to eliminate background information allows a sensitive and critical identification of DT and resilin and the possibility to observe the DT signal *in vivo* and *in situ*.

*Wing and head areas of Resilin*

Via MISM, we show the presence of Resilin in all orders in various locations plus validating existing data. A critical and clear comparison with published data is laborious because of the variable set ups additionally to the morphological and structural diversities of the samples. This diversity possibly influences the mechanistic usage of Resilin and thereby directly the variation of detectable DT and leads to major differences even in locations with high preferences for Resilin presence. The most prominent one of these is the flight system. In our study we successfully present references like the wing blade areas in *P. marginata* and *A. mellifera* in the wing blade (like Haas *et al.*, 2000 a,b and Ma *et al*, 2015) with counts over 1000. In other species such as *Tribolium castaneum* (data not shown) and different *Drosophila* species (with the exception of *D. hydei* (Lerch *et al*., 2020)), we were unable to detect DT signals in the wing blade. Wing blade DT signals have been additionally detected in *Calliphora vicina* (Lehmann, 2011; Appel, Michel & Gorb 2023) and Dermaptera (Haas *et al.* 2000b). The common area in our study is the wing hinge, the highly stressed attachment side between body and wing.

The amount and seize of Resilin dots in the wing hinges vary highly. In *D. melanogaster* and *D. suzukii*, we find between 6-7 dots, and two areas in the haltere (Lerch *et al*., 2020; 2022). In *Ephestia kuehnelli* we found around 5 dots, in the elytra and the hindwing of *T. molitor*, there were 6 and 8 dots, respectively. We found one or two dots in the wing hinge of *Apis mellifera*; these could be the same areas which were named by Nachtigall *et al.* (Nachtigall *et al,* 1998). The wing is a complex puzzle of sclerites and membranous parts, powered by muscles. Even small locations could provide mechanical function. We cannot exclude to miss some smaller Resilin/DT areas with our method. In summary, we did not find any general distribution and shape of the wing hinge dots. Only one dot (triangular shaped) in *D. melanogaster*, *D. suzukii* and *D. hydei* seems to be conserved suggesting common biomechanical function. This underlines differences in Resilin presence within a genus. Based on these observations, we imply that lesser amounts of dots is compensated by dot size, and that depending on the wing movement and mechanics the amount of dots changes. It follows that insects with lesser wing blade area, smaller or lesser wings, need more Resilin in smaller dots. Hence, Diptera, who have one wing pair and one pair of halteres, Coleoptera with the elytra and one pair of wings, Hemiptera with often a kind of elytra structure and a pair of wings, these should have smaller and more DT areas in the wing hinge than Orthoptera and Blattodea, who have a wing pair and the tegmen. This remains to be studied in detail. Another important factor depending on the wing morphology to influence the DT distribution of the wing hinge could be the flapping behavior, which differ between species and order (Weis-Fogh, 1972; Sudo *et al.*, 2005 or Wootton, 2020). Similar could be true for the shapes of Resilin in these locations.

As shown in *D. melanogaster* and *D. suzukii*, the dots of the wing hinge are necessary for holding the wings in the resting position (Lerch *et al.*, 2020; 2022), similar to the usage in the wing blade for the folding mechanisms in Dermaptera (Haas, Gorb and Wootton, 2000; Appel, Michels and Gorb 2023) or Coleoptera (Haas, Gorb and Blickhan, 2000; Appel, Michels and Gorb 2023). For a complete understanding of the relation of forces and composition in this location, we suggest a detailed mapping of the different wing hinges, following the example of Appel, Michels and Gorb who presented recently the importance of Resilin in the flight system, highlighting the diversity of morphological use of Resilin with respect to this kind of mechanical stress (Appel, Michels and Gorb, 2023).

The mouth parts of insects that also contain DT undergo diverse morphological adaptations to stress. We showed for some, like Lepidoptera (Hepburn 1971) or in general for Diptera or Hemiptera or Hymenoptera (suggest of Rehder 1987 and Groneberg *et al.* 1993; 1997), that this is the case. For Coleoptera, Orthoptera and Blattodea, we presented images of Resilin-network proteins in their feeding mechanisms for the first time. Because of their variations, a general description at this point of study is not reliable. In comparison to the wing hinge, where size and intensity of Resilin areas did not seem to be correlated (see also Lerch *et al,* 2020), in the proboscis this is a different story. In the more detailed studies of *D. melanogaster* the larger proboscis areas had lesser DT concentration, then smaller areas (Lerch *et al.* 2020; 2022). For distinct species and different proboscis morphologies specific correlations may be possible reflecting usage. An exclusion of missed DT areas because of the threshold seems much more unlikely in comparison to the fiddly wing hinge.

Weis-Fogh and Anderson suggested 1964 that Resilin occurs in all kinds of elastic or flexible cuticle structures. The presence in the diversity of cuticle elements of eye lenses, genital systems, jump mechanism and so on seems to follow their observations based on the presented reviewed literature. Still, it remains a structured identification of a variety of insect species of all orders and movable cuticle segments to fully confirm this kind of hypothesis.
